# TRIP12 governs DNA Polymerase β involvement in DNA damage response and repair

**DOI:** 10.1101/2024.04.08.588474

**Authors:** Burcu Inanc, Qingming Fang, Joel F. Andrews, Xuemei Zeng, Jennifer Clark, Jianfeng Li, Nupur B. Dey, Md Ibrahim, Peter Sykora, Zhongxun Yu, Andrea Braganza, Marcel Verheij, Jos Jonkers, Nathan A. Yates, Conchita Vens, Robert W. Sobol

## Abstract

The multitude of DNA lesion types, and the nuclear dynamic context in which they occur, present a challenge for genome integrity maintenance as this requires the engagement of different DNA repair pathways. Specific ‘repair controllers’ that facilitate DNA repair pathway crosstalk between double strand break (DSB) repair and base excision repair (BER), and regulate BER protein trafficking at lesion sites, have yet to be identified. We find that DNA polymerase β (Polβ), crucial for BER, is ubiquitylated in a BER complex-dependent manner by TRIP12, an E3 ligase that partners with UBR5 and restrains DSB repair signaling. Here we find that, TRIP12, but not UBR5, controls cellular levels and chromatin loading of Polβ. Required for Polβ foci formation, TRIP12 regulates Polβ involvement after DNA damage. Notably, excessive TRIP12-mediated shuttling of Polβ affects DSB formation and radiation sensitivity, underscoring its precedence for BER. We conclude that the herein discovered trafficking function at the nexus of DNA repair signaling pathways, towards Polβ-directed BER, optimizes DNA repair pathway choice at complex lesion sites.

**Graphical Abstract:** 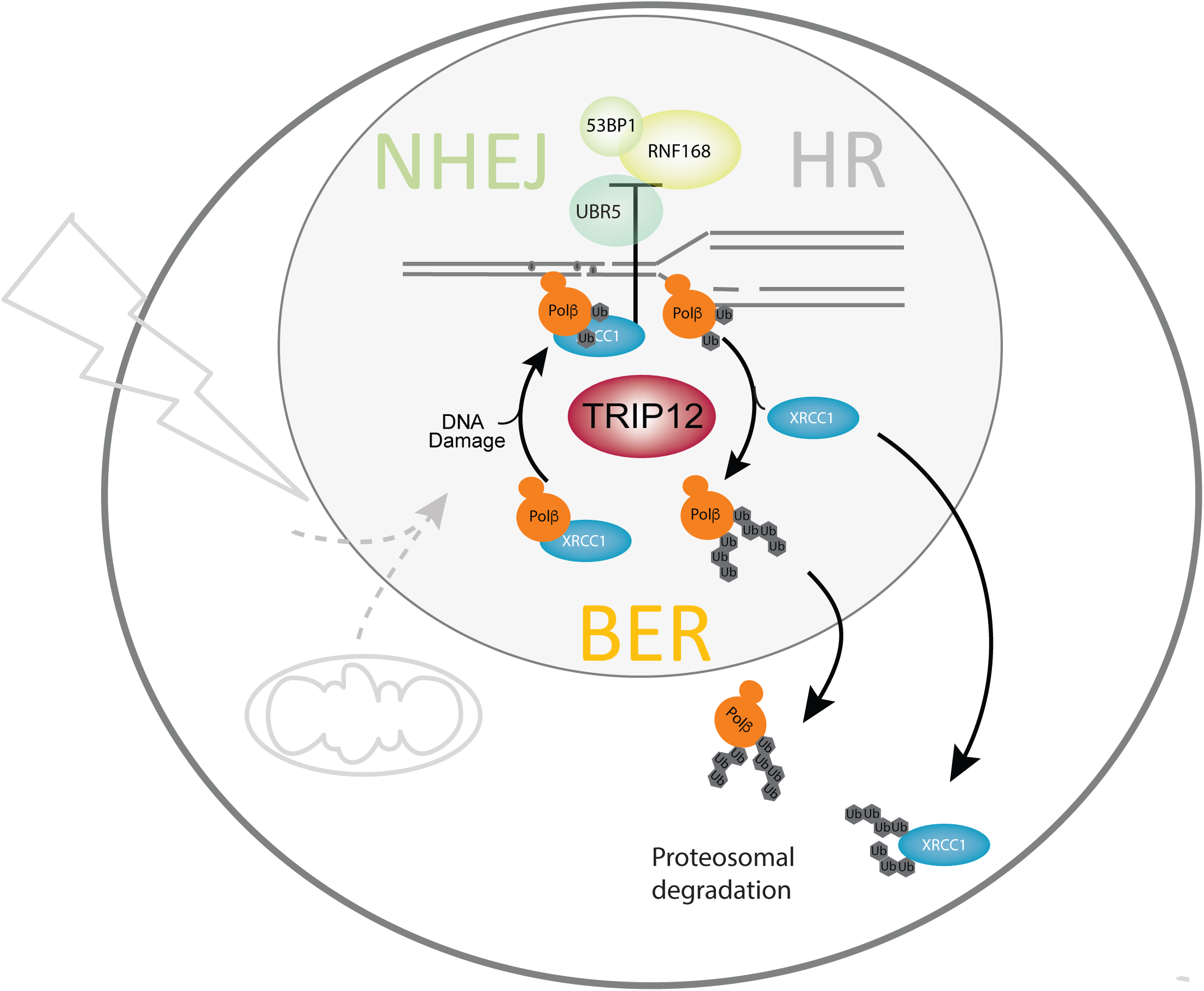

## Introduction

Genome integrity maintenance is challenged by the multitude of different DNA lesion types and the nuclear dynamic context in which they occur. Resolution of DNA damage caused by exogenous or endogenous sources requires the engagement of different DNA repair pathways. This is typically determined by the type of lesion and the cell cycle phase. DNA double-strand break (DSB) repair pathways are tightly regulated to assure efficient repair in different cell cycle phases and chromatin conditions (1,2). Single-strand breaks (SSB) and damaged bases are mainly repaired by DNA polymerase β (Polβ) mediated processes that also consider the cell cycle phase through protein complex composition and stability (3). Replicating DNA or complex DNA lesion sites that contain different lesions in close proximity pose particularly difficult repair conditions as they may require the involvement of intrinsically very distinct repair pathways. Thus, repair pathway choice and sequence are crucial to the success of DNA damage repair under such conditions.

Non-homologous end joining (NHEJ) and homologous recombination (HR) are the most prominent DSB repair options and their interaction and relative contribution has been actively researched in recent years (4,5). DSBs, such as those caused by radiation or generated due to DNA crosslinks from metabolic formaldehyde, cisplatin or similar crosslinkers, can create marked threats to DNA replication and cellular division. These potentially lethal lesions are resolved by DSB repair processes. Base excision repair (BER), on the other hand, deals with base lesions or contributes to SSB repair (SSBR), abundantly generated by metabolic processes, reactive oxygen species (ROS), deamination or depurination (6). Greatly outnumbering DSBs, radiation also causes a multitude of BER-specific lesion types (7,8). Of these, oxidative base lesions and SSBs are the most predominant.

Different DNA lesion types will therefore promulgate repair protein “trafficking” issues when adjacent to each other. In fact, radiation induced complex DSBs, that also comprise multiple base damages, are the most difficult to repair and hence most detrimental to cellular survival and genomic integrity. The chromatin context in which those lesions occur is also an important factor in DNA damage repair that needs to consider replication, transcription, and different (hetero)-chromatic conditions. Even though close links between distinct lesion repair pathways would be expected, no studies exist that demonstrate an inter-pathway collaboration of BER and DSB repair and few report the involvement of individual BER elements in DSB repair processes (9–11). Crosstalk mechanisms that enable DNA repair pathway choice at lesions and chromatin sites that are common to both, and the orchestration of protein complex formation thereof, remains elusive.

Chromatin and repair enzyme modifications by ubiquitylation are relevant processes that regulate the DNA damage response (DDR) and DSB repair (12). An RNF8/RNF168 E3 ubiquitin ligase-coordinated cascade of non-proteolytic ubiquitylation events results in chromatin marks in the vicinity of DSBs. RNF168-dependent chromatin ubiquitylation mediates the accrual of the DSB repair proteins 53BP1 and BRCA1 to the lesion site (13) and additional studies have shown how RNF168-mediated 53BP1 engagement can support the regulation of DSB repair (14–16). 53BP1 recruitment blocks resection, an important HR engaging step, to ultimately promote NHEJ. Resection and therefore HR, on the other hand, is facilitated by BRCA1 by counteracting 53BP1 recruitment at such ubiquitylated sites (17). A recent report further supports a role for RNF168 in facilitating HR through PALB2 loading in a BRCA1-independent manner (18,19). Together, this however highlights the intricate molecular mechanisms by which chromatin ubiquitylation directs DSB repair pathway choice. Surprisingly, to date no similar processes have been linked directly to BER/SSBR.

A different E3 ligase, TRIP12 (ULF), has also been shown to regulate DDR protein homeostasis. TRIP12 prevents 53BP1 hyper-accumulation by controlling RNF168 residence at break sites (20) thus indicating an involvement of TRIP12 in DSB repair suppression. TRIP12 was also reported to affect PARP-inhibitor efficacy (21,22) and to interact with Ku70 (23), while its expression was found to be regulated by p16 (24,25). Despite this and a reported effect on the USP7-regulated stabilization of p53 (26), little is known about TRIP12’s overall impact on DNA repair and cellular functions.

Polβ has a crucial function in BER and SSBR. Upon lesion recognition and removal by glycosylases and AP-endonucleases, Polβ executes end-tailoring and DNA synthesis at the gap (27). The 5’dRP lyase and nucleotidyl transferase activities allow Polβ, supported by XRCC1, to execute nick and single-strand break repair in many cases (28). The very distinctive nature of Polβ’s tissue and developmental expression pattern points to processes that tightly regulate Polβ levels to assure context-appropriate activities (29). Deregulated expression has indeed been reported to lead to interference in replication processes and to defects in Polβ sub-cellular localization resulting in altered damage response (30–35). Such findings led us to postulate the existence of molecular and cellular mechanisms that counteract the dangers of such “over”-repair by stringently controlling Polβ levels to exclude Polβ from such disruptive activities. Our past studies revealed elements in the governance of Polβ levels and stability that ultimately also define BER complex make-up. We elucidated the regulation of the BER/SSBR complex architecture through proteolytic ubiquitylation events and HSP90 binding via its interaction with XRCC1 in response to damage (3). In this earlier study (3), we showed that separation-of-function mutants of Polβ failed to interact with XRCC1 and were targeted for ubiquitylation and degradation. A knock-in mouse model expressing such a separation-of-function mutant of Polβ confirmed reduced Polβ protein levels *in vivo* (36). Importantly, we also found that the proliferation status of the cell and the nature of the damaging agent defined BER complex formation and Polβ stability (3). Solely in proliferating cells and distinctively different from the BER complex formation pattern observed after alkylating agents, radiation exposure led to increased levels of the Hsp90 bound XRCC1 (3). These data led us to hypothesize differential and context dependent BER/SSBR complex formation and pathway engagement during replication and at complex lesion sites. Two select ubiquitylation sites on Polβ suggested an unidentified regulatory component of BER/SSBR. We therefore searched for a ubiquitin ligase that fulfills this role in Polβ regulation and discovered how this new element in BER/SSBR (TRIP12) provides a ‘repair-controller’ function.

## Materials and Methods

### Materials

All materials and supplies are listed in **Supplementary Table S1**.

### Cellular models

Cell line models were developed by lentiviral expression of the indicated proteins (Flag-Polβ, myc-TRIP12, copGFP-Polβ) as indicated in detail in the **Supplementary Document S1** and as listed in **Supplementary Table S1**. In some cases, modified cells were modified by a second transduction, such as expressing TRIP12-shRNA in cells modified for expression of Flag-Polβ, and using lentiviral vectors with different selection makers (Puromycin, Geneticin, Hygromycin). Flag-Polβ, myc-TRIP12, myc-TRIP12-HECT and myc-TRIP12-SB vectors were mainly used to assess binding interactions or ubiquitylation assays, as indicated throughout. copGFP-Polβ based models were generated for laser-induced micro-irradiation experiments, radiation or cisplatin induced foci analysis and nuclear co-localization analyses. For over-expression models, cell lines were developed with elevated Flag-Polβ expression and respective mutants. Several different TRIP12 shRNA-based vectors were used to establish the cellular role of TRIP12 expression in Polβ stability and radiation response.

### Methods

Detailed methods are described in the **Supplementary Document S1**

In brief, standard cell lysis, chromatin fraction isolation, immunostaining and immunoblotting and immunofluorescence microscopy procedures were applied using the antibodies and conditions as indicated in the figures and legends and detailed in **Supplementary Document S1** and **Supplementary Table S1**.

For complex partner identification and immunoprecipitation, anti-Flag M2 affinity gel was used to immunoprecipitate the individual proteins after cell lysis with Pierce IP lysis buffer and according to the manufacturer’s protocols. 27 individual IP samples were prepared (n=9 per condition) and loaded onto SDS-PAGE gels for label-free differential mass spectrometry (dMS) analysis. Tryptic peptides extracted from each IP product were separated by reverse-phased nano-flow liquid-chromatography (EASY-nLC II, Thermo Scientific, San Jose, CA) and analyzed on an LTQ/Orbitrap Velos Elite hybrid mass spectrometer (Thermo-Fisher, San Jose, CA). Mass spectrometry data collected for each sample was analyzed using dMS software (Infoclinika, Bellevue WA). The high-resolution full MS spectra were aligned and the m/z, charge state, retention time and intensity data for all molecular features detected in the full scan mass spectra were integrated and matched to protein identification results as detailed in **Supplementary Document S1**. Student’s t-test implemented in MATLAB® was used to determine the statistical significance of the difference on the abundance of identified proteins/features in different IP samples. The abundance values in Flag-Polβ(WT) and Flag-Polβ(TM) samples were normalized to the Polβ level in each sample to calculate the WT/TM ratio used to assess the differential interaction between Flag-Polβ(WT) and Flag-Polβ(TM) samples.

Recombinant proteins (His-tagged ubiquitin (His-Ub) were generated as detailed in **Supplementary Table S1** and **Supplementary Document S1**. To study interactions with TRIP12 or ubiquitylation of Polβ, immunoprecipitations were performed using TRIP12, Myc and anti-Polβ clone 61 antibodies (**Supplementary Table 1**) as indicated in the figures and following standard protocols (**Supplementary Document S1**). To assess ubiquitylation activity of TRIP12 and the HECT or substrate binding (SB) domains of myc-TRIP12, myc-HECT, myc-HECT(C2007A) and myc-TRIP12-SB were immunoprecipitated using an anti-myc antibody or Myc-Trap agarose and exposed to *in vitro* ubiquitylation assays (**Supplementary Document S1**) with or without purified Polβ. Ubiquitylation of purified Polβ was examined after gel-electrophoresis and immunoblotting with anti-Polβ Ab (Clone61), anti-ubiquitin and anti-His-tag antibodies. Auto-ubiquitylation of the TRIP12 HECT domain was examined after gel-electrophoresis and immunoblotting as described above with an anti-ubiquitin or anti-His-tag antibody (**Supplementary Document S1**).

For Polβ stability and degradation evaluation, cells were seeded and treated with 0.2mM Cycloheximide (Cyc) or with Cyc (0.2mM) and MG132 (25μM) for the time periods as indicated in the figures.

Radiation sensitivity and H_2_O_2_ induced cytotoxicity were determined using the colony formation assay as described in **Supplementary Document S1**. The CometChip assay was used to assess DNA damage and repair kinetics and the DNA fiber assay was used to evaluate the reported replication parameters as outlined in **Supplementary Document S1**.

### Statistical analysis

The statistical analysis procedures and parameters for the different analyses are described in **Supplementary Document S1**.

In brief, averages, and standard deviations (SD) were calculated from the means (on technical replicates) of multiple independent experiments (n = number of independent experiments as indicated in figure legends) unless stated otherwise. ANOVA was used to test for significant differences, generally compared to controls and as indicated in the figure legends. Foci dot plots show the distribution of the foci number per cell values and contain pooled foci data from all independent experiments, with a minimum of 50 analyzed cells for each experiment. A non-parametric test (Kruskal-Wallis with Dunn’s multiple comparison test) was used if the data did not pass the normality test. A second analysis considers the inter-experimental variation and compares the means of the individual experiments by calculating the average and standard deviation (SD) of the mean foci number per cell values derived from the independent experiments (**Supplementary Document S1**).

As dependence of the individual data points in the survival dose response data precludes ANOVA analyses, non-linear regressions were computed using Y=100/(1+(X^HillSlope)/(IC50^HillSlope)) (with Y= Survival in % and X=H_2_O_2_ concentration) on the normalized H_2_O_2_ dose response survival data to calculate IC_50_ (half maximal inhibitory concentration) values for each individual experiment. Similarly, D_37_ (radiation dose permitting 37% cell survival) values were calculated from linear quadratic fits (Y=exp(-α*X-β*X^2) with X=radiation dose and Y=surviving fraction) on the survival curves. One-way ANOVA was then used to test for significant differences in these response parameter values as stated in the text. P-values are indicated by asterisks with *p<0.05, **p<0.01, ***p<0.001, ****p<0.0001.

## Results

### DDR suppressor TRIP12 is a novel partner of the BER protein Polβ

The majority of Polβ partners with XRCC1 to conduct BER (37). Yet, XRCC1 and Polβ function independently, in different BER sub-complexes, for yet undefined roles in DNA metabolism (3). We recapitulated these main cellular BER complexes (herein termed Complex A, comprised of Polβ and XRCC1 and Complex B, XRCC1-devoid complexes) by exploiting a separation-of-function mutant of Polβ that does not bind to XRCC1, the V303-loop mutant Polβ(TM) (**Figure S1A**). This approach allowed us to reveal novel partners of Polβ that are not dependent on XRCC1 (**Figure 1A**). Using label-free differential mass spectrometry (dMS), an unbiased quantitative proteomic method (38,39), we discovered TRIP12, a DSB repair-related E3 ubiquitin ligase, as a novel Polβ partner. We found TRIP12 to bind preferentially to the Polβ-centered Complex B, suggesting involvement in BER complex regulation (**Figures 1B, S1B, S1C** and **Table S2**). Polβ(TM) cellular levels drop significantly (**Figures S1B, 1C)**, as also reported previously (3). Yet, we observed a strong interaction with TRIP12 (**Figures 1B-C**). Other novel partners found include RFC1, EIF, BCKDHA, SPT16 and SSRP1 (**Supplementary Table S2**, **Figure S1D**). Quantitative dMS and immunoprecipitation/immunoblot (IP/IB) analysis confirms the lack of XRCC1 and known XRCC1-mediated BER factors (LIG3, PARP2, PNKP, Aprataxin) bound to Polβ(TM), highlighting Polβ specificity of the approach, and points to the dominance of TRIP12 among non-BER proteins in both the transgenic and endogenous setting and in different cell lines (**Figures 1C-D, S1C-E**).

**Figure 1.**
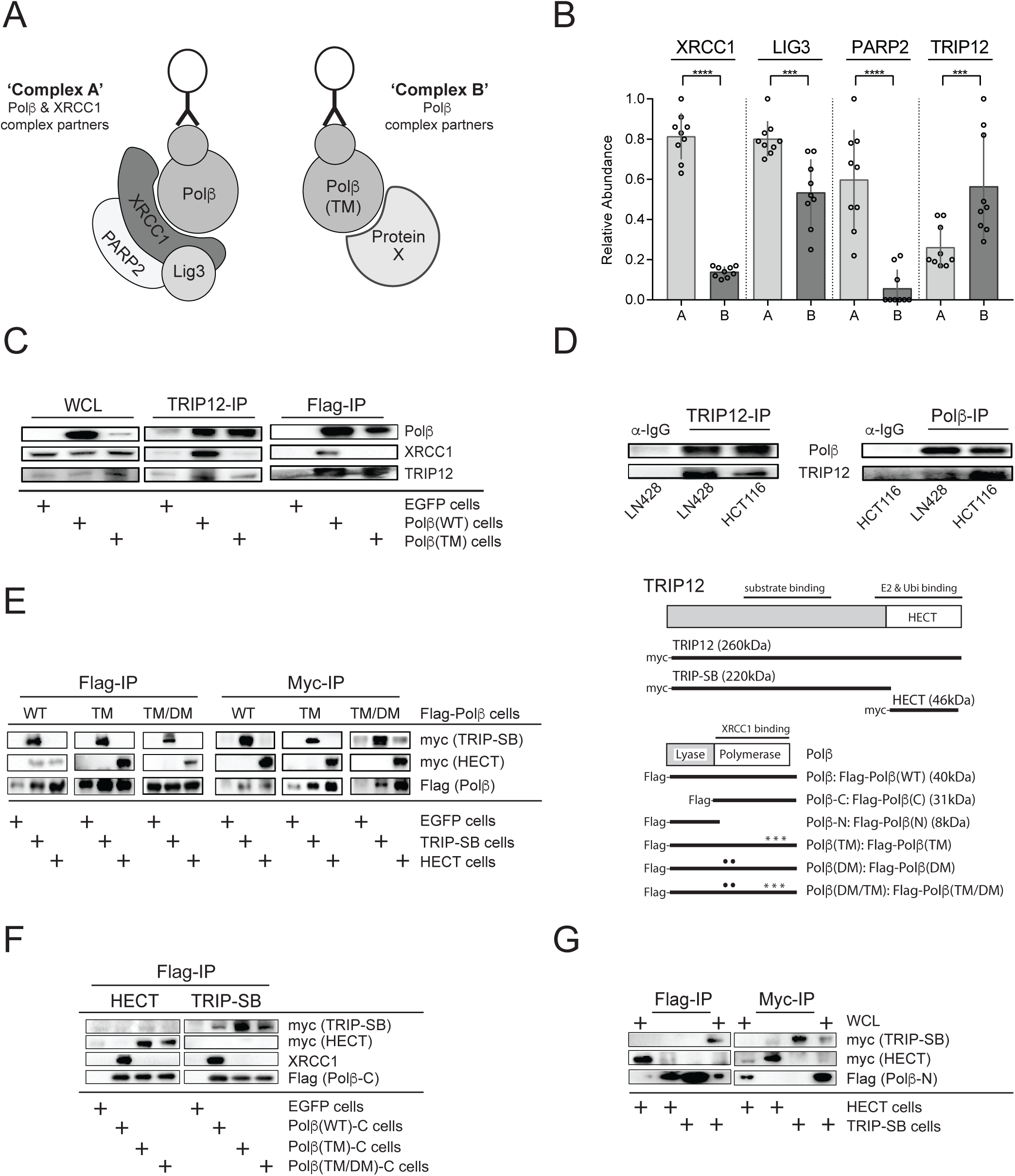
TRIP12 as a novel Polβ partner. **(A)** BER complex compositions and differential partner identification strategy. Cartoon depicts immunoprecipitation scheme used to discover novel interacting BER protein partners. **(B)** TRIP12 is a novel Polβ binding partner with an increased affinity to the XRCC1-free complex (Complex B). Quantification of selected proteins bound to Complex A or Complex B, as indicated, by label-free differential mass spectrometry. The relative abundances of prototypic peptides with unique amino acid sequences AIGSTSKPQESPK, SEAHTADGISIR, VNNGNTAPEDSSPAK and LSTQSNSNNIEPAR were used as surrogate measures of XRCC1, LIG3, PARP2 and TRIP12, respectively. Peptide abundance levels were normalized to Polβ in each immunoprecipitated sample and are shown as dots on bar plots with mean and SD. Significant differences between the complexes A and B are marked by asterisks (***p<0.001, ****p<0.0001; ANOVA). **(C)** BER complex dependent interaction of TRIP12. Interaction of TRIP12 with Flag-Polβ(WT) or Flag-Polβ(TM) as revealed by immunoprecipitation (IP) from whole cell lysates (WCL) and immunoblotting (IB) as indicated. **(D)** Interaction of endogenous Polβ with TRIP12. IP/IB of endogenous proteins in the indicated cell lines are shown. **(E)** Domain interaction mapping. Flag-Polβ binding to myc-TRIP12 domains as depicted in the scheme shown to the right and determined by IP/IB. **(F)** Domain interaction mapping of myc-TRIP12 to the C-terminal domain of Polβ wildtype and mutants by IP/IB. **(G)** Domain interaction mapping of myc-TRIP12 to the N-terminal domain of Polβ wildtype and mutants by IP/IB.

Domain mapping of the interaction between the TRIP12 E3 ligase active site HECT domain or the remaining N-terminal E3 ligase substrate binding domain (TRIP-SB) shows Polβ binding at TRIP12-SB and a preferential binding to the HECT and TRIP-SB domains in the Polβ-centric Complex B as revealed by the interaction with Polβ(TM) (**Figures 1E-G, S1F**). This binding is mediated by the C-terminus of Polβ that also binds to XRCC1 and contains the ubiquitylation target sites (K206, K244; mutated to alanine in Polβ(DM)) (3).

#### TRIP12 ubiquitylates Polβ and targets it for degradation

The interaction of TRIP12 with Polβ and its complex-selective nature suggests a role for TRIP12 as the long-sought Polβ E3 ligase regulating the observed cell cycle and DNA damage type dependent stability of Polβ (3,36). We therefore determined whether TRIP12 ubiquitylates Polβ and whether this TRIP12-mediated ubiquitylation depends on the previously identified lysines (K206/K244) (3). An *in vitro* ‘on-bead’ assay (40–43) was used to show that TRIP12 is capable of ubiquitylating purified recombinant Polβ with evidence of prominent mono and (poly)-ubiquitylated species. This is mediated by the full length TRIP12 or the TRIP12-HECT domain and is specific to the HECT domain active site cysteine (C2007) (**Figures 2A, 2B, S2A**). *In vivo* evaluation of the ubiquitylation of Polβ confirmed TRIP12-dependent (poly)ubiquitylation that is increased in the triple mutant Polβ(TM) which does not bind XRCC1 and forms BER Complex B (**Figure 2C**). TRIP12-dependent (poly)ubiquitylation is abrogated when additionally modifying the potential ubiquitylation target sites in the Polβ(TM/DM) mutant (**Figures 2C, S2B**). Consistent with its proposed function (44), we observe binding of the TRIP12-HECT domain to ubiquitylated Polβ (**Figure S2C**). Loss of XRCC1 binding results in 5-fold decreased basal levels in such Polβ mutants (Polβ(TM)); addition of the K206/K244 mutations (Polβ(TM/DM)) however reverts this instability phenotype (3). Here we found that TRIP12 affects Polβ levels, revealing its role in this Polβ degradation route (**Figure 2D**). Further underlining a complex-specific (and XRCC1-dependent) activity, its loss stabilizes Polβ(TM) most profoundly (**Figures 2D, S2D, S2E**). This effect on Polβ levels is dependent on the potential ubiquitylation target sites K206/K244 (mutated in Polβ(TM/DM)) despite similarly efficient binding to full-length TRIP12 (**Figure S1F**). As shown previously, if bound to XRCC1 (Complex A) in undamaged cells, Polβ is stable and not subject to proteosomal degradation within a 6h timeframe, however, is rapidly degraded if unable to bind to XRCC1 (3) (**Figure S2F**). We further show that this degradation of (XRCC1-free) Polβ is regulated by TRIP12 through proteasome-mediated degradation processes (**Figure S2F, S2G**).

**Figure 2.**
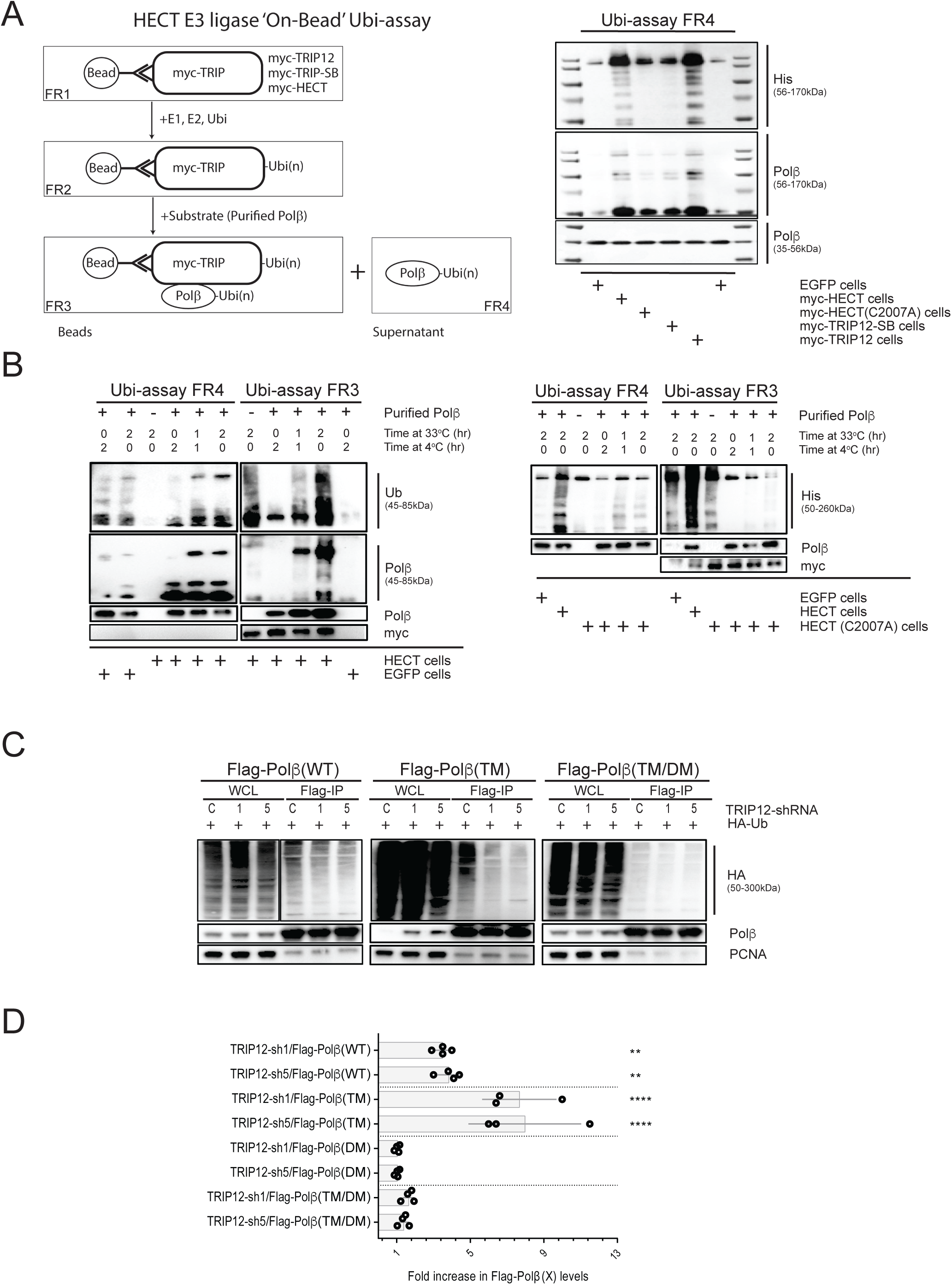
TRIP12’s role in Polβ ubiquitylation and degradation. **(A)** *Ex vivo* Polβ-ubiquitylation by TRIP12. Polβ-ubiquitylation by TRIP12 as identified by immunoblot (right) after the ‘on-bead’ Ubi-assay as outlined in the scheme (left) using recombinant and purified His-Ubiquitin, E1/E2 and Polβ together with myc-HECT, myc-HECT(C2007A), myc-TRIP12-SB or myc-TRIP12 expressed in and isolated from LN428 cells. Boxes indicate the fractions analyzed (FR1-FR4). Immunoblot of FR4 (right) indicates that Polβ is ubiquitylated by full length myc-TRIP12 or the HECT domain but not the Substrate-binding domain or the HECT domain with the active site mutation (C2007A). **(B)** TRIP12-HECT dependent ubiquitylation. Immunoblots as in (A), using FR3 and FR4 from the Ubi-assay as indicated in A with myc-HECT (wild-type or C2007A mutant) after different incubation times. **(C)** TRIP12-dependent ubiquitylation of transgenic Polβ in cells. Ubiquitylation of Flag-tagged wildtype Polβ (Flag-Polβ(WT)), XRCC1-binding mutant Polβ(L301R/V303R/V306R) (Flag-Polβ(TM)) and of the ubiquitylation and XRCC1-binding mutant Polβ(L301R/V303R/V306R/K206A/K244A) (Flag-Polβ(TM/DM)) was determined by IP/IB following transfection of HA-ubiquitin and using two different shRNA (sh1 and 5) to TRIP12. **(D)** TRIP12 affects cellular Polβ levels. Control-shRNA (SCR) normalized Polβ levels in TRIP12-deficient LN428 cells (as in **Figure S2DE**) as determined by multiple immunoblots (representative IB in **Figure S2D**), indicated by dots with SD and n=3-4. Asterisks indicate statistically significant differences (**p<0.01, ****p<0.0001, ANOVA) to the corresponding control-shRNA results.

### TRIP12’s impact on base excision repair processes

While TRIP12 contributes to Polβ degradation in the cytosol (**Figure 2**), it mostly resides in the nucleus (**Figure 3A**) (45,46) where it paradoxically promotes Polβ chromatin association (endogenous Polβ **in Figure 3B**) via the Polβ K206/K244 ubiquitylation sites (transgenic Polβ(DM) in **Figure S3A**). This prompted us to investigate TRIP12’s role in BER beyond an influence on BER complex specific Polβ degradation. At the chromatin level, TRIP12 has been reported to prevent excessive DNA damage signaling and DSB repair protein accumulation to contain the cellular response to DSBs (20). After confirming that TRIP12 depletion affects RNF168 levels also in our cellular models (**Figure S3B**), we first questioned whether TRIP12 affects Polβ’s primary function in BER or affects BER overall. Initial recruitment or retention of Polβ at laser-induced damage sites is not significantly affected by TRIP12 (ANOVA p=0.2 to 0.65; **Figure 3C, S3C**). There is also no evidence that TRIP12 is essential to efficiently execute oxidative or alkylation damage repair (**Figures 3D, 3E, S3D**). Testing a potential involvement in cellular fitness, we find that loss of TRIP12 does not affect proliferation or survival (**Figures 3F, 3G**). TRIP12 appears not essential for canonical BER activities, or its loss is compensated for by other regulators.

**Figure 3.**
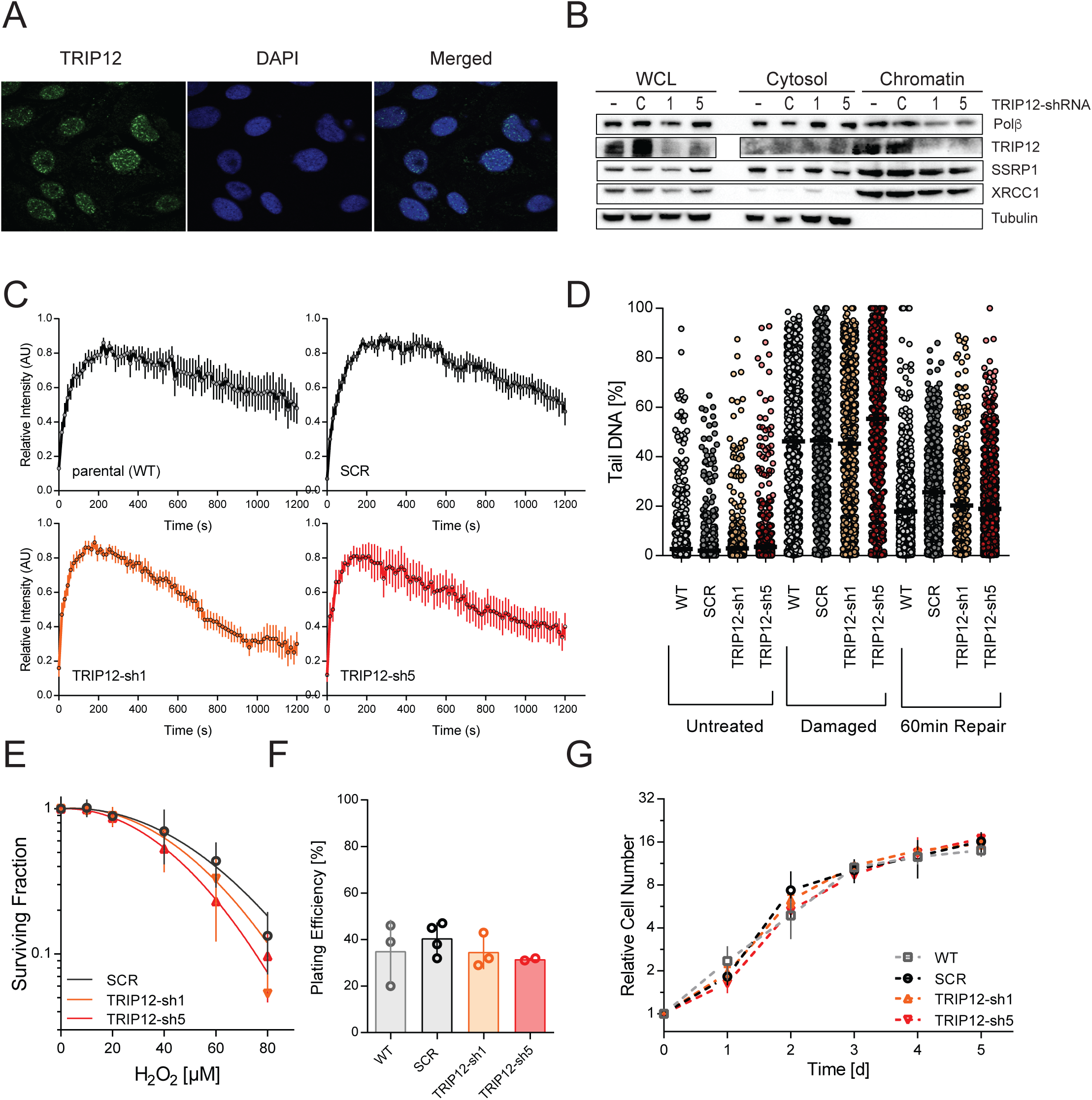
TRIP12-mediated Polβ chromatin retention and BER function. **(A)** Prominent nuclear localization of TRIP12. Immunofluorescence images with TRIP12 antibody. **(B)** TRIP12-mediated Polβ chromatin retention. Immunoblots show Polβ, XRCC1, tubulin (cytosol fraction loading control) and SSRP1 (chromatin fraction loading control) levels in whole cell lysates (WCL), the cytosolic or chromatin fraction in two different TRIP12-KD (knockdown) cell lines (TRIP12-sh1 and -sh5), scrambled controls (C) and the parental LN428 cell line (-) as indicated. **(C)** Laser damage induced focal recruitment of Polβ, independent of TRIP12. Quantification of laser (405nm) induced local recruitment and retention of copGFP fused Polβ (copGFP-Polβ) in TRIP12 knockdown (TRIP12-sh1 and -sh5) and control LN428 cell lines. Data are from n=2 independent experiments with each n=10 cells. **(D)** Effective oxidative damage repair in TRIP12-depleted cells. Oxidative damage and repair as determined by alkaline CometChip analyses (Tail DNA in %) following a 30 min exposure to 250 μM H_2_O_2_ (damaged) and at 60 min post exposure (repair) in TRIP12-KD (TRIP12-sh1 and -sh5), scrambled (SCR) or parental (WT) control cell lines. **(E)** Cellular response to oxidative damage is not affected by the loss of TRIP12. H_2_O_2_ sensitivity of TRIP12-depleted cell lines (TRIP12-sh1 and - sh5) is not significantly different from scrambled controls (SCR) as determined by clonogenic survival and curve fit comparisons or H_2_O_2_ IC_50_ determinations (ANOVA). Shown are the mean surviving fractions of 3 independent experiments +/- SD. **(F)** Depletion of TRIP12 does not affect cellular survival. No significant changes (ANOVA) were observed in the clonogenic survival of TRIP12-depleted cell lines (TRIP12-sh1 and -sh5), scrambled controls (SCR) and parental LN428 cells as determined by colony formation assays. The mean of 3-4 independent experiments, each indicated by dots with +/- SD, are shown. **(G)** Depletion of TRIP12 does not affect cell growth. TRIP12 depleted cell lines (TRIP12-sh1 and -sh5), LN428 parental cells (WT) and scrambled control (SCR) growth as determined by the MTT assay (mean and SD of n=4).

### TRIP12 facilitates Polβ engagement after radiation

Polβ has also been reported to participate in non-canonical BER-like processes in DSB repair. We previously reported that XRCC1-dependent functions of Polβ differ and depend on damage type and proliferation status, thereby linking BER complex make-up to a radiation damage and/or replication context (3). We observed enhanced HSP90/XRCC1 binding after radiation in proliferating cells that consequently promotes the Polβ-centric Complex B (XRCC1-free), herein observed to be preferred by TRIP12 (**Figure 1**) (3). This prompted an assessment of the Polβ response to radiation. Here we find that after exposing cells to radiation, Polβ has both a rapid and persistent focal appearance in a radiation dose (thus damage load) dependent manner that indicates involvement in radiation-induced DNA damage repair (**Figures 4A, S4A-B**). Polβ foci are not associated with radiation induced DSB response and repair markers (γH2AX, Rad51 and 53BP1), thereby largely excluding localization at (chromatin marked) DSBs or a significant participation in DSB repair (**Figure 4B**). We also observed that Polβ is generally barred from ongoing replication since Polβ foci do not form at actively replicating sites and do not show incorporation of large quantities of EDU (**Figure S4C**). Local recruitment of Polβ is accompanied by XRCC1 (**Figure 4B**). Importantly, TRIP12 promotes Polβ accumulation and foci formation, since TRIP12 depletion lowers baseline foci levels in untreated cells and abrogates Polβ foci induction by irradiation (**Figures 4C-E**). TRIP12 depletion affects the formation of large and persistent Polβ foci most profoundly, which appear to be characteristic to radiation exposure (**Figures 4C, S4D-F**). Notably, by 24h and consistent with its role in Polβ degradation, TRIP12 lowered cytoplasmic levels of copGFP-Polβ (**Figures 4A, 4C**), further indicating radiation-induced TRIP12 activation that also facilitates cytoplasmic Polβ degradation as described above (**Figures 2D, S2F**). Cisplatin treatment, in contrast, solely induces small Polβ foci that are however not dependent on TRIP12 (**Figures S4E, S4F**).

**Figure 4.**
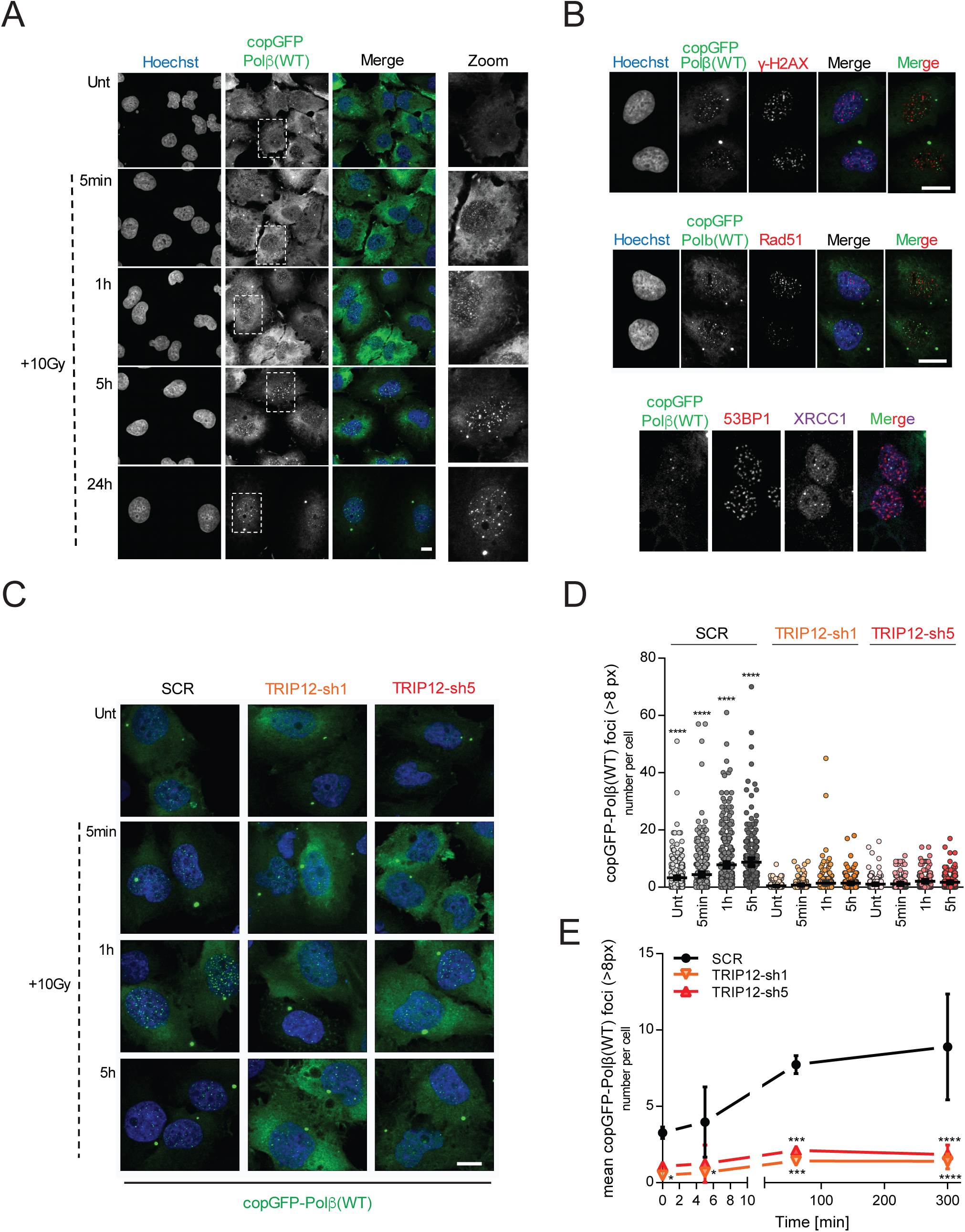
TRIP12 promotes radiation induced Polβ foci formation. **(A)** Polβ foci formation by radiation. Representative images of copGFP-fused Polβ (copGFP-Polβ) foci in LN428 cells at different time points after radiation (10Gy). **(B)** No evidence of co-localization of Polβ (copGFP-Polβ with γH2AX (top), Rad51 (middle) or 53BP1 (bottom) in response to radiation (10Gy, 5 hours). Radiation induced Polβ foci do co-localize with XRCC1 (bottom). **(C)** TRIP12-mediated Polβ foci formation. Representative images of TRIP12-controlled Polβ recruitment after radiation (10Gy) in scrambled (SCR) control cells and TRIP12 depleted (TRIP12-sh1 and -sh5) cells. **(D)** Late and large Polβ foci and Polβ foci rich cells are particularly affected by TRIP12 depletion. Graph shows quantification of radiation-induced Polβ (copGFP-Polβ) foci in scrambled (SCR) control and TRIP12-sh1 and -sh5 cells as indicated. Data show the number of large (>8px) Polβ foci per cell from n=3 independent experiments with >50 analyzed cells each, bars indicate means with SD. Asterisks mark multiple comparison adjusted p-values (****p<0.0001) in the Kruskal Wallis test comparing the different cell line results at each time point to each other. **(E)** Robust abrogation of radiation induced Polβ foci by TRIP12 depletion. Quantification of radiation-induced Polβ (copGFP-Polβ) foci in scrambled (SCR) control and TRIP12-sh1 and -sh5 cells as in (D). Graph demonstrates the inter-experimental variation not visible in (D) and shows the average and SD of the means from n=3 independent experiments over time. Control SCR data used as reference, asterisks indicate significantly different mean foci numbers with *p<0.05, ***p<0.001 and ****p<0.0001 (ANOVA). ANOVA reports a significant interaction with p<0.05. Radiation induced foci are significantly different from untreated, only in the SCR for the 1h and 5h data points with p<0.001 and p<0.0001, respectively.

TRIP12’s role in Polβ recruitment after radiation is surprising when considering its previously reported DDR suppressive function (20) and suggests precedence for BER over DSB repair. The orchestration of repair protein engagement is indeed particularly challenging at complex lesion sites that are typical to radiation exposure (47). Risks due to a lack of repair co-ordination is two-fold: strand incision during repair can cause DSBs at counter-posing BER lesions while base lesions can also hamper DSB repair. Thus, BER and DSB repair elements need to be tightly regulated both locally and in a timely manner. Complex radiation-induced lesions and stalled replication sites with extensive BER lesions may therefore require a ruling in favor of BER (and prior to DSB repair) for optimal repair (48). This led us to propose a DDR trafficking controller role for TRIP12 that ultimately prioritizes BER (through Polβ) over DSB repair in the competition for access to lesions that may activate or require both repair processes.

### Role of Polβ levels in cellular survival and radiation response

Interference in such a postulated DDR trafficking role could evoke a survival detriment. However, despite a large impact on the nuclear engagement of Polβ, TRIP12 depletion did not affect cellular survival (**Figures 3E-G**). Radiation-induced complex lesion sites require highly balanced and coordinated BER/SSBR processes to prevent DSB formation from opposing lesion incisions (49). While impacting Polβ recruitment after radiation, TRIP12 depletion however also alleviates constraints on DSB signaling which in turn facilitates compensatory DSB repair for cellular survival. In line with this, Lukas and colleagues found that depletion of TRIP12 led to an increase in the levels of RNF168 and promotes ubiquitylation near DNA damage sites with little increase in survival after radiation, as compared to controls (20). In line with these earlier findings, we find that survival after radiation is not affected by TRIP12 depletion (**Figure S5A**). We therefore argued that, instead, excessive TRIP12-mediated shuttling of Polβ due to Polβ overabundance at supra-physiological levels (over-expression such as seen in some cancer types, developmental stages and tissues) could expose prioritization of BER and the postulated concomitant suppression of DSB repair signaling. Supporting our proposition, we find that deregulation by Polβ over-expression causes increased residual DSBs after radiation (**Figures 5A-B**). In part, these also result from interference in replication-associated processes or BER that caused stalled replication forks (**Figure 5B, S5B**). The interference in replication is also evident in the impaired HU recovery from stalled replication caused by Polβ excess (**Figure 5C**). Our data further show that the ultimate impact from Polβ over-expression on cellular survival and radiation response is as important as the loss of the SSBR/BER protein XRCC1 (**Figure 5D**). Together, from this we conclude that excess Polβ results in an over-proportional intrusion in lesion repair causing replication fork re-installment problems and secondary DSBs from those attempts or at deregulated clustered lesion sites.

**Figure 5.**
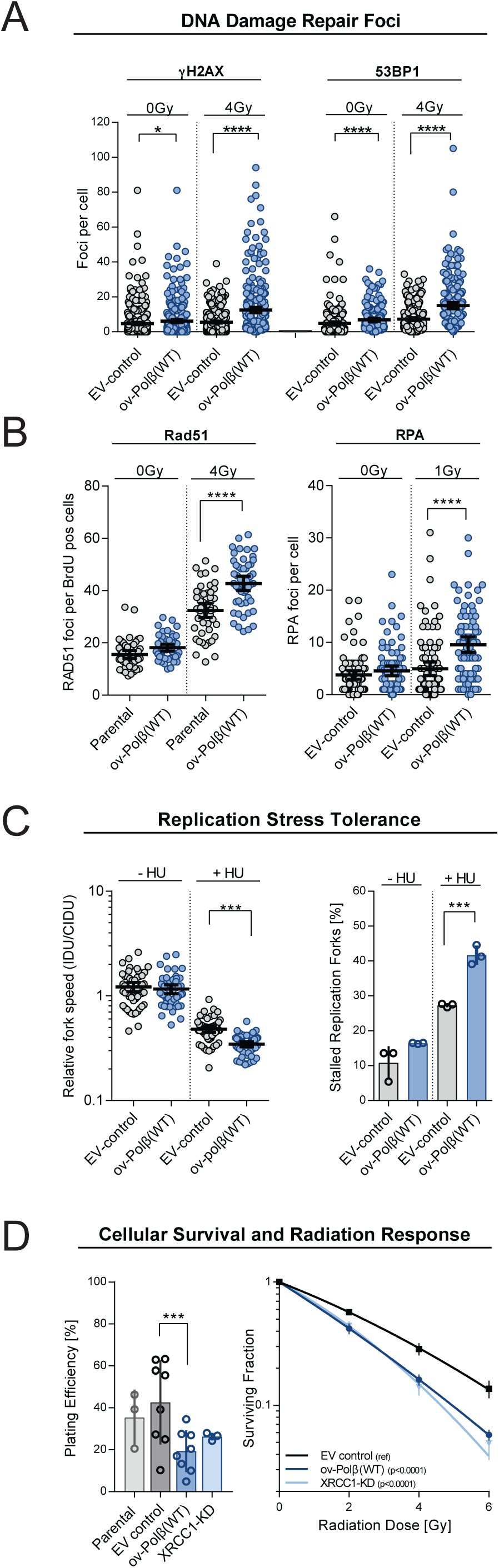
Forced Polβ imbalance and intrusion in replication associated repair. **(A)** Polβ over-expression results in increased residual DSBs. Forced Polβ shuttling by excess Polβ results in increased residual ψH2AX and 53BP1 foci per cell after radiation (4Gy). Polβ over-expressing LN428 cells (ov-Polβ(WT)) were compared to empty vector controls (EV-control). Dot plots show foci per cell counts of n=3 independent experiments with minimal 50 cells each: *p<0.05 and ****p< 0.0001 (ANOVA). **(B)** Replication associated damage due to Polβ imbalance. Increased Rad51 and RPA foci formation after radiation in Polβ over-expressing cells (ov-Polβ(WT)). Graph shows the mean with SD and the individual RAD51 and RPA foci per cell counts in replicating, BrdU positive, cells. Data are from n=3 independent experiments with 50 cells each; ****p<0.0001 (ANOVA, Kruskal Wallis for RPA foci due to normality violation). Not marked is the significant induction of RAD51 foci by radiation in both cell lines with p<0.0001 (ANOVA). **(C)** Recovery from replication arrest after HU treatment is obstructed by excess Polβ shuttling. Fiber assays shows decreased fork speed and an increased fraction of stalled replication forks after 200µM HU treatment in Polβ over-expressing cells (ov-Polβ(WT)) as compared to empty vector controls (EV-control). Left graph shows the mean, SD and individual IDU/CIDU length fractions of n=3 independent experiments. Right graph demonstrates the increase in % stalled replication forks in n=3 independent experiments. Values are mean and SD; ***p<0.001 (ANOVA). **(D)** Radio-sensitization by deregulated Polβ. Cellular survival (clonogenicity as plating efficiency) and survival after radiation drops in Polβ over-expressing cells (ov-Polβ(WT)) to the same extent as in XRCC1-depleted (XRCC1-KD) cells. Bars and values show the mean and SD of the averages from 3-7 independent experiments as indicated by the dots: ***p<0.001 for the plating efficiency data (ANOVA). P values indicated in the radiation response curve graphs assess the likelihood of the data curve fits to be similar. Radiation response parameters (D37%) in the ov-Polβ(WT) and XRCC1-KD differ significantly from the reference (ref) EV-control cell line with p<0.01 and p<0.01 (ANOVA) respectively.

### TRIP12 and its Polβ ubiquitylating activity controls Polβ repair function by chromatin engagement

These findings (**Figure 5**) highlight the relevance of a finely tuned balance between BER and DSB repair protein levels in the maintenance of chromosomal integrity. TRIP12’s Polβ shuttling and ubiquitylation activity, together with its reported DSB repair confinement function, points to TRIP12 as a likely candidate for such a cellular role. Confirming TRIP12’s governance in Polβ shuttling and a ruling in favor for BER, TRIP12 depletion rescued cells from the Polβ over-expression phenotype (**Figure 6A**) that caused increased DSBs and cell death after radiation. Abrogating TRIP12 ubiquitylation sites on Polβ completely suppresses the local Polβ accumulation after radiation (**Figure 6B**). Ubiquitylation by TRIP12 is required for shuttling in favor of Polβ, as shown by the reduction of residual DSBs (**Figures 6C**) that result from it. Finally, abrogation of TRIP12 ubiquitylation also ultimately impedes the survival detriment caused by excess Polβ and further confirms a functional link to TRIP12 (**Figure 6D**). Thus, TRIP12 governs radiation response and controls Polβ engagement through Polβ ubiquitylation.

**Figure 6.**
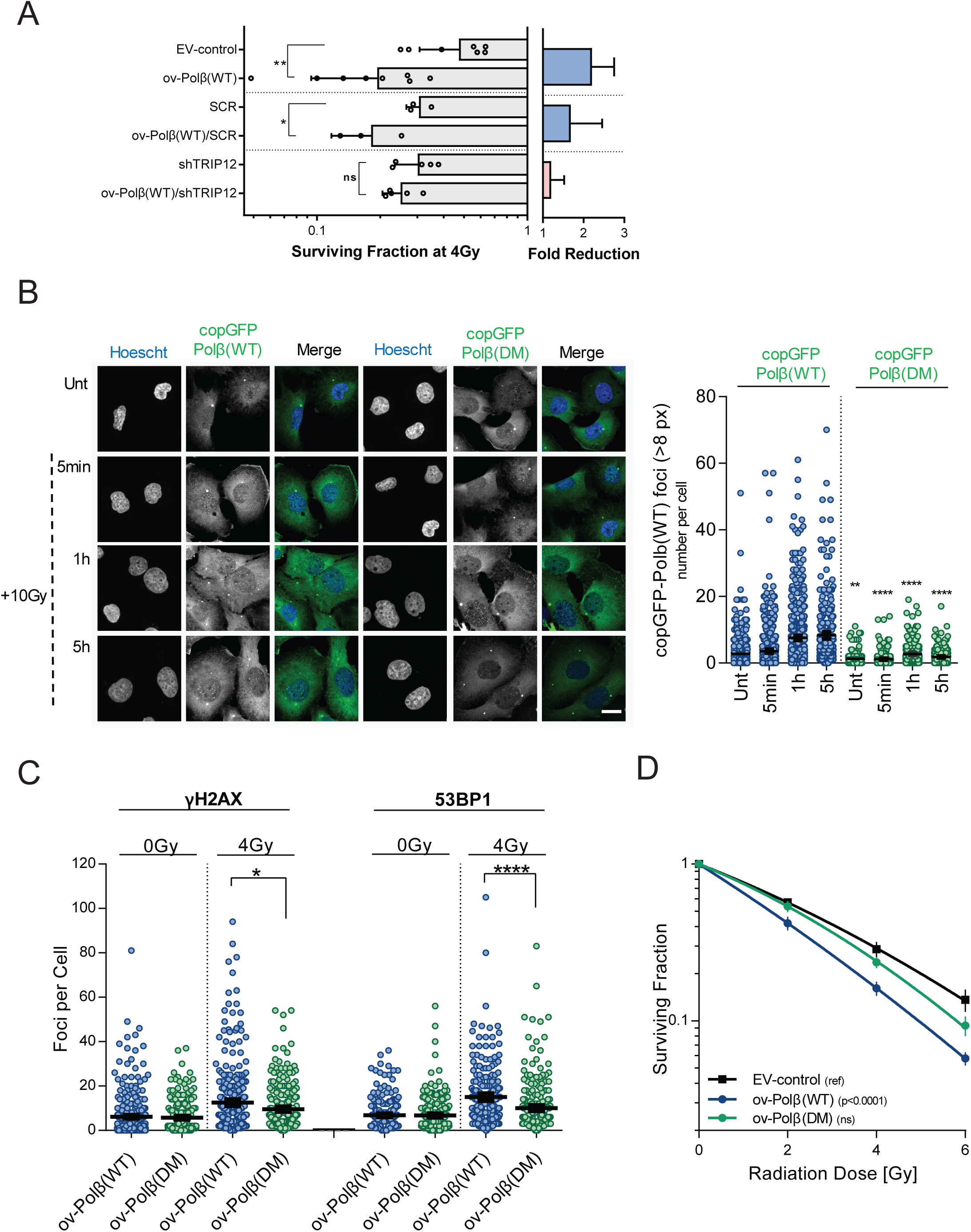
TRIP12 governs radiation response through Polβ ubiquitylation. **(A)** TRIP12 depletion rescues cells from Polβ over-expression mediated radiosensitization. Reduced survival after radiation by the over-expression of wildtype Polβ(ov-Polβ(WT)) (Figure 5D) in empty vector (EV-control) or SCR (scrambled shRNA) controls is rescued by TRIP12 depletion (TRIP12-sh1 and -sh5). Graph shows the average clonogenic survival after 4Gy and the SD in n=3-7 independent experiments as indicated by the dots. The Polβ over-expression induced fold reduction in radiation survival is indicated to the right; *p<0.05 and **p<0.01 (ANOVA). **(B)** Mutation of the TRIP12 ubiquitylation sites (K206A/K244A) on Polβ abrogates Polβ focal accumulation after radiation. Representative images and quantifications of radiation-induced Polβ (copGFP-Polβ(WT)) and ubiquitylation mutant Polβ(DM) (copGFP-Polβ(DM)) foci. Graph shows the number of large (>8px) mutant or wildtype Polβ foci per cell at different time points after radiation from n=3 independent experiments with means and SD; *p<0.05 and **p<0.01 (ANOVA). **(C)** Abrogation of TRIP12 ubiquitylation sites on Polβ reduces DSB induction. Residual ɣH2AX and 53BP1 foci at 24h are shown in untreated and with 4Gy irradiated cells that overexpress wildtype Polβ (ov-Polβ(WT)) or the Polβ K206A/K244A ubiquitylation mutant (ov-Polβ(DM); *p<0.05 and **p<0.01 (ANOVA). **(D)** TRIP12 ubiquitylation site mutation in Polβ rescues cells from Polβ over-expression mediated radiosensitivity. Polβ(WT) over-expressing cells are compared to Polβ(DM) over-expressing and empty vector (EV-control) LN428 control cells. N=3-8 independent experiments with mean and SEM. P values indicated in the radiation response curve graphs assess the likelihood of the data curve fits to be like the reference EV-control response curve. Radiation response parameters (D37%) in the ov-Polβ(DM) cell line is not significantly different from the EV-control (ns), compared to the ov-Polβ(WT) that differ significantly from the reference (ref) EV-control cell line with p<0.01(ANOVA).

### TRIP12 has a trafficking role in DNA damage repair

Despite the observed decrease and deregulation of Polβ in chromatin, TRIP12 depletion also causes extended RNF168-mediated chromatin signaling that leads to larger 53BP1 foci (20). Together, our data suggests that TRIP12 can direct DNA repair activity at lesion sites by (i) temporarily restraining chromatin ubiquitylation which fosters DSB repair while (ii) concomitantly engaging Polβ for BER at these sites. Testing this role in local repair segregation further, we find that TRIP12 restrains nuclear co-localization of 53BP1 and Polβ (**Figure 7A**). The data further confirms that TRIP12 is an early mediator of 53BP1 and Polβ segregation at radiation-induced lesions, as indicated by the significant drop in the Mander’s overlap coefficient as early as 30min post irradiation in TRIP12 expressing cells (**Figures 7A, S5C**, with p<0.0001).

**Figure 7.**
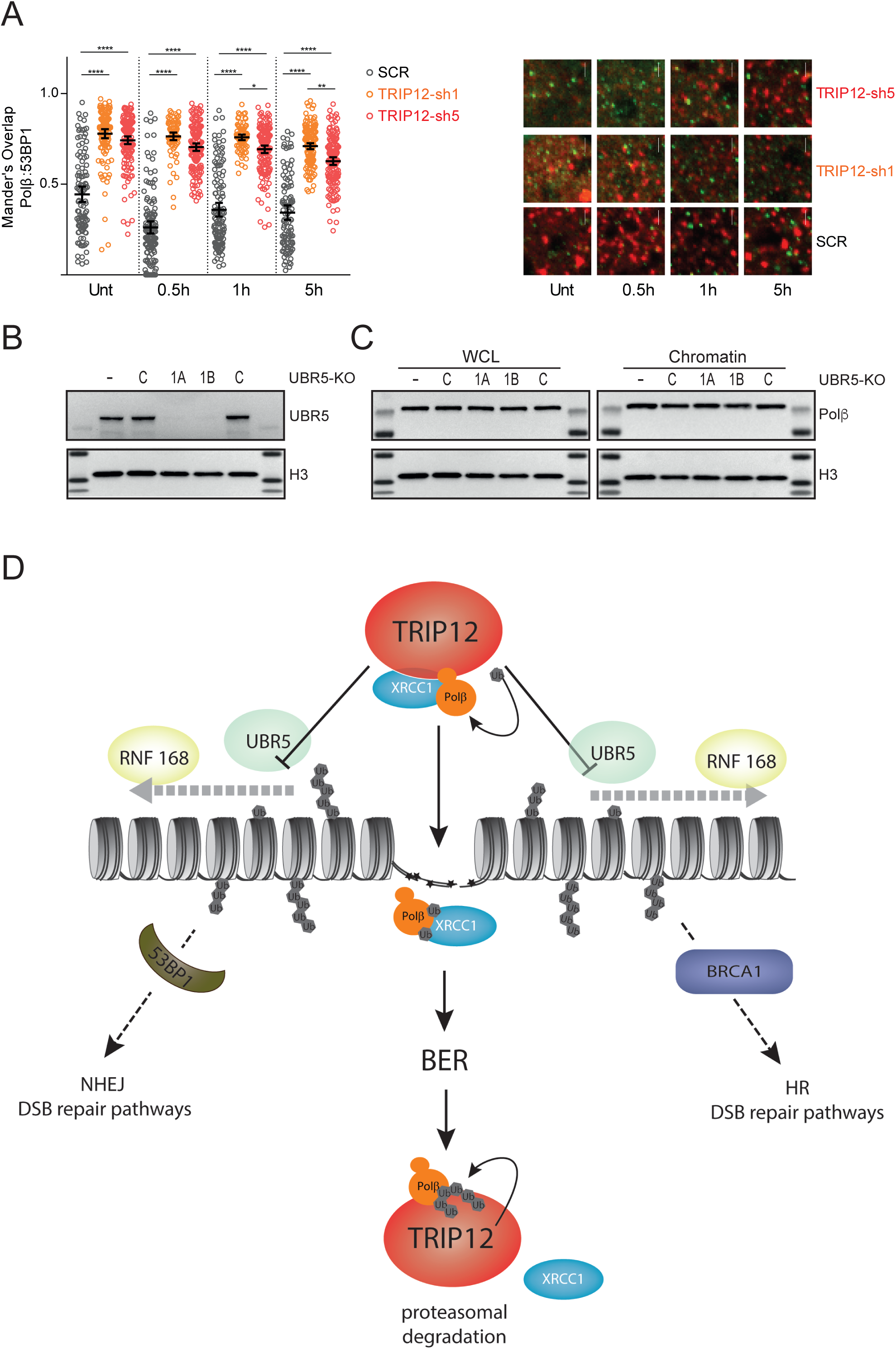
TRIP12’s controller function in DNA repair pathway engagement and choice. **(A)** TRIP12 depletion enhances co-localization of Polβ and 53BP1. copGFP-Polβ (green) expressing and scrambled (SCR) control or TRIP12 depleted (TRIP12-sh1 and -sh5) LN428 cells, pre and post (Unt, 0.5, 1, and 5 hours) irradiation (10Gy), were imaged and stained for 53BP1 by immunofluorescence (red) after fixation. Shown are representative images of the stained and imaged nuclei (right) and the quantified analysis (left, Mander’s overlap coefficients of individual nuclei with mean and 95% CI; *p<0.05, **p<0.01, ***p<0.001, ****p<0.0001 from ANOVA) demonstrating TRIP12-controlled local segregation of Polβ and 53BP1. **(B)** Loss of UBR5 expression as determined by immunoblotting of whole cell lysates of two different UBR5-KO cell lines (1A and 1B) as compared to parental LN428 cells (-) and control gRNA cells (C) and to Histone H3 as loading control. **(C)** UBR5 loss does not affect cellular Polβ levels or chromatin-associated Polβ levels. Polβ levels in (left) whole cell lysates (WCL) or (right) the chromatin fraction from parental LN428 cells (-), control gRNA cells (C) and two different UBR5-KO cell lines (1A and 1B). The WCL immunoblot is identical to (B) showing identical histone H3 loading control images and is repeated for comparison purposes. **(D)** TRIP12-mediated prevention of the RNF168 promoted histone ubiquitylation extension surrounding DNA lesions restrains DNA double-strand break signaling in favor of Polβ. Enabled by its ubiquitylation activity on Polβ, TRIP12 exerts its repair traffic control function by assisting BER and deviating DSB repair at the same time. Complex damage sites with clustered lesions can cause DSBs if not repaired appropriately and DSB repair attempts at DSBs residing in such regions may fail due to the presence of such BER-targeted lesions since they prevent synthesis and nuclease activities. A cellular control mechanism that channels DDR and SBR to yield the right of way to BER may be therefore relevant in these cases. A two-step activity can be proposed in which TRIP12 engages Polβ at nuclear damage sites but also initiates BER complex disassembly, freeing Polβ from XRCC1 for distinct repair activities and thereafter destines Polβ for proteasome degradation upon repair completion.

Another contender that could support this trafficking activity is the E3 ubiquitin ligase UBR5, reported, in conjunction with TRIP12, to suppress RNF168-facilitated 53BP1 chromatin loading (20). We find that in contrast to TRIP12, UBR5 loss (**Figure 7B**) does not alter cellular Polβ levels or its chromatin loading (**Figure 7C**). Since UBR5 also suppresses RNF168, this therefore also excludes indirect effects caused by excessive spreading of ubiquitylated chromatin. The uncoupling of UBR5 in the TRIP12 governance of Polβ-mediated repair may enable distinct responses and allow for lesion, repair and/or cell cycle phase specificity. Furthermore, it underscores TRIP12’s unique and active trafficking role in Polβ engagement and degradation (**Figure 7D**).

## Discussion

The data presented here grant TRIP12 a central molecular role in DNA damage response and repair pathway orchestration and reveal an important function in Polβ engagement and BER homeostasis. We show that TRIP12 binds to Polβ, through ubiquitylation, controls cellular levels of Polβ and governs Polβ chromatin retention upon DNA damage. Together, TRIP12 regulates Polβ involvement in the maze of repair pathways in response to DNA damage. The herein discovered trafficking role for TRIP12 at the nexus of DNA damage response and repair pathways towards Polβ-directed BER results in the segregation of Polβ directed repair from RNF168/53BP1 initiated DNA damage repair. The previously reported TRIP12-mediated containment of RNF168 and 53BP1 (20) thus further supports its function in Polβ engagement and BER at lesion sites and repair phases (see **Graphical Abstract**). Such a ‘repair-controller’ or trafficking role is particularly important during replication and at complex lesion sites to assure optimal DNA repair pathway choice and sequence to prevent damage aggravation.

### Importance of DNA repair pathway orchestration

The vital importance of genome maintenance is underscored by the evolution of multiple genome repair mechanisms, each of which function on a specific type or class of damaged DNA. Of these, the BER pathway plays a critical role in repairing the most abundant type of DNA lesions. DNA SSBs, nicks, and replication-blocking base lesions are the most crucial among BER targets. If not repaired, they can give rise to DNA double-strand breaks (DSBs) that ultimately require repair by non-homologous end-joining (NHEJ) or the homologous recombination (HR) pathways, depending on the context they arise. BER-targeted lesions may also occur during replication or pose a threat to genomic regions undergoing active DNA synthesis, including those regions involved in HR.

Struggling with endogenously induced lesions and ubiquitous exposure to ionizing radiation that causes a broad spectrum of base lesions and high numbers of SSBs, cells rely on a network of DNA repair pathways. Ionizing radiation induces clustered DNA lesions with base damage, SSBs and DSBs in close proximity. They are therefore particularly challenging lesions to repair that require a well-coordinated BER mediated strand incision sequence to prevent DSB formation (49–53). Since the complexity of this type of damage also hampers DSB repair if located close to or at DSBs, cellular mechanisms that direct the order of repair activities are critical for genome maintenance. A prioritization of BER and the prevention of, in this case, futile DSB repair, may also be important for cellular survival. Similar arguments apply to BER-targeted lesions that would otherwise block DNA synthesis and replication. Functional or physical crosstalk among DNA repair pathways is emerging as an essential aspect of the overall cellular response to DNA damage (54–57). This has been extensively described for repair pathways dealing with DSBs or crosslinks. However, the BER/DSB repair pathway orchestration postulated here has not yet been defined and links repair pathways across classes that are intrinsically very different. Our study portrays TRIP12 as a repair coordinator and controller at the nexus of the main cellular repair activities (**Figure 7D**, **Graphical Abstract**).

The recently discovered binding of TRIP12 to PARP1, another important BER and SSBR member, and its ability to regulate steady state levels of PARP1 through polyubiquitylation, is in line with our model and our proposed repair coordination and control function (21). Like the notion elucidated by us, the authors suggested that the relevance of this process lays in the prevention of supra-physiological PARP1 accumulation and activity. Indeed, in our study we were able to demonstrate the detrimental effects of supra-physiological Polβ levels in our over-expression models and thus a role for TRIP12 in genomic stability.

### TRIP12 trafficking role in radiation damage repair

Our study uncovers the cellular function of TRIP12 and identifies TRIP12 as a key regulator of BER repair activity through Polβ engagement. Here, we show ionizing radiation induced Polβ foci formation and reveal that TRIP12 is essential for this Polβ engagement in the cellular response to radiation. TRIP12 requirement is evident for those foci that arise under unchallenged conditions and may be, in part, replication-associated (time 0 in **Figure 4** and a fraction of foci in **Figure S4C** at this time point) and for those that were induced by radiation (**Figure 4**). Polβ recruitment at lesions induced by laser irradiation intensities that do not cause DSBs, or clustered lesions, is however not affected (**Figures 3C** and **S3C**), nor is the small induction of Polβ foci by cisplatin which also does not directly induce DSBs (**Figure S4E-F**). In contrast, but consistent with the herein proposed trafficking role of TRIP12 at the nexus of BER and DSB repair, TRIP12 is clearly involved in foci induction after radiation that causes complex base and DSB lesions (**Figures 4D** and **4E**). The lack of Polβ co-localisation with key DSB repair markers such as γH2AX and 53BP1 when TRIP12 is expressed is also consistent with the proposed restraining role of TRIP12 at such sites (**Figures 4B and 7A**). This functional link between TRIP12 and Polβ is further strengthened by the inability of Polβ mutants, devoid of TRIP12-mediated ubiquitylation (**Figures 2C and S2A**), to form Polβ foci at background levels or after radiation, consistent with the TRIP12 knockdown data (**Figure 6B**). More so, we were able to confirm TRIP12’s involvement in the nuclear activities of Polβ by showing the rescue in DSB induction and the radiation survival detriment from Polβ deregulation in TRIP12 ubiquitylation mutants.

Initially, and despite the drastic reduction in Polβ engagement, we were not able to identify cellular defects in TRIP12-deficient cells (**Figure 3E-3G**) except for elevated expression of RNA168 (**Figure S3B**). This lack of an impact on cellular survival or radiation sensitivity is not surprising as Polβ-deficient cells have been reported to survive well *in culture* and show similar radiation survival as their wildtype counterparts (29,58–61). Indeed, a role for Polβ in the cellular radiation response could only be identified after isolating replicating cells or applying dominant negative transgenes that interfered in BER (59–61). Overall, this observation is also consistent with earlier reports from Gudjonsson et al (20), who did not report a deficit in cellular survival upon depletion of TRIP12 and from Kajiro et al that describe a minor impact of TRIP12 HECT domain mutations on proliferation of ES cells (62). The lack of radiation sensitivity is also in-line with the notion that TRIP12 deficiency unlocks DSB repair opportunities (20,63), which may rescue any potential cellular impact from DSBs caused by the loss of Polβ-mediated repair. Together, this further suggests that TRIP12 deficiency is tolerable for cellular survival despite its detrimental effects on Polβ engagement. However, an important impact on genome maintenance can be deduced from the observed changes in DSB repair, a predicted preference towards the inherently error prone NHEJ pathway by deregulated 53BP1 and from reports that show an impact of TRIP12 haploinsufficiency on nervous system development and function (64,65). Consistent with this phenotype, Onishi et al reported neuronal differentiation defects in the absence of Polβ (66). Together, the TRIP12 repair trafficking role also solves the conundrum as to its repair restraining activity and points to repair governance beyond spatial restriction (63).

### TRIP12’s role in Polβ homeostasis

Polβ and XRCC1 are important elements in BER; their equilibrium appears to be critical for BER function (3,36,37,67). Of these two, Polβ provides both DNA polymerase and 5’dRP lyase activities essential to complete base lesion repair (27,29,68). By performing co-localization experiments after radiation, we observed that Polβ resides at most XRCC1 sites (**Figure 4B**), underlining the relevance of XRCC1/Polβ complex formation (37). Our previous findings and reports from others revealed that both Polβ and XRCC1 play common but also separate and independent roles in DNA metabolism and repair (3,69). We discovered that the composition of BER protein complexes depends on the cell cycle and DNA damage type (3) and observed a shift towards HSP90-bound XRCC1, devoid of Polβ. Since Polβ levels remained constant it suggested a considerable pool of a Polβ-centric BER complex (herein termed Complex B) and a mechanism to mediate this shift that depended on Polβ as it was absent in Polβ knockout cells (3). The considerable increase in Polβ degradation, when unable to bind to XRCC1 (Polβ(TM)), further substantiated the suspicion of a complex-dependent and Polβ-specific E3 ligase activity in these previous studies (36). Herein, we identified, via label-free dMS, TRIP12 as this complex dependent E3 ubiquitin ligase. Polβ is a substrate for TRIP12 both *in vitro* and in cells and the interaction is between the substrate binding domain (TRIP-SB) of TRIP12 and the C-terminal domain of Polβ. At this stage, we are not able to disentangle, whether TRIP12 activities can contribute to a shift towards the XRCC1-free complex or whether directed by other mechanisms, this state enables TRIP12 interference, thus ubiquitylation and Polβ foci formation. We did however observe a reduction in cytoplasmic Polβ levels after radiation at late time points (**Figure 4AC**). Together with the expected overall increase in levels due to the lack of degradation processes, this reduction was absent in TRIP12 knockdown cells. TRIP12’s contribution to Polβ elimination supports a role in re-establishing baseline values after successful repair, further highlighting the importance of well-balanced Polβ levels. This may address the observed paradoxical role of TRIP12 in promoting Polβ engagement in the nucleus in response to damage on one hand while also facilitating Polβ degradation after the insult.

### Relevance of Polβ homeostasis and Polβ over-expression

Early studies showed that Polβ expression is regulated in a highly tissue-dependent manner. While some tissues overexpress Polβ such as testis, cerebellum, tonsil, gall bladder and salivary gland, others show little expression of Polβ, such as heart and smooth muscle (70). The very distinct spatial and temporal expression pattern during mouse development further demonstrates the existence of regulatory mechanisms that assure tightly controlled Polβ levels (66). From genetic manipulation studies we can conclude that the lack of Polβ is incompatible with life and results in neonatal death and enhanced neuronal cell apoptosis (29,66). Importantly, constituently over-expressed Polβ also results in normal tissue development defects (accelerated onset of cataract) and enhanced tumor occurrence (30,31) and, reversely, constituently decreased Polβ levels result in growth retardation in a mouse model that expresses XRCC1 binding impaired Polβ (36). Here we show that over-expression causes DSBs, the induction of RPA foci and HU recovery defects that point to an interference and involvement in replication-associated processes, a postulation also brought forward by others (71). The observed damage reflects the consequences of unscheduled and exacerbated activity of the normal cellular involvement of Polβ in DNA metabolism that may occur in some tissues if not controlled locally. The lack of EDU co-localisation does not discount an involvement in replication as these were assessed in a non-over-expressed model and are radiation induced Polβ foci (at complex lesions). Interestingly, in this context we find RFC1 to be one of the Complex B specific partners (**Supplementary Table S2**). We were also able to demonstrate the harmful effects of deregulated Polβ levels by over-expressing Polβ and showing increased DSB formation and radiation sensitivity. Since we see a dependence on TRIP12-mediated ubiquitylation, the prevention of DSB repair initiation through a TRIP12-governed chromatin lock may be the cause of the replication associated DSBs raised by blocking lesions. A link difficult to prove as TRIP12 also governs Polβ loading. Interestingly, high expression of Polβ has been demonstrated in many cancer tissues including HNSCC (72–77) and a study reports that down-regulation of TRIP12 caused radio-sensitization in HPV-positive HNSCC cell lines (24).

### Polβ in TRIP12 governed processes

Little is known about the cellular role of TRIP12, a HECT-type E3 ubiquitin ligase. ARF and App-BP1 have been reported to be among TRIP12 ubiquitylation targets, implicating TRIP12 in ubiquitin fusion degradation, neddylation or ARF regulated p53 response (44,78–80). The first indications for an involvement in DNA repair processes came from data revealing a role in 53BP1 foci formation (20) and a link to the DNA damage response through USP7 ubiquitylation (26). Gudjonsson et al also showed that TRIP12 acts as a suppressor of RNF168, reducing the accumulation of ’Lys-63’-linked histone H2A and H2AX at DNA damage sites and revealed its partnership with UBR5. TRIP12 thereby acts as a guard against excessive spreading of ubiquitylated chromatin at damaged chromosomes that is thought to facilitate the accumulation of repair proteins (20,81). Here, we were able to juxtapose TRIP12’s DSB repair restraining function with a repair promoting activity through the direct engagement of Polβ, crucial for BER. The lack of co-localization of Polβ with 53BP1 (**Figure 4B**), unless TRIP12 is depleted (**Figure 7A**), supports this link. Interestingly, we find that in contrast to TRIP12, its partner UBR5 does not affect the chromatin association of Polβ (**Figure 7C**). The uncoupling of UBR5 in TRIP12’s activities regarding Polβ engagement and BER suggests a supportive modulatory function to further discriminate both functions.

Indeed, BER may not require extensive chromatin rearrangement as indicated by fast recruitment kinetics in heterochromatic regions (82). The repair of radiation-induced clustered lesions will benefit from early BER engagement and restricted chromatin rearrangement to prevent DSB formation or DSB repair attempts at lesion sites. In this context, Eccles et al showed the importance of, then unknown, nuclear factors in the sequence of events at such sites (83). At this point, we are not able to discern whether the late and large Polβ foci are a consequence of retention due to continued loading attempts by TRIP12 from incomplete repair or from the engagement in late, secondary DSB repair processes. The impact of TRIP12 deficiency in the early and small foci however confirms a loading role for most of these foci. This is also consistent with a damage-type specific trafficking role of TRIP12 as shown by the lack of such a response modulation after H_2_O_2_ or cisplatin. Since radiation induces a large amount of both Polβ and 53BP1 foci, the drop in Polβ/53BP1 co-localization reveals an active segregation process early after radiation that is TRIP12-dependent (**Figures 7A and S5C**). Combined with TRIP12’s shown Polβ shuttling activity, this early BER engagement and 53BP1 exclusion further supports our hypothesis that assigns TRIP12 a role in the sequence of repair activities. We postulate that the provenance of TRIP12 ubiquitylation activity on Polβ is to enable a timely order of repair by ranking BER activities over those from the DSB repair machinery through an active role in Polβ chromatin association and the concomitant regulation of initial DNA damage response modulators.

As described above and evident in the Polβ over-expression models, we have also seen a second TRIP12 dependent process: an influence on replication-associated processes that may account for the drop in background (un-irradiated) Polβ foci levels in the TRIP12-sh and Polβ(DM) cells. BER lesions are abundant and the conversion of such lesions into lethal DSBs or replication blocks can therefore be detrimental for the cell as highlighted by the synthetic lethal interaction of PARP inhibitors and HR defects (84,85). While barring HR, pending BER completion, will be of benefit for the cell, so would a control over replication preparation. A direct role of the TRIP12 target USP7 in replication has been suggested (86). Later studies elaborate on this link by showing that USP7 is necessary for replication-fork progression (87). Even though detailed elements of these processes have yet to be defined, a similar trafficking role as for DSB repair could be proposed here that favours BER over replication preparation via USP7, while DSB repair pathway choice is influenced by 53BP1 confinement in the respective chromatin context (88).

TRIP12’s reported DDR restraining function was puzzling at first (63). Here, we were able to settle the question as to why such a restraint may be important: in concert with its Polβ loading function, RNF168 determined boundaries allow the control of repair traffic at complex damage sites, ultimately providing a negative regulation of DSB repair (55) in favor of BER.

### Dual function of TRIP12: Repair pathway coordination and protein homeostasis

Here, we observed a clear role for TRIP12 to govern Polβ chromatin loading as well as cellular Polβ levels. Ubiquitylation plays a role in protein stability for many DNA repair proteins whereas non-proteolytic functions of ubiquitin (either mono-ubiquitin or poly-ubiquitin) are essential for signaling in DNA replication and DNA repair (89,90). As part of our dynamic BER model (3), we deduced that ubiquitylation of Polβ may facilitate repair pathway choice such as that seen with regard to ubiquitin in DSB repair processes (89,91). The poly-ubiquitin mark can play both a signalling role as well as facilitate proteolysis (92,93). Herein, we suggest that Polβ is trafficked by TRIP12-mediated ubiquitylation, possibly assisted by additional E3 ligases, that were identified to be Complex B enriched in the *dMS* screen and then further poly-ubiquitylated to regulate Polβ protein levels (**Figure 7D**). We observe TRIP12-dependent generation of both, mono- and poly-ubiquitylated species of Polβ (**Figures 2A-B**). This is in-line with a role for ubiquitylation in both signalling and protein homeostasis that may depend on ubiquitin chain types (*manuscript in preparation*). Consistent with a role for Polβ ubiquitylation in repair and signalling, we show that TRIP12 and the ubiquitylation of Polβ at K206/K244 is essential for Polβ chromatin association and foci induction (**Figures 3B, S3A, 6**) and in the maintenance of cellular Polβ levels (**Figure 2D**).

In conclusion, we propose that TRIP12 functions as a ‘repair-traffic-controller’, through ubiquitylation, facilitating pathway crosstalk across significantly different repair processes that require a different extent of chromatin modulation. By coordinating Polβ and DSB repair engagement, TRIP12 may ultimately prevent the conversion of base lesions and single strand breaks to cytotoxic and genotoxic DSBs.

## SUPPLEMENTARY DATA

Supplemental Information includes Detailed Methods, five figures (Supplementary Figures S1-S5) and two tables (Supplementary Tables S1, S2).

## Acknowledgements

This study was supported by funds from the National Institutes of Health [grant numbers CA148629, ES014811, ES029518 and CA238061 to R.W.S.] and the Dutch Cancer Society KWF [grant number NKI-2010-4877 to C.V.]. This study used the Hillman Cancer Center Proteomics Facility that is supported in part by award P30CA047904 from NIH. Support for the USA Mitchell Cancer Institute (MCI) Cellular and Biomolecular Imaging Facility and the USA MCI Flow Cytometry Facility was provided by funds from the Abraham A. Mitchell Distinguished Investigator award and USA MCI Core Facility funds. Support for the NKI microscopy facility was provided by operating funds of the Netherlands Cancer Institute (NKI). Support is also provided by the Legoretta Cancer Center Endowment Fund (to RWS). We thank S.H. Wilson (NIEHS/NIH) for the gift of recombinant human DNA polymerase β used for some of this project and B. Vogelstein for the gift of the HCT116 cells. We also thank T. Sixma for carefully reading the manuscript and offering constructive comments.

## Author Contributions

Conceptualization: C.V. and R.W.S.; Supervision: R.W.S., C.V. and N.A.Y. Methodology: Q.F, B.I., N.A.Y., J.A., J.L., X.Z., P.S., N.A.Y., J.J., M.V., R.W.S. and C.V.; Investigation: B.I., Q.F. J.A., M.I., J.C., J.L., P.S., Z.Y., A.B., C.V. and R.W.S.; Data analysis and synthesis: Q.F., C.V., B.I., R.W.S and N.A.Y., Resources: R.W.S., C.V., N.A.Y., J.J. and M.V. Writing original draft: Q.F., B.I., R.W.S. and C.V.; Writing-editing: C.V. and R.W.S.; Funding Acquisition: R.W.S, C.V., N.A.Y., J.J. and M.V.

