## Supplementary Material for "TRIP12 governs DNA Polymerase β involvement in DNA damage response and repair"

Supplemental Material includes:

Supplementary Document 1 - Method Details
Supplementary Figures S1-S5 with corresponding figure legends
Supplementary Tables S1, S2

#### Supplementary Document 1

**TRIP12 governs DNA Polymerase β involvement in DNA damage response and repair**

Burcu Inanc, Qingming Fang, Joel Andrews, Xuemei Zeng, Jennifer Clark, Jianfeng Li, Nupur B. Dey, Md Ibrahim, Peter Sykora, Zhongxun Yu, Andrea Braganza, Marcel Verheij, Jos Jonkers, Nathan A. Yates, Conchita Vens and Robert W. Sobol

Supplemental information includes: Method Details

##### Plasmid and vector development

Plasmids and lentiviral vectors developed previously, obtained commercially or from colleagues are all cited in the **Supplementary Table S1**. Gateway-ready ENTRY vectors (Thermo Fischer Scientific) encoding TRIP12 (and the indicated TRIP12 fragments) with an N-terminal Myc-tag were prepared by GenScript USA Inc. Gateway-ready ENTRY vectors encoding the C-terminal domain (AA 91-335) of Polβ(WT), Polβ(TM) or Polβ(TM/DM); each with an N-terminal Flag-tag, were prepared as follows: the corresponding Flag-Polβ-C fragment was PCR amplified (using oligonucleotides Flag-Polβ-C-F and Flag-Polβ-C-R) from pENTR-Flag-Polβ (WT), pENTR-Flag-Polβ(TM) or pENTR-Flag-Polβ (TM/DM) (see **Supplementary Table S1**) and cloned into pENTR/D-TOPO via TOPO cloning. pENTR-Myc-HECT(C2007A) was constructed by mutating residue C2007 to C2007A with the Quickchange II XL site-directed mutagenesis kit and the primers are listed in the **Supplementary Table S1** (HECT-C2007A-FW and HECT-C2007A-Re).

The pLVX lentiviral expression vectors were developed using the pENTR plasmids listed in **Supplementary Table S1** by transferring the encoded open reading frame into a Gateway-modified lentiviral vector (pLVX-GWB-IRES-puro, pLVX-GWB-IRES-Neo vector or pLVX-GWB-IRES-Hygro) via an LR reaction using the Gateway LR Clonase II enzyme mix as per the manufacturer's instruction, as we have described (1). Positive clones were selected, and plasmids were extracted with the QIAprep Spin Miniprep Kit (Qiagen).

The lentiviral expression vectors for expression of copGFP-Polβ(WT) and copGFP-Polβ(TM) fusion proteins were constructed and developed previously (2). Using the same protocol, we constructed lentiviral vectors for expression of copGFP-Polβ(DM) and copGFP-Polβ(TM/DM). These copGFP-Polβ fusion plasmids were constructed using the parent vector, pCT-CMV-copGFP-MCS-EF1-puro (System Biosciences). Forward (GFP-PolβC24F) and reverse (GFP-PolβC24R) primers were designed (see **Supplementary Table S1**). The Polβ open reading frame for the corresponding mutant was then PCR-amplified to engineer the restriction enzyme sites XbaI and BamHI for in-frame cloning using standard protocols, with pENTR-Flag-Polβ(DM) or pENTR-Polβ(TM/DM) as template. PCR products were digested with XbaI and BamHI and purified fragments were ligated into the pCT-CMV-copGFP-MCS-EF1-puro lentiviral vector (previously digested by XbaI and BamHI). Positive colonies were selected and sequenced. Plasmids with the correct sequence were used for lentivirus production as described below.

Lentiviral vectors expressing the scrambled shRNA (SCR) and each of five shRNAs specific to TRIP12 were from Sigma and prepared as we have described previously (2,3). The sequence of each shRNA is listed in the **Supplementary Table S1**.

**Lentivirus production and cell transduction**

Lentiviral particles were generated by co-transfection of 4 plasmids into 293-FT cells using FuGene 6 Transfection reagent (Promega, Cat# E2311): the packaging vectors pMD2.g(VSVG), pVSV-REV and pMDLg/pRRE together with the appropriate shuttle vectors, as listed in the **Supplementary Table S1**. Forty-eight hours after transfection, lentivirus-containing supernatant was collected and passed through 0.45 mM filters to isolate the viral particles as we described previously (2-4).

Lentiviral transduction was performed as follows: cells (1-1.5x$10^5$) were seeded into 6-well plates. 24 hrs later, lentiviral particles were mixed with polybrene (2μg/ml) and added to the cells. Cells were incubated at 32˚C overnight. Medium with lentiviral particles was removed and replaced with fresh medium. Cells were cultured for 48 hrs at 37˚C before selection with antibiotics puromycin (Sigma-Aldrich, Cat# P9620), hygromycin (Sigma-Aldrich, Cat# G1272) or G418 (Corning, Cat# 30-234-CI), for 1-2 weeks. Whole cell lysates were analyzed by immunoblotting to validate the expression of the desired proteins. All cells were cultured at 5% $CO_2$ and 37˚C. All the cell lines developed and used in this study are listed in the **Supplementary Table S1**.

##### Plasmid transfection

Transient transfections using the FuGene 6 transfection reagent (Promega) were performed when using the plasmid vectors pcDNA-HA-ubiquitin, pRS1436 or pRS1427. Briefly, plasmid DNA (12μg) was mixed with 36μl of transfection reagent and incubated for 30 min at room temperature. The mixture was then added to the cells, mixed gently and the cells were then cultured for 48 hrs. Cells were collected and cell lysates prepared as described below.

##### Cell extract preparation

Cell extracts (whole cell lysates, WCL) were prepared from cells with different genetic modifications and/or treated with different drugs and for different times as indicated in the text. Cells (2.5-10 x$10^5$) were seeded into a 60-mm cell culture dish. After further incubation for 24hrs, cells were treated as indicated, washed twice with cold PBS, collected and lysed with an appropriate volume of 2x clear Laemmli buffer (2% SDS, 20% glycerol, 62.5mmol/l Tris-HCl pH6.8). Cell lysates were boiled for 10 min and quantified with the DC protein assay kit following the microplate protocol provided by the company (Bio-Rad).

##### Immunoprecipitation

To screen for binding partners of Polβ or the Polβ/XRCC1 complex, anti-Flag M2 affinity gel was used to immunoprecipitate proteins from cell lysates of LN428/Flag-Polβ(WT), LN428/Flag-Polβ(TM) or LN428/EGFP cells. Briefly, confluent cells (70-90%) from two 150mm dishes were collected and lysed in 1ml of Pierce IP lysis buffer (Thermo Fisher Scientific) with protease inhibitor. The anti-Flag M2 affinity gel (Sigma-Aldrich) was prepared according to the protocol provided by the company. The Flag M2 affinity gel was washed three times with 5x bead volume of 0.1M Glycine pH3.5, three times with 5x bead volume of TBS buffer (50mM Tris-HCl pH7.4, 150mM NaCl), and twice with 5x bead volume of IP lysis buffer. The gel was suspended in 150μl of IP lysis buffer with protease inhibitor and mixed with 1ml cell lysate. The mixture was shaken overnight at 4˚C. The next day, the mixture was centrifuged to pellet the gel and the supernatant was removed. The gel was then washed five times with 1ml IP lysis buffer and three times with 1ml TBS buffer containing protease inhibitors. Finally, 50μl of sample buffer was added to the gel and the gel was boiled for 5 min. Each gel was centrifuged and the supernatant that contained the eluted IP product was collected. In total, 27 individual IP samples were prepared (n=9/condition) and a 25μl aliquot of each supernatant was sent for label-free differential mass spectrometry (dMS) analysis (see below). The remainder of the IP product was used for immunoblotting to confirm the dMS data.

To study the interaction of TRIP12 with Polβ, TRIP12 antibody was used to immunoprecipitate endogenous TRIP12; Myc antibody was used to immunoprecipitate Myc-TRIP12, Myc-TRIP12-SB, Myc-HECT, and Myc-HECT(C2007A); anti-Polβ antibody (Clone 61) was used to immunoprecipitate endogenous Polβ; M2 affinity gels were used to immunoprecipitate Flag-Polβ, Flag-Polβ mutants and Flag-Polβ fragments. The rabbit IgG anti-TurboGFP or mouse IgG anti-GFP antibodies were used as negative controls. The cell lysis and agarose preparation, binding and washing and the elution was performed as described above.

To study the ubiquitylation of Polβ in cells, immunoprecipitation was performed with the corresponding primary antibody (as indicated in the figures) and protein G (Santa Cruz Biotechnology, Cat # sc-2003). Protein G was washed twice with TBS buffer and pelleted by centrifuging at 10,000rpm for 20 sec at 4˚C. The cell lysates were prepared as described above. The mixture of cell lysates with protein G was shaken for more than 4 hrs at 4˚C. Thereafter, beads were washed three times with TBS buffer containing protease inhibitors. Finally, 30μl of sample buffer was added to the beads. Beads were boiled for 5 min to elute the immunoprecipitation products. The levels of pulled down ubiquitylated Polβ or loading Polβ levels were examined by immunoblot with antibodies to HA and M2, respectively.

**Immunoblot**

Whole cell lysates (8-20μg) or 5-15μl of immunoprecipitated proteins were loaded onto precast NuPAGE® Novex® 4-12% Bis-Tris gels (Invitrogen), run 2-3hrs at 130V. Gel electrophoresis separated proteins were transferred onto a 0.45μm pore-size nitrocellulose membrane (Bio-Rad) for 2-3 h or overnight, at 0.35A. The membrane was first washed with TBS supplemented with 5% blotting grade non-fat dry milk (Bio-

Rad) (1 hr, room temperature) and subsequently blotted with the primary antibodies in B-TBST (TBS buffer with 0.05% Tween-20 and supplemented with 5% blotting grade non-fat dry milk) for 2 hrs at room temperature. The primary antibodies and their dilutions are listed in the **Supplementary Table S1**. After washing, membranes were incubated with secondary antibodies in B-TBST for 1 hr (room temperature). The following HRP conjugated secondary antibodies were used: Immun-Star Goat anti-mouse-HRP conjugate and Immun-Star Goat anti-rabbit-HRP conjugate (see **Supplementary Table S1**). After washing, the membrane was illuminated with Immun-Star HRP peroxide buffer with luminol/enhancer firstly. If no signal was detected, SuperSignal west femto maximum sensitivity substrate (Thermo Fisher Scientific) was used to allow the detection of the secondary antibody labelled bands. If necessary, protein bands were quantified using Image J software (Image J 1.48v, http://imagej.nih.gov/ijjava 1.6.0_65).

**Mass spectrometry analysis**

The immunoprecipitates were loaded on SDS-PAGE gels (NuPAGE® Novex® 4-12% Bis-Tris Protein Gels) and proteins were separated from low molecular weight reagents using short gel (~1cm) fractionation (5,6). The gels were stained by Coomassie blue. Gel regions were excised as indicated and processed for tryptic digestion as previously described (7). Liquid-chromatography Fourier transform mass spectrometry was used to measure the mass-to-charge (m/z) ratio, retention time, and intensity of more than 100,000 peptide signals that are detected in high-resolution full-scan mass spectra. Briefly, a 1µl aliquot of tryptic peptides extracted from each IP product was separated by reverse-phased nano-flow liquid-chromatography (EASY-nLC II, Thermo Scientific, San Jose, CA) and analyzed on a LTQ/Orbitrap Velos Elite hybrid mass spectrometer (Thermo-Fisher, San Jose, CA). Solvent A (0.1% formic acid in HPLC grade water) and solvent B (0.1% formic acid in 100% acetonitrile) were used as the mobile phase. Peptides were first loaded onto a 20µl capillary sample trap column and desalted online with a 6µl volume of solvent A and loaded on a homemade slurry-packed capillary column (75µm ID x 360µm OD x 150mm) that contained C-18 silica-bonded stationary phase (5µm diameter, 300Ǻ pore size). A 100 min solvent gradient (0-90 min, 3-33%B, 90-92 min, 33-80%B, 92-98 min, 80% B, 98-100 min, 80-0%B) and 0.2µl/min flow rate were used to elute and ionize peptides prior to mass spectrometry (MS) analysis. All MS data was collected using positive electrospray ionization mode, with an FTMS1 AGC targets = 100000, a maximum injection time = 200ms, an electrospray ionization voltage = 2.5kV, and a capillary temperature = 325°C. The data dependent scan mode was used to concurrently acquire one high-resolution full scan mass spectra at a resolution setting of 60,000 and 20 low-resolution tandem mass spectra (MS/MS) every three seconds for the duration of the 100-minute analysis.

**Label-free dMS analysis**

Mass spectrometry data collected for each sample was analyzed using dMS software (Infoclinika, Bellevue WA). Briefly, the high-resolution full MS spectra were aligned and the m/z, charge state, retention time and intensity data for all molecular features detected in the full scan mass spectra were integrated and matched to protein identification results. Data analysis was performed within CHORUS, a cloud computing data analysis suite (https://chorusproject.org/). The 170,758 dMS features considered in this study were selected using a retention time range of 15-80 minutes, a peptide molecular weight between 700 and 5000 Da, and a minimum of two isotopes. Low abundance features with a mean peak area below the 5$^{th}$ percentile were excluded from the study. The COMET data base search engine (8) implemented within Chorus was used to identify the peptides and proteins that were matched to each molecular feature using the following modifications: static modification of cysteine (carboxyamidomethylation, +57.02 Da) and variable modification of methionine (oxidation, +15.99 Da). The mass tolerance was set to 20 ppm for precursor ions and 0.8 Da for fragment ions.

For proteins identified by multiple peptides, practical selection criteria were used to choose a single peptide that provides a robust surrogate measure of relative protein abundance. The selection criteria were based upon the signal intensities and the strength of correlation with other peptides originating from the same protein. Proteins with single identified peptide, or proteins whose associated peptides displaying no strong correlation with each other (mean Pearson's correlation coefficient >0.5) were excluded from further statistical analysis. The intensities of each surrogate peptide were normalized to its maximum intensity so that each peptide had maximum value of 1.

Student's t test implemented in MATLAB® was used to determine the statistical significance of the difference on the abundance of identified proteins/features in different IP samples. Feature selection was based on an ANOVA test p-value cutoff of 0.001 and a minimum fold-change increase of 20 when comparing IP samples of Flag-Polβ(WT) or Flag-Polβ(TM) to EGFP. The abundance values in Flag-Polβ(WT) and Flag-Polβ(TM) samples were normalized to the Polβ level in each sample to calculate the WT/TM ratio used to assess the differential interaction between Flag-Polβ(WT) and Flag-Polβ(TM) samples.

**Recombinant Proteins**

Recombinant human DNA polymerase β, expressed in *E. coli* and purified, was a generous gift from S.H. Wilson (NIEHS/NIH).

Recombinant, His-tagged ubiquitin (His-Ub) was generated and purified as described (6). Briefly, the pDEST-17-His-Ub plasmid was transformed into One-Shot BL21(DE3)pLysS chemically competent cells. To induce the expression of the His-Ub protein, bacterial cultures were allowed to grow at 37˚C and 250 rpm to an $A_{600}$ = 0.5–0.7. At this time, isopropyl β-d-1-thiogalactopyranoside (Sigma Aldrich) was added to a final concentration of 1μm. The culture was allowed to grow for another 3.5 hrs. The bacteria were then pelleted at 6000g for 15 min at 4˚C, followed by resuspension of the pellet in Buffer A (50mM Tris-HCl, pH7.6, 100mM NaCl, 0.1mM PMSF, and 1 tablet protease inhibitor for 20ml of Buffer A). The bacterial suspensions were pelleted at 6000g at 4˚C for 15 min, supernatants discarded, and the pellets frozen overnight at −80˚C. The recombinant His-Ub proteins were purified using the TALON® metal affinity resin and HisTALON™ buffer (TaKaRa) set according to the manufacturer's protocol. The purified protein was dialyzed with Dialysis Buffer (50mM Tris-HCl, pH7.6, 100mM NaCl, 0.5mM EDTA and 5mM DTT), changing the buffer four times, every 6–8 hrs. The dialyzed proteins were concentrated using Amicon ultracentrifugal filter devices (Millipore Sigma), and protein concentrations were measured using the DC protein assay with BSA as the standard.

**Polβ stability and degradation**

To evaluate the role of TRIP12 on the stability of Polβ(WT) or Polβ(TM), LN428/Flag-Polβ(WT)/TRIP12-KD, LN428/Flag-Polβ(TM)/TRIP12-KD, LN428/Flag-Polβ(WT)/SCR and LN428/Flag-Polβ(TM)/SCR cells were seeded and after 24 hr treated with 0.2mM Cycloheximide (Cyc) (Sigma-Aldrich) or with Cyc (0.2mM) and MG132 (25µM) (SelleckChem) for the time periods as indicated in the Figure. Compounds were removed and whole cell lysates (WCL) were prepared and quantified as described above. Polβ, XRCC1, PCNA and TRIP12 levels were determined by immunoblotting and the intensity of the bands was quantified using the Image J program. Levels of PCNA and XRCC1 were used to control for loading differences. The changes in Polβ levels as defined by the ratio of Polβ/PCNA and of Polβ/XRCC1 to normalize for loading variation were averaged and are plotted.

***In vitro* ubiquitylation activity assay**

To assess ubiquitylation activity of TRIP12 and the HECT or substrate binding (SB) domains of myc-TRIP12, myc-HECT, myc-HECT(C2007A) and myc-TRIP12-SB were immunoprecipitated using an anti-myc antibody or the Myc-Trap agarose. The Myc antibody bound with its antigen were incubated overnight (4˚C), then incubated with protein G beads for 4 hrs. The Myc-Trap agarose beads were used as per the manufacturer's instructions (Chromotek). The beads were washed five times with TBS buffer and then washed twice with 500µl ubiquitylation reaction buffer (10mM Tris-HCl, pH7.5, 100mM NaCl and 0.5mM DTT). The *in vitro* ubiquitylation assay was performed in buffer mixture (30µl) containing 0.1µM E1 (Boston Biochem), 0.25µM of the E2 enzyme mix (Life Sensors, containing: UBE2S, UBE2L3, UBE2D3, UBE2N, UBE2A, UBE2D1, UBE2G2, UBE2Z), 1µM ubiquitin aldehyde (Boston Biochem), 0.75mg/ml His-ubiquitin and/or 1x Magnesium/ATP cocktail (20mM MOPS, pH 7.2, 25mM β-glycerophosphate, 5mM EGTA, 1mM $Na_3VO_4$, 1mM dithiothreitol, 75mM $MgCl_2$, and 0.5mM ATP), with or without 1000ng of purified Polβ and samples were incubated for 2 hrs at 33˚C while mixing the samples every 20 minutes. Samples were then centrifuged for 30 sec at 13,000rpm and the supernatant was transferred to a new tube containing 30µl of blue Laemelli buffer. The beads were then heated to 70˚C for 15 min. Anti-His, anti-ubiquitin and anti-Polβ Ab (Clone61) antibodies were used to detect ubiquitylation of TRIP12 and ubiquitylated forms of Polβ.

To determine whether the HECT domain of TRIP12 ubiquitylates purified Polβ and itself, the Myc-IP products from LN428/EGFP, LN428/Myc-HECT or LN428/Myc-HECT(C2007A) cells were separated into 4 parts for the following different procedures: (1) Ubiquitylation buffer incubated for 2 hrs at 33˚C; (2) Purified Polβ mixed with ubiquitylation buffer and incubated for 2 hrs at 4˚C; (3) Purified Polβ mixed with ubiquitylation buffer and incubated for 1hr at 33˚C followed by 1 hr at 4°C; (4) Purified Polβ, mixed with ubiquitylation buffer and incubated for 2 hrs at 33˚C. Ubiquitylation of purified Polβ was examined after gel-electrophoresis and immunoblotting with anti-Polβ Ab (Clone61), anti-ubiquitin and anti-His-tag antibodies. Auto-ubiquitylation of the TRIP12 HECT domain was examined after gel-electrophoresis and immunoblotting as described above with an anti-ubiquitin or anti-His-tag antibody.

**Determination of ubiquitylation activity in cells**

LN428/Flag-Polβ(WT)/TRIP12-KD, LN428/Flag-Polβ(TM)/TRIP12-KD, LN428/Flag-Polβ(DM)/TRIP12-KD or LN428/Flag-Polβ(TM/DM)/TRIP12-KD cells were transiently transfected with pcDNA-HA-ubiquitin. Cells were lysed 48 hrs later and ubiquitylated proteins were immunoprecipitated with the anti-HA antibody. Immunoprecipitates were submitted to gel electrophoresis and immunoblotting and probed with the HA antibody or anti-Flag (M2) antibody.

**Quantitative RT-PCR (qRT-PCR) analysis**

TRIP12 and Polβ mRNA levels were measured by qRT–PCR using the Applied Biosystems StepOnePlus system (2,6). Briefly, 80,000 cells were lysed and reverse transcribed using the Applied Biosystems Taqman Gene Expression Cells-to-CT kit. Analysis of mRNA expression was performed via the ΔΔCT method using TaqMan Gene Expression Assays for human Polβ and human TRIP12 (see Supplementary Table S1), normalized to the expression of human β-actin. Samples were run in triplicate and the results are the mean ± SD of all three analyses.

**Clonogenic assay**

Radiation sensitivity was assessed by the colony formation assay. Proliferating LN428 cells were plated at varying cell concentrations. After 5h, cells were exposed to gamma rays from a Gammacell-40 Exactor (Best Theratronics Ltd., Ottawa, Ontario, Canada) at a dose rate of approximately 1 Gy/min at room temperature. Cells were allowed to grow for another 14 days to form colonies. Plates were rinsed in PBS and colonies fixed and stained overnight in 6% glutaraldehyde and 0.5% crystal violet. Colonies consisting of approximately 100 cells or more were counted by eye under an inverted dissecting microscope. Survival was calculated relative to the plating efficiency of unirradiated controls.

$H_2O_2$-induced cytotoxicity was also evaluated using a clonogenic assay. For $H_2O_2$-induced cytotoxicity, cells from LN428/SCR (Control), LN428/TRIP12-sh1, LN428/TRIP12-sh2 and LN428/TRIP12-sh5 cell lines were seeded at 100cells/well in 6-well plates. After 24 hours, cells were treated with freshly prepared $H_2O_2$ in selection media (0, 10, 20, 40, 60, 80 µM). Cells were incubated for 10 days without removal of the compound. Cells were then rinsed with PBS after the removal of media. 1ml of 0.5% Crystal Violet in ethanol was added to each well and removed after colonies were stained for 2-3 min at room temperature. Wells were washed once with cold water and dried at room temperature. The results are reported as the surviving fraction calculated from the ratio of number of colonies in $H_2O_2$ treated wells to untreated wells.

**CometChip assay**

DNA damage in LN428, LN428/SCR, LN428/TRIP12-sh1 and LN428/TRIP12-sh5 cells was assessed using the CometChip (Trevigen/BioTechne), allowing for high-throughput 96-well measurement of a large number of cells in multiple treatment groups, as recently described (9). Cell size was established using the Countess® II Automated cell counter (ThermoFisher Scientific) with the 30 micron CometChips used in all experiments. Cells were seeded at 2000/cells per well of a 96-well plate. After 24 hours, cells were exposed to media containing the vehicle control (PBS in complete media), $H_2O_2$ and incubated at 37˚C (**Figure 3D**, 250µM, 30 min; Figure S3D (left), 0-250µM, 30 min) or MMS and incubated at 37˚C (**Figure S3D** (right), 0-4 mM, 30 min). Cells were washed with complete media and either a) immediately loaded onto the CometChip or b) left to recover (repair) in conditioned media at 37˚C, as indicated in the figure legends (**Figures 3D, S3D**, repair time = 60 min). Once the exposure or repair course was completed, cells were trypsinized and gravity loaded into the CometChip apparatus (Trevigen/BioTechne) for 30 minutes at 4˚C. Each well in the 96-well CometChip contains approximately 500 micro-wells. The CometChip was then sealed with low melting point agarose (0.8%, Thermo Fisher Scientific) and submerged in detergent-based lysis solution for 1 h at 4˚C. In all experiments the CometChip was run under alkaline (pH >13) comet conditions (200mM NaOH, 1mM

EDTA, 0.1% Triton X-100). Electrophoresis was conducted at 22V for 50 min at 4˚C. After electrophoresis, the CometChip was re-equilibrated to neutral pH using Tris buffer (0.4M Tris·Cl, pH7.4). DNA was stained with 1x SYBR® Gold (Thermo Fisher Scientific) diluted in Tris buffer (20mM Tris·Cl, pH7.4) for 30 min and de-stained for 1 h (20mM Tris·Cl, pH7.4). Image acquisition was conducted on the Celigo S imaging cytometer (Nexcelom Bioscience, Lawrence, MA) at a resolution of 1 micron/pixel. Image analysis was conducted using CometChip analysis software (Trevigen/BioTechne) with the box size set to 220x180 pixels with acquired data exported to Prism (GraphPad) for further statistical analysis.

**Chromatin fraction isolation**

Chromatin fraction isolation was performed as follows: 100mm plates with 80-90% confluent cells were washed twice with cold PBS. Cytoplasmic lysis buffer (300µl; 10mM Tris-HCl pH8.0, 0.34M Sucrose, 3mM $CaCl_2$, 2mM Magnesium acetate, 0.1mM EDTA, 0.5% Nonidet P-40, protease inhibitor) was added to the plates and cells were scraped and transferred to 2ml Eppendorf tubes. Cells were incubated for 15 min on ice and then were centrifuged for 10 min for 4000rpm at 4˚C. The supernatant contained the cytoplasmic fraction and was transferred to a new tube. Pellets were carefully washed with 300µl of wash-buffer (10mM Tris-HCl pH8.0, 0.34M sucrose, 3mM $CaCl_2$, 2mM magnesium acetate, 0.1mM EDTA, protease inhibitor) and then removed with a pipette without resuspension and centrifugation. Nuclei were lysed in 300µl of nuclear lysis buffer (20mM HEPES pH 8.0, 3mM EDTA, 10% glycerol, 150mM potassium acetate, 1.5mM $MgCl_2$, 0.1% Nonidet P-40, protease inhibitor) and incubated on ice for 30 min. Nuclei lysates were next centrifuged at 13000rpm for 10min at 4˚C. Supernatant is the nuclear fraction and was transferred to a new tube. The pellet was washed with 300µl of nuclear lysis buffer and then with 750µl nuclease incubation buffer (150mM HEPES pH 8.0, 10% glycerol, 50mM potassium acetate, 100mM KCl, 1.5mM $MgCl_2$, protease inhibitor) carefully with a pipette without resuspension and centrifugation. The pellet was resuspended in nuclease incubation buffer (75µl) with 1.5µl benzonase (Sigma-Aldrich) and incubated for 15 min at 37˚C and mixed the resuspension every 5min. The resuspension was centrifuged with 13000rpm for 15 min at 4˚C. Chromatin fraction (supernatant) was collected for immunoblotting analysis.

**DNA fiber assay and analysis**

LN428 cells were pulse-labeled with 50μM CldU for 20 min, washed three times with medium and incubated in 2mM HU for 90 min thereafter. After HU treatment cells were washed three times with medium and then pulse-labeled with 500μM IdU for 20min. Labeled cells were harvested, and DNA fiber spreads were prepared as described in Henry-Mowatt et al (10). DNA fiber spreads on acid-treated slides were incubated for 1 hr with rat-anti-BrdU monoclonal antibody for CIdU detection and mouse-anti-BrdU for IdU detection. CIdU and IdU incorporated regions were visualized with Alexa Fluor 555– and Alexa Fluor 488–conjugated secondary antibodies, respectively. Image acquisition was performed using a fluorescent microscope (Zeiss). Lengths and numbers of red or green labeled fibers were measured using ImageJ to quantify replication fork speed and the fraction of individual replication structures.

**Laser micro-irradiation**

Cells were seeded into glass-bottom dishes and allowed to grow for 24 hours. Live cells were imaged with a Nikon A1rsi laser scanning confocal microscope equipped with live-cell incubation chamber maintained at 5% $CO_2$ and 37˚C, using a 20x objective (NA=0.75) at 1024x1024 pixels resolution. For each experiment, a field was selected, and cells were irradiated using a custom micro-irradiation script implemented in NIS-Elements. Briefly, twelve cells were selected for irradiation, and two of these were randomly assigned as controls; control cells did not receive laser irradiation. A pre-irradiation image was collected. For irradiation, a region of interest (ROI) of 3x3 pixels was created within each selected cell nucleus and then the ROI was scanned once with the 405 nm laser at a scan rate of 8 frames per second. Sequential images were then collected every 15 sec for 10 min. Fluorescence intensity at sites of recruitment was then measured using a custom measurement script implemented in FIJI, and intensity data was normalized to maximum recruitment intensity, as we described previously (4,11).

**Immunofluorescence confocal microscopy**

Cells were grown on glass coverslips in 6-well plates. To induce copGFP-Polβ foci formation, cells were treated with various doses of radiation or with cis-Pt (30μM). Fixation was performed in 4% paraformaldehyde (PFA) for 10 min at room temperature, followed by a brief permeabilization using 0.1% TritonX-100 in PBS for 5 min. Cells were then stained with Hoechst to visualize nuclear DNA. Coverslips were mounted onto glass slides with Fluoro-Gel (Electron Microscopy Sciences).

For γ-H2AX, 53BP1, RAD51, XRCC1 and RPA staining, cells cultured on coverslips were treated with radiation or mock-treated and allowed to recover for the indicated times. Cells were then fixed with 4% PFA for 10 min and briefly permeabilized with 0.1% TritonX-100 solution in PBS. Cells were rinsed with PBS and blocked in blocking buffer (1% bovine serum albumin or 10% normal goat serum in PBS) for 30 min and subsequently incubated with the respective primary antibodies for 1 to 2 hrs at 37˚C, followed by three PBS washes and an incubation with fluorescent goat anti-mouse and goat anti-rabbit secondary antibodies (see **Supplementary Table S1** for primary and antibodies used, with dilutions for each). Nuclei were stained with NucBlue (Life Technology) or Hoechst and coverslips were mounted to slides using Prolong Gold or Fluoro-Gel (Thermo Scientific).

Mounted slides were imaged with a Nikon A1rsi laser scanning confocal microscope, using a 100x objective (N.A. 1.45) and 1x confocal zoom controlled by NIS-Elements software. 5-10 image stacks per field were collected, and maximum intensity projections were generated from the collected image stacks. Digital images (5 stacks) were also acquired using a confocal microscope (Leica SP5, Leica Microsystems GmbH), equipped with 63x and 100x objectives. Automated foci analysis were performed in Image J using an in-house (NKI) available macro after optimizing the parameters and as described previously (12). Results from automated counts were validated by visual inspection.

For immunostaining of endogenous TRIP12, 100,000 cells per well were seeded to glass coverslips in a 6-well plate and allowed to grow for 48 h. Media was removed and coverslips were rinsed with cold PBS. Fixation was performed with ice-cold methanol for 10 mins. Cells were then permeabilized using 0.1% Triton X-100 in PBS for 5 min, rinsed with PBS and incubated with blocking buffer (5% NGS, 3% BSA, PBS) for 30 min. Primary antibody to TRIP12 in blocking buffer (1:1000) was added to the cells for 1 hr at room temperature followed by three PBS washes. Secondary fluorescent anti-rabbit antibody (1:2000 diluted in 3% BSA in PBS) was incubated for 30 min at room temperature followed by three PBS washes. Nuclei were stained using NucBlue and coverslips were mounted on glass slides using Prolong Gold anti-fade mounting media.

Ongoing DNA synthesis or cell proliferation markers for foci analysis was detected by treating LN428/copGFP-Polβ cells either with 10μM EDU 10 min before fixation, or with 10μM EDU 10 min before irradiation. This was followed by EDU staining using Click-iT EdU Imaging Systems (Invitrogen, Carlsbad, CA, USA) according to the manufacturer's instructions.

**Immunofluorescence co-localization analysis**

To determine if ionizing radiation (IR) treatment alters the colocalization of Polβ and 53BP1, $2x10^5$ LN428/copGFP-Polβ/Scr, LN428/copGFP-Polβ/TRIP12-sh1 or LN428/copGFP-Polβ/TRIP12-sh5 cells were seeded in a 35mm tissue culture dish with a cover glass bottom (World Precision Instruments, FD35-100) for 24 hrs. Cells without IR treatment or post 10Gy IR treatment for 0, 0.5, 1, and 5 hours were fixed with 4% paraformaldehyde for 15 min, then permeabilized with 0.1% Triton X-100 for 15 min. After washing 3 times with PBS and 5 times with 0.5% BSA in PBS, cells were blocked with 2% bovine serum albumin for 45 min. Mouse anti-γ-H2AX (Sigma, #05-636) antibody at 1:100 dilution and Rabbit anti-53BP1 (Thermo Scientific, #MA5-32653) at 1:200 dilution were incubated with cells for 2 hrs. After washing, Alexa Fluor 647 Goat anti-mouse IgG and Alexa Fluor 568 Goat anti-Rabbit IgG at 1:1000 dilution were cultured with cells for 1 h. After washing, DAPI mounting media was added and a coverslip was adhered to dish. Cells were imaged with a Nikon A1rsi confocal microscope.

Analysis**:** Per-nucleus colocalization of copGFP-Polβ and 53BP1 signals was performed using NIS-Elements by first thresholding to the DAPI signal to identify nuclei, and then designating each individual nucleus a region of interest (ROI). Mander's overlap values for the pixels within each ROI were measured and exported (n=80-150 cells per experimental condition). Statistical analysis was performed by Two-Way ANOVA using GraphPad PRISM.

**Development of LN428/UBR5-KO and LN428/Cas9 cells**

We developed LN428 cell lines with a stable knockout of UBR5 using the one vector CRISPR/Cas9 system (plentiCRISPR-v2; to deliver hSpcas9 and puromycin resistance). The plentiCRISPR-v2 vector was obtained from Addgene (plasmid #52961). To perform the knockout of UBR5, we designed the guide RNA (gRNA) using the ChopChop software package (http://chopchop.cbu.uib.no). The resulting gRNA (UBR5-5’-AGCATTGCTACCTTACGCTGTGG) was predicted to cut at genomic location chr8:102293777 in exon 34 of the UBR5 gene. Control plentiCRISPR-v2 expressing hSpcas9 and the control gRNA (5’-GCGTACCACACCCGTCGCAT) was a gift from Wim Vermeulen (Erasmus MC, The Netherlands). The plasmids were used to generate lentivirus to express either Cas9+CgRNA or Cas9+UBR5-gRNA1 and LN428 cells were transduced as indicated above. Cells were then maintained in media containing puromycin (1µg/ml) for 16 days before protein immunoblot validation of the knockout was performed. Details of the technique have been described by us previously (13) and earlier by Sanjana et al (14).

**Statistical analysis**

Averages and standard deviations (SD) were calculated from the means (on technical replicates) of multiple independent experiments (n = number of independent experiments as indicated in figure legends) unless stated otherwise. ANOVA was used to test for significant differences, generally compared to controls and as indicated in the figure legends. Foci dot plots show the distribution of the foci number per cell values and contain pooled foci data from all independent experiments with a minimum of 50 analyzed cells for each experiment. Some foci data (individual foci number per cell) did not pass the normality test (D’Agostino & Pearson test). In these cases, a non-parametric test (Kruskal-Wallis with Dunn’s multiple comparison test) was used. To further support the conclusions from such analyses, a second analysis was performed that instead shows and compares the means of the individual experiments by calculating the average and standard deviation (SD) of the mean foci number per cell values derived from the independent experiments. These data were used to consider the inter-experimental variation when assessing a possible statistical difference to controls with ANOVA. P-values are indicated by asterisks with *$p<0.05$, **$p<0.01$, ***$p<0.001$, ****$p<0.0001$. The dependence of the data points in the survival dose response data precludes ANOVA analyses. Non-linear regressions were computed using Y=100/(1+(X^HillSlope)/(IC50^HillSlope)) (with Y= Survival in % and X=$H_2O_2$ concentration) on the normalized $H_2O_2$ dose response survival data to calculate $IC_{50}$ (half maximal inhibitory concentration) values for each individual experiment. Similarly, $D_{37}$ (radiation dose permitting 37% cell survival) values were calculated from linear quadratic fits (Y=exp(-α*X-β*X^2) with X=radiation dose and

Y=surviving fraction) on the survival curves. One-way ANOVA was then used to test for significant differences in these response parameter values as stated in the text. Figure legends in the survival curves show the results of the statistical tests for differences in the best-fit parameter ($\alpha$ and $\beta$) with p from extra-sum-of-squares F test (1000 iterations of fits). Statistical analyses were performed using GraphPad PRISM.

#### Supplementary Information

**TRIP12 governs DNA Polymerase β involvement in DNA damage response and repair**

Burcu Inanc, Qingming Fang, Joel Andrews, Xuemei Zeng, Jennifer Clark, Jianfeng Li, Nupur B. Dey, Md Ibrahim, Peter Sykora, Zhongxun Yu, Andrea Braganza, Marcel Verheij, Jos Jonkers, Nathan A. Yates, Conchita Vens and Robert W. Sobol

Supplementary information includes:

Supplementary Figures S1-S5 with corresponding figure legends

### Figure S1

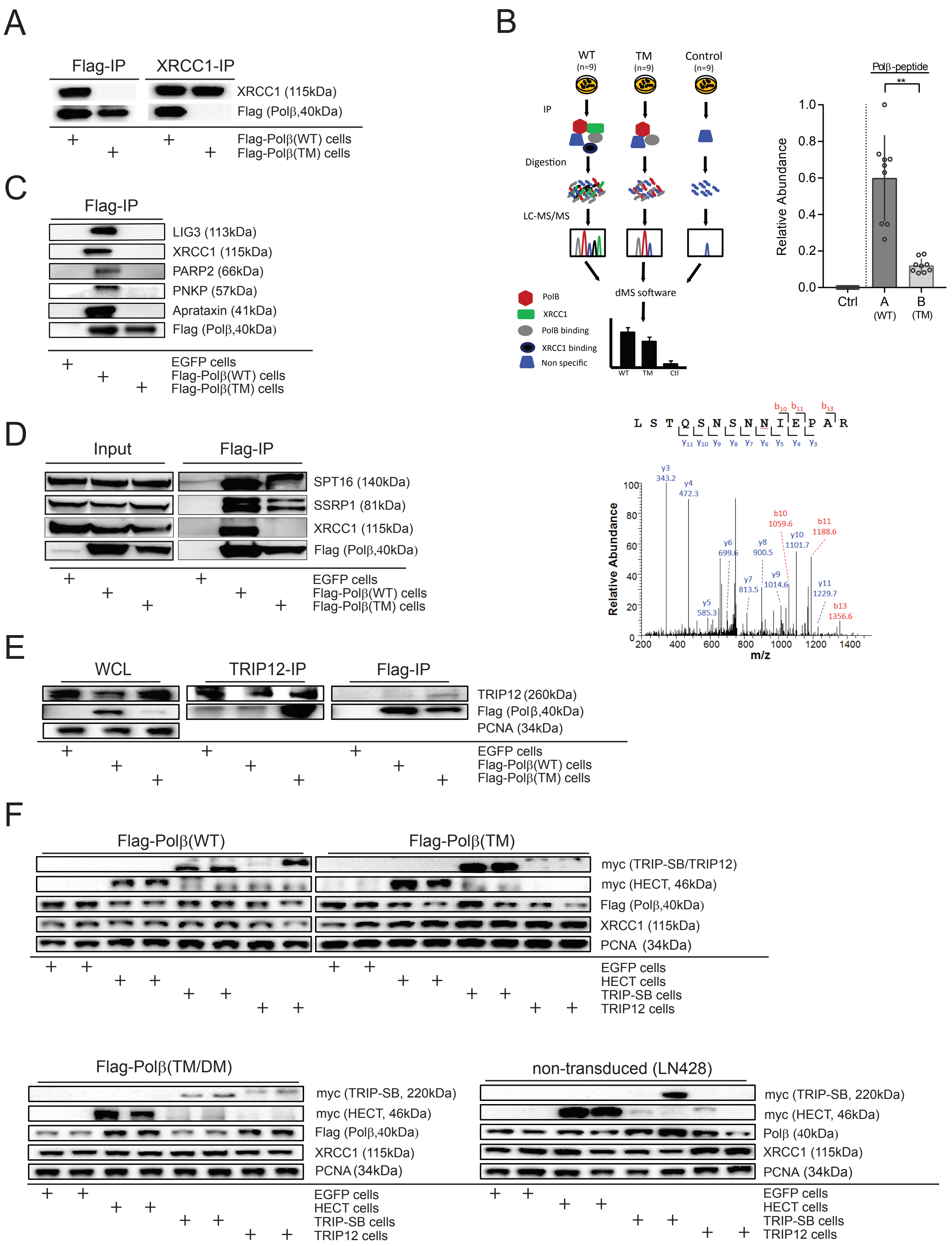

A
Flag-IP
XRCC1-IP
XRCC1 (115kDa)
Flag (Polβ,40kDa)
Flag-Polβ(WT) cells
Flag-Polβ(TM) cells
B
WT (n=9)
TM (n=9)
Control (n=9)
IP
Digestion
LC-MS/MS
dMS software
PolB
XRCC1
PolB binding
XRCC1 binding
Non specific
Polβ-peptide
Relative Abundance
Ctrl
A (WT)
B (TM)
L S T Q S N S N N I E P A R
Relative Abundance
m/z
C
Flag-IP
LIG3 (113kDa)
XRCC1 (115kDa)
PARP2 (66kDa)
PNKP (57kDa)
Aprataxin (41kDa)
Flag (Polβ,40kDa)
EGFP cells
Flag-Polβ(WT) cells
Flag-Polβ(TM) cells
D
Input
Flag-IP
SPT16 (140kDa)
SSRP1 (81kDa)
XRCC1 (115kDa)
Flag (Polβ,40kDa)
EGFP cells
Flag-Polβ(WT) cells
Flag-Polβ(TM) cells
E
WCL
TRIP12-IP
Flag-IP
TRIP12 (260kDa)
Flag (Polβ,40kDa)
PCNA (34kDa)
EGFP cells
Flag-Polβ(WT) cells
Flag-Polβ(TM) cells
F
Flag-Polβ(WT)
Flag-Polβ(TM)
myc (TRIP-SB/TRIP12)
myc (HECT, 46kDa)
Flag (Polβ,40kDa)
XRCC1 (115kDa)
PCNA (34kDa)
EGFP cells
HECT cells
TRIP-SB cells
TRIP12 cells
Flag-Polβ(TM/DM)
non-transduced (LN428)
myc (TRIP-SB, 220kDa)
myc (HECT, 46kDa)
Flag (Polβ,40kDa)
Polβ (40kDa)
XRCC1 (115kDa)
PCNA (34kDa)
EGFP cells
HECT cells
TRIP-SB cells
TRIP12 cells

**Figure S1**

**(A)** Polβ(TM) mutant does not bind to XRCC1. Immunoprecipitation/immunoblot (IP/IB) analysis of transgenic Flag-Polβ (Flag-M2 antibody) and endogenous XRCC1 in LN428 cells expressing either Flag-Polβ(WT) or Flag-Polβ(TM). The anti-Flag or anti-XRCC1 IP complexes (as indicated at the top) were probed by immunoblot for the presence of XRCC1 or Flag-Polβ by anti-XRCC1 and anti-Flag as indicated on the right. **(B)** Label-free differential mass spectrometry (dMS) workflow schema (left), tandem mass spectrum example (bottom) and quantification results (right) for peptide IDEFLATGK from Polβ. Bar graph shows the mean +/- SD and the individual sample results as dots. The abundance values in the immunoprecipitated samples from LN428 cells expressing Flag-Polβ(WT) or Flag-Polβ(TM) were normalized to endogenous Polβ level. **(C)** Independent IB analyses of Flag-Polβ(WT) and Flag-Polβ(TM) immunoprecipitation confirms the lack of the BER proteins Ligase III, XRCC1, PARP2, PNKP and Aprataxin in complex B as generated by immunoprecipitation with Flag-Polβ(TM) and compared to Flag-Polβ(WT). **(D)** Consistent with the dMS data (**Supplementary Table S2**), we find XRCC1-independent but Polβ-mediated interaction with the FACT complex proteins SPT16 and SSRP1. **(E)** Interaction of endogenous TRIP12 with Flag-Polβ(WT) and Flag-Polβ(TM) by IP/IB as in **Figure 1C** in a second independent cell line (T98G)**. (F)** Expression profile of LN428 cells modified by expression of Myc-TRIP12-SB (N-terminal E3 ligase substrate binding domain), HECT (TRIP12 E3 ligase active site HECT domain), Flag-Polβ(WT) and Flag-Polβ(TM) used in **Figures 1** and **S1**. Flag-Polβ proteins were detected by the anti-FLAG antibody, whereas endogenous Polβ was detected by the Polβ monoclonal antibody 61. An anti-myc antibody was used to depict full length myc-TRIP12, myc-TRIP-SB and the myc-HECT domain of TRIP12.

### Figure S2

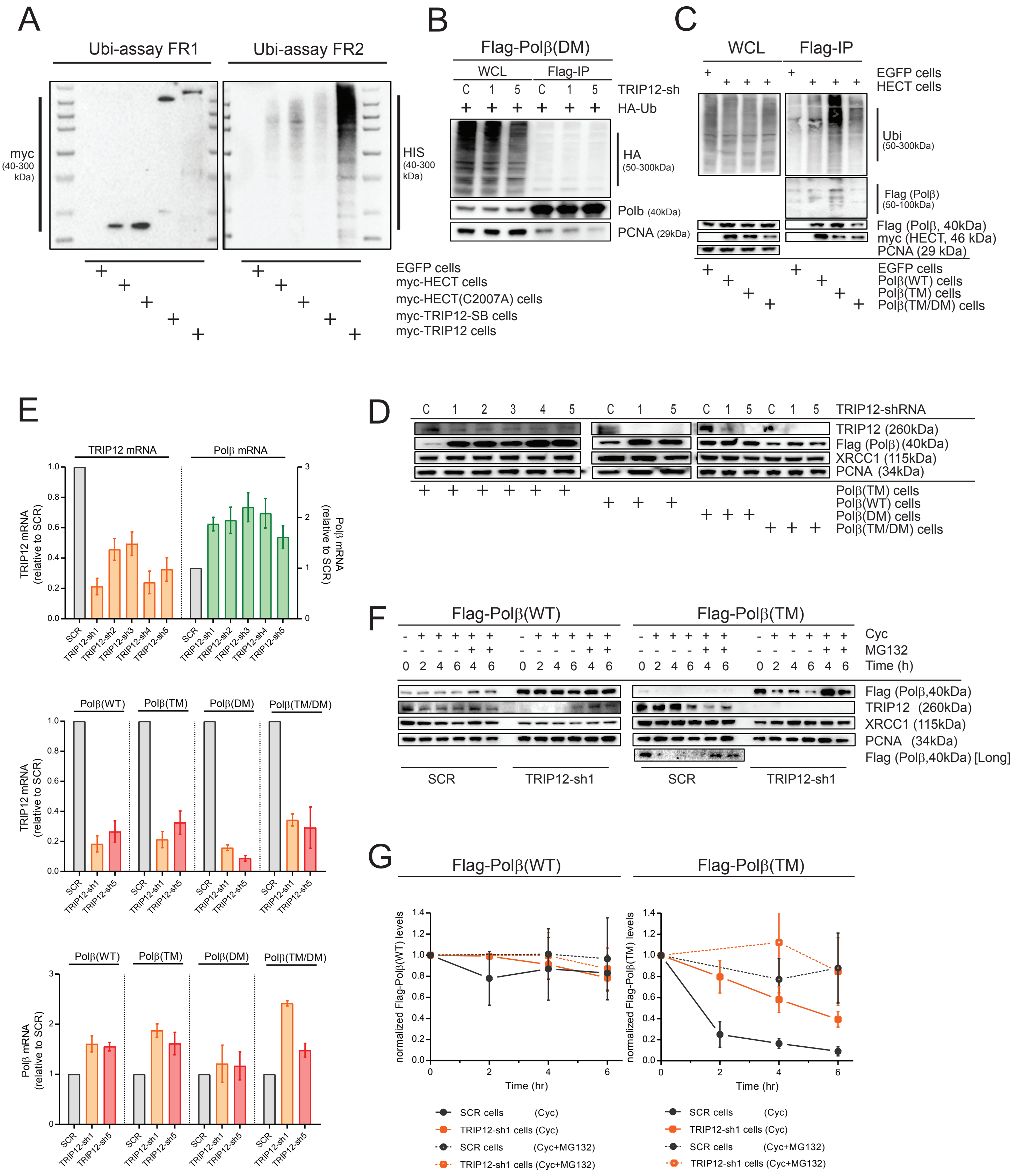

A
Ubi-assay FR1
Ubi-assay FR2
myc (40-300 kDa)
HIS (40-300 kDa)
EGFP cells
myc-HECT cells
myc-HECT(C2007A) cells
myc-TRIP12-SB cells
myc-TRIP12 cells
B
Flag-Polβ(DM)
WCL
Flag-IP
TRIP12-sh
HA-Ub
HA (50-300kDa)
Polb (40kDa)
PCNA (29kDa)
C
WCL
Flag-IP
EGFP cells
HECT cells
Ubi (50-300kDa)
Flag (Polβ) (50-100kDa)
Flag (Polβ, 40kDa)
myc (HECT, 46 kDa)
PCNA (29 kDa)
EGFP cells
Polβ(WT) cells
Polβ(TM) cells
Polβ(TM/DM) cells
D
TRIP12-shRNA
TRIP12 (260kDa)
Flag (Polβ) (40kDa)
XRCC1 (115kDa)
PCNA (34kDa)
Polβ(TM) cells
Polβ(WT) cells
Polβ(DM) cells
Polβ(TM/DM) cells
E
TRIP12 mRNA
Polβ mRNA
TRIP12 mRNA (relative to SCR)
Polβ mRNA (relative to SCR)
SCR
TRIP12-sh1
TRIP12-sh2
TRIP12-sh3
TRIP12-sh4
TRIP12-sh5
Polβ(WT)
Polβ(TM)
Polβ(DM)
Polβ(TM/DM)
F
Flag-Polβ(WT)
Flag-Polβ(TM)
Cyc
MG132
Time (h)
Flag (Polβ,40kDa)
TRIP12 (260kDa)
XRCC1 (115kDa)
PCNA (34kDa)
Flag (Polβ,40kDa) [Long]
SCR
TRIP12-sh1
G
Flag-Polβ(WT)
Flag-Polβ(TM)
normalized Flag-Polβ(WT) levels
normalized Flag-Polβ(TM) levels
Time (hr)
SCR cells (Cyc)
TRIP12-sh1 cells (Cyc)
SCR cells (Cyc+MG132)
TRIP12-sh1 cells (Cyc+MG132)

**Figure S2**

**(A)** Capture of myc-fusion proteins (myc-HECT, myc-HECT(C2007A), myc-TRIP12-SB and myc-TRIP12) by MYC-Trap agarose beads (left) and ubiquitylation reaction products in the absence of Polβ (right). Shown is the immunoblot of the bead-captured fusion proteins from fractions 1 (FR1) and 2 (FR2) of the Ubi-assay as indicated in Figure 2A probed with the anti-myc Ab (left) for FR1 and with the anti-His Ab (right) after incubation with His-Ubiquitin and the E1/E2 reaction cocktail for FR2 . **(B)** Lack of ubiquitylation in transgenic K206A/K244A mutant Polβ(DM) as determined by IP/IB following transfection of HA-ubiquitin in LN428 cells as in **Figure 2C**. **(C)** TRIP12-HECT domain binding to ubiquitylated Polβ. Myc-HECT shows preferential binding to the ubiquitylated form of Polβ as determined by IP/IB in LN428 cells expressing myc-HECT and Flag-Polβ(WT), Flag-Polβ(TM) or Flag-Polβ(TM/DM), followed by detection with an anti-Flag and anti-ubiquitin antibodies as specified in Materials and Methods (STAR Methods). **(D)** TRIP12 depletion (knockdown by shRNA) reduces TRIP12 protein levels and stabilizes Polβ(TM) and Polβ(WT) but not the Polβ(DM) variant. Representative immunoblots of cell lysates following TRIP12 depletion by shRNAs (#1 to 5) are shown for TRIP12, Polβ, PARP1, XRCC1 and PCNA in support of **Figure 2D**. **(E)** Polβ wildtype or mutant mRNA levels are not reduced by TRIP12 knock down. The mRNA levels of TRIP12 and the indicated Flag-Polβ variants as determined by qRT-PCR **(**n=3, SD) following TRIP12 depletion via shRNA in LN428 cells, corresponding to **Figure S2C** and **Figure 2D**. **(F-G)** TRIP12 knockdown rescues Polβ(TM) degradation**.** Representative immunoblots and Flag-Polβ(WT) or Flag-Polβ(TM) quantifications in cell lysates prepared from TRIP12-depleted (TRIP12-sh1) and scrambled shRNA (SCR) transduced LN428 cells, treated with the protein synthesis inhibitor cyclohexamide (Cyc) and the proteasome inhibitor MG132 for times periods as indicated: Flag-Polβ(WT) (left) and Flag-Polβ(TM) expressing cells (right). Levels of TRIP12, XRCC1 and PCNA are shown as indicated and are the mean of n=3 with SD.

### Figure S3

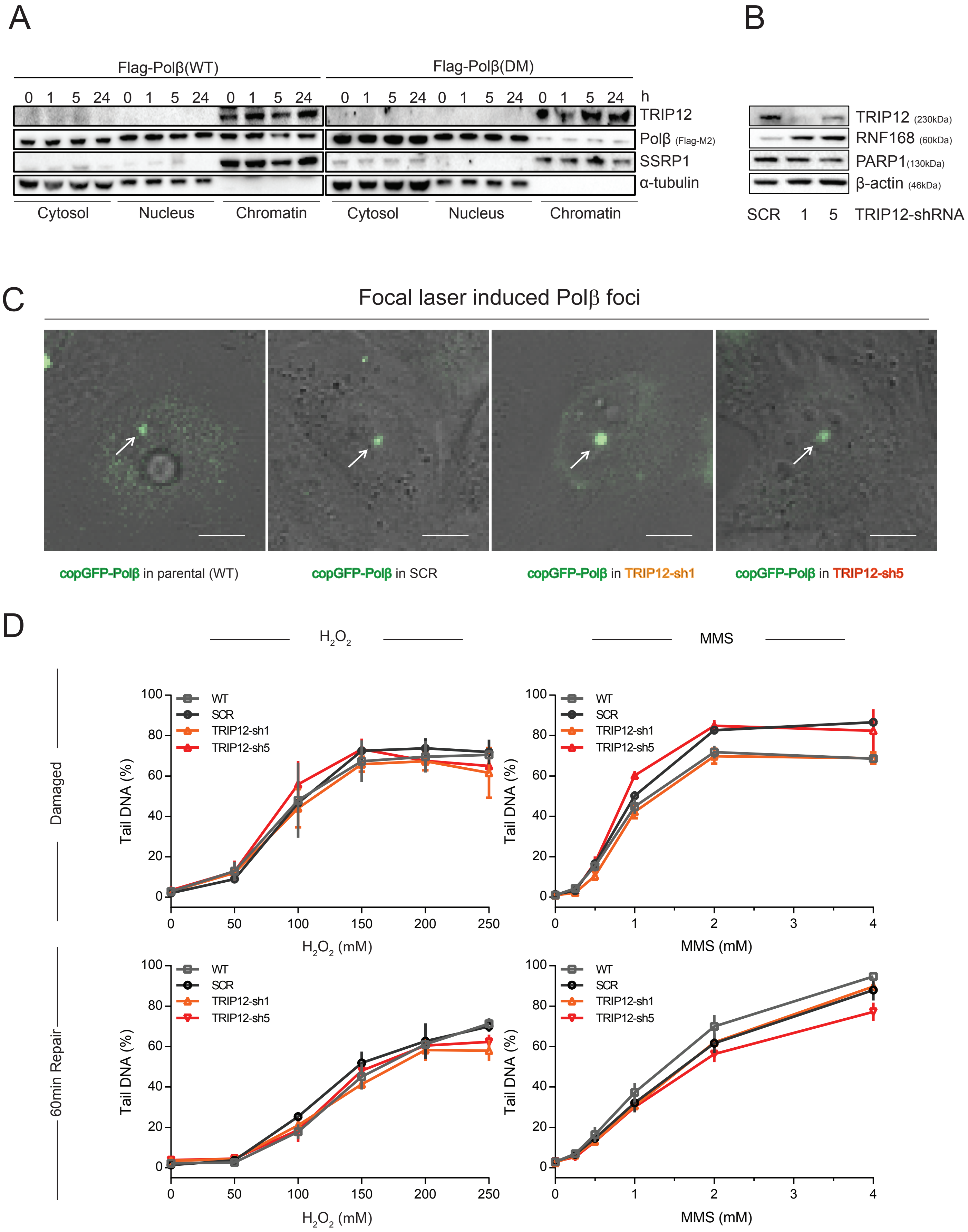
A
Flag-Polβ(WT)
Flag-Polβ(DM)
0 1 5 24 0 1 5 24 0 1 5 24 0 1 5 24 0 1 5 24 0 1 5 24 h
TRIP12
Polβ (Flag-M2)
SSRP1
α-tubulin
Cytosol
Nucleus
Chromatin
Cytosol
Nucleus
Chromatin
B
TRIP12 (230kDa)
RNF168 (60kDa)
PARP1(130kDa)
β-actin (46kDa)
SCR 1 5 TRIP12-shRNA
C
Focal laser induced Polβ foci
copGFP-Polβ in parental (WT)
copGFP-Polβ in SCR
copGFP-Polβ in TRIP12-sh1
copGFP-Polβ in TRIP12-sh5
D
H2O2
MMS
Damaged
60min Repair
Tail DNA (%)
WT
SCR
TRIP12-sh1
TRIP12-sh5
H2O2 (mM)
MMS (mM)

**Figure S3**

**(A)** Mutation of Polβ ubiquitylation sites (K206A/K244A = DM) alters sub-cellular distribution and shows a reduced chromatin association of transgenic Flag-Polβ. Immunoblots show TRIP12, Flag-Polβ, tubulin (cytosol/soluble nuclear fraction loading control) and SSRP1 (chromatin fraction loading control) levels in the cytosolic, nucleoplasmic or chromatin fraction in LN428 cells expressing either transgenic Flag-Polβ(WT) (left) or Flag-Polβ(DM) (right), as indicated. **(B)** TRIP12-mediated change in total cellular levels of RNF168. Immunoblots show RNF168, PARP1 and β-actin (loading control) levels in whole cell lysates from TRIP12 depleted cell lines (TRIP12-sh1 and -sh5) compared to the scrambled control (SCR), as indicated. **(C)** Representative images of laser (405nm) damage-induced Polβ recruitment and retention in TRIP12-KD cells (TRIP12-sh1, TRIP12-sh5), scrambled (SCR) cells and parental (WT) LN428 control cells. **(D)** Oxidative damage and repair as determined by alkaline CometChip analysis (Tail DNA in %) following a 30 min exposure to varying concentrations of $H_2O_2$ (damaged) and at 60 min post exposure (repair) in TRIP12-KD (sh1 and sh5), scrambled (SCR) or parental (WT) control cell lines. The variation in the effect size of the TRIP12-sh transduction is within the response variation of the controls resulting in non-significant alterations when combined.

### Figure S4A-B

## A

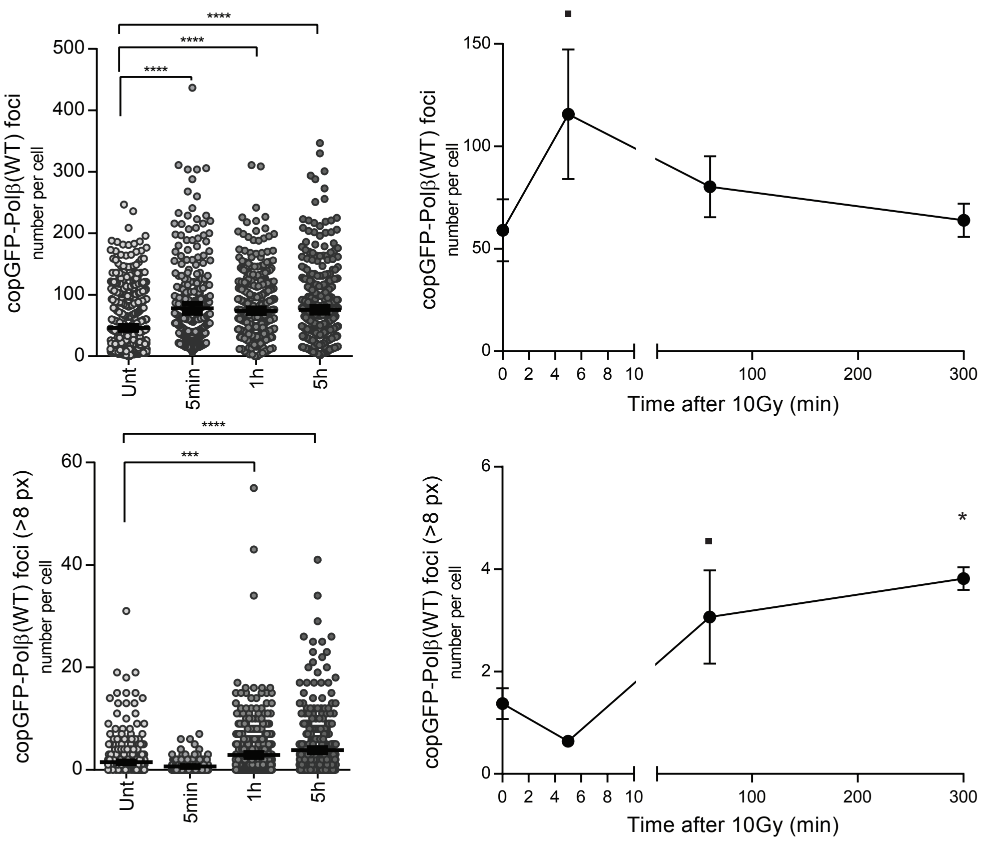

## B

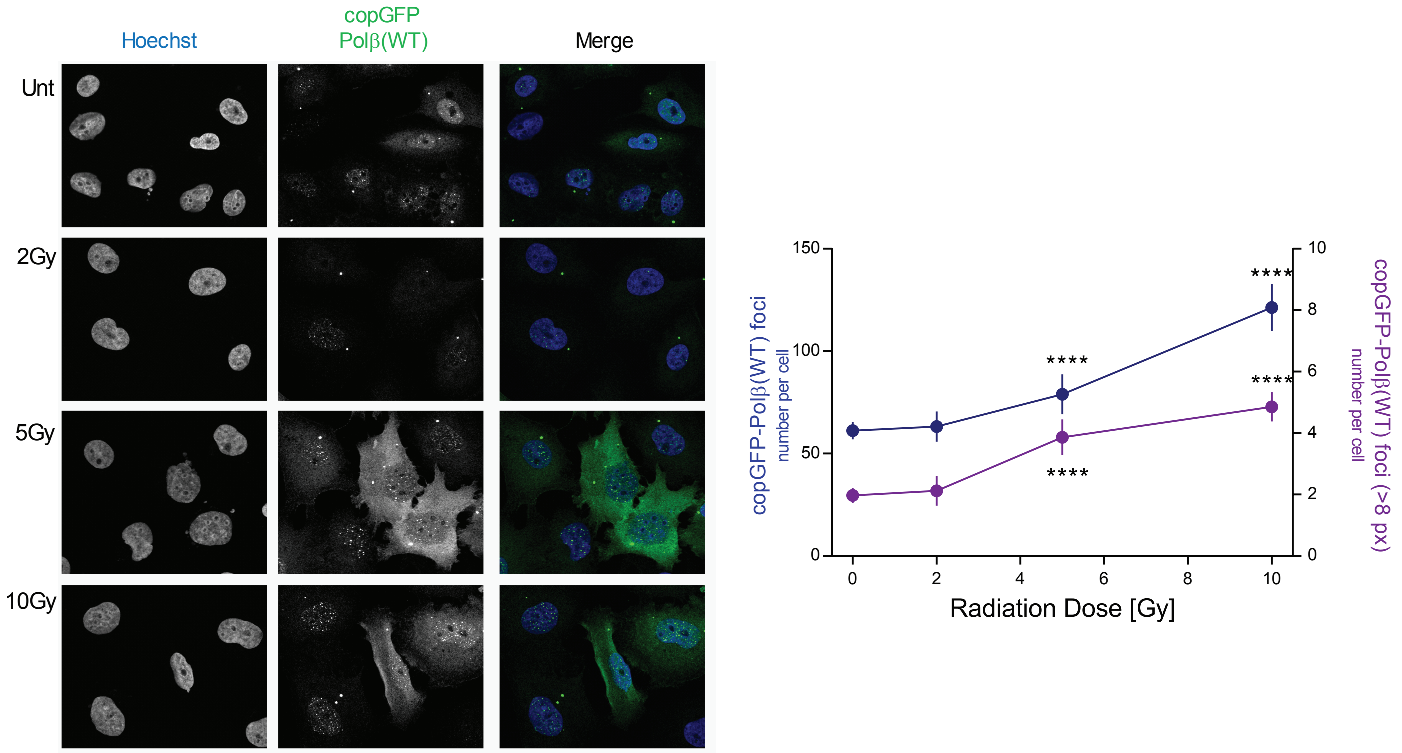

### Figure S4C

C

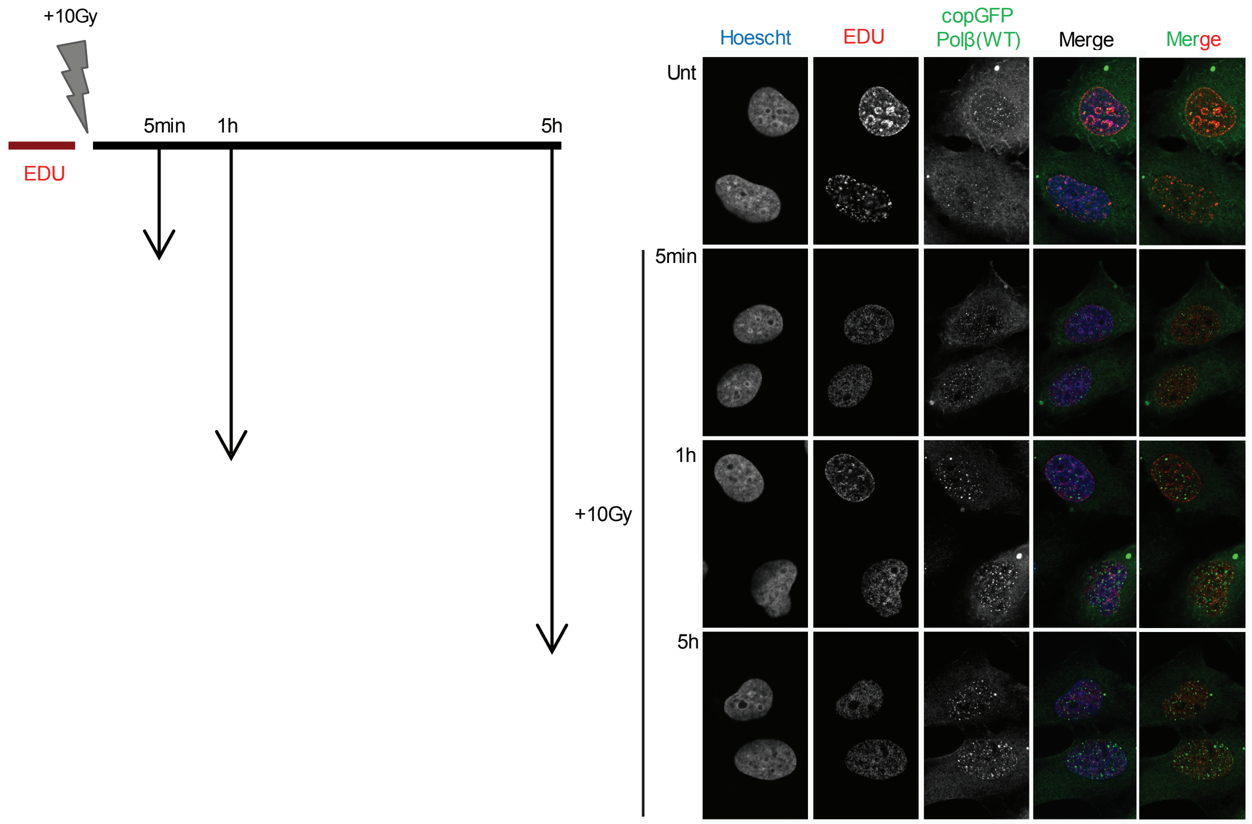
+10Gy
5min
1h
5h
EDU
Hoescht
EDU
copGFP
Polβ(WT)
Merge
Merge
Unt
5min
1h
5h
+10Gy

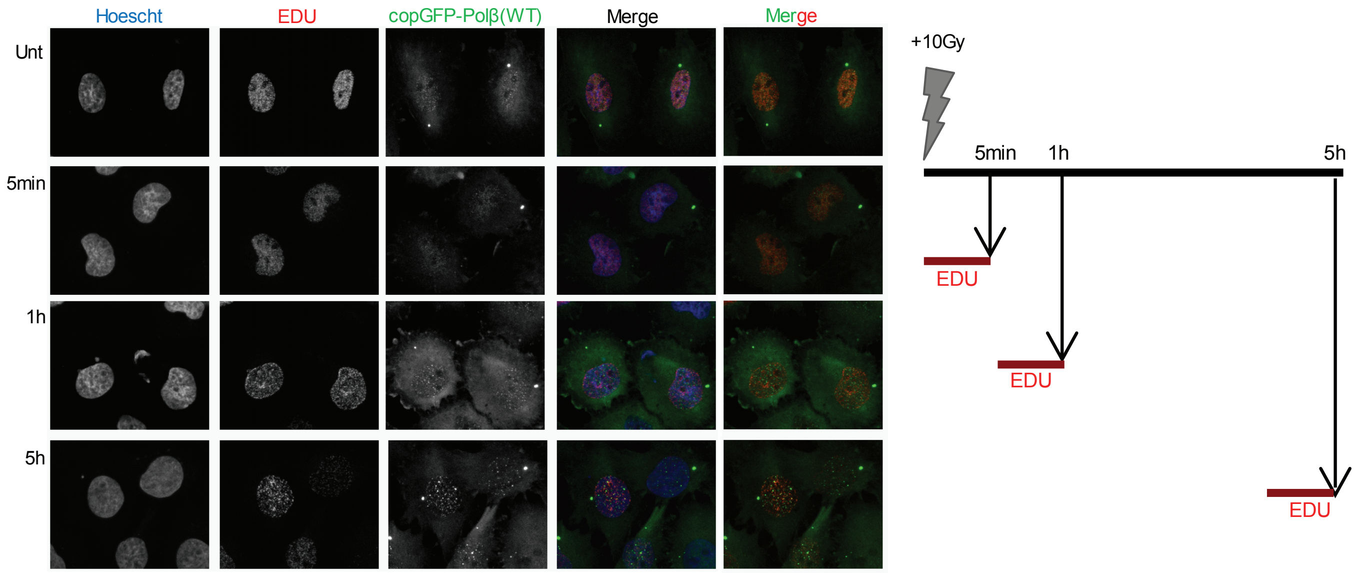
Hoescht
EDU
copGFP-Polβ(WT)
Merge
Merge
Unt
5min
1h
5h
+10Gy
5min
1h
5h
EDU
EDU
EDU

### Figure S4DEF

D

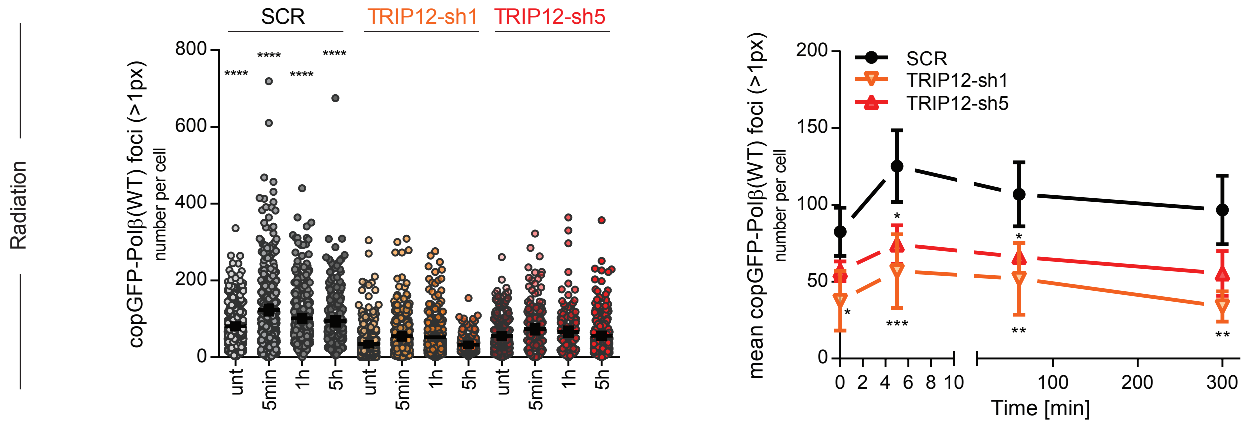

E

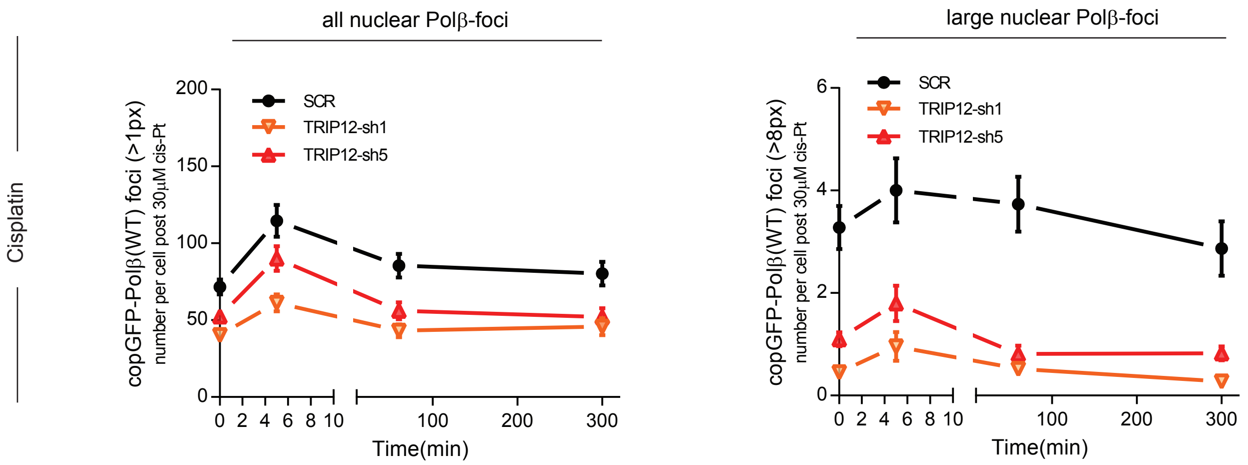

F

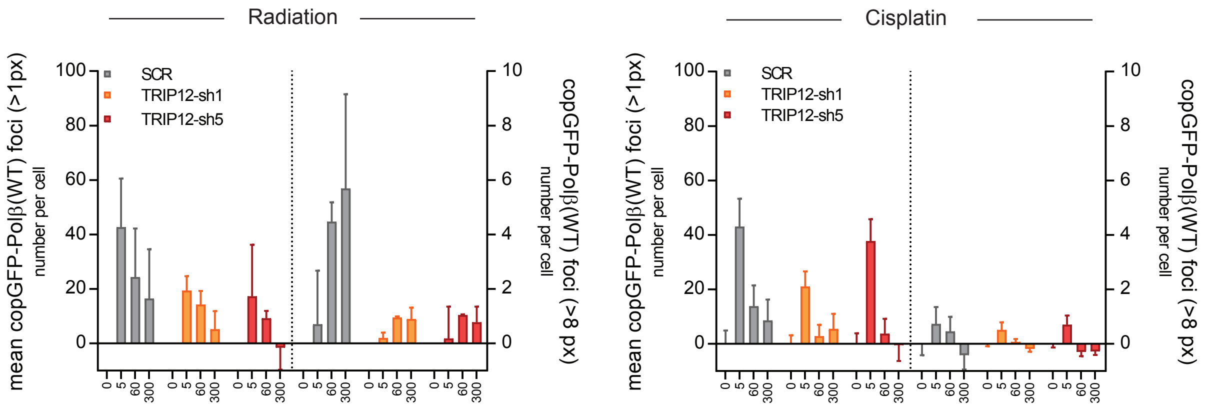

**Figure S4**

**(A)** Characterization of radiation-induced Polβ foci over time. Graphs to the left show the copGFP-Polβ foci distribution of all (>1px, upper panel) or large (>8px, lower panel) Polβ foci in untreated cells (Unt) and in cells at 5min, 1h or 5h after 10Gy (n=3 independent experiments with >50 cells each). The averages of the mean foci count values for these experiments with SD are shown in the plots to the right. At early time points foci are small while larger foci accumulate with time (lower panel). ▪ p<0.06, * p<0.05, *** p<0.001 and **** p<0.05 in the Kruskal Wallis (left) and ANOVA tests (right). **(B)** Polβ foci formation is radiation dose dependent and correlates with the number of radiation-induced lesions. Representative images (left) and quantification results showing the average copGFP-Polβ foci counts per cell of all foci (left Y axis and blue values) and large foci (>8px, right Y axis with purple values) at 1h after radiation. Data show mean and SD of pooled foci counts of 2-3 independent experiments and **** indicate multiple comparison adjusted p<0.001 to the respective un-irradiated controls (ANOVA). **(C)** After radiation, copGFP-Polβ is excluded from sites of active DNA synthesis, labelled with EDU for 10 min before radiation (upper panel). Extensive DNA synthesis is not visible at copGFP-Polβ foci sites after 10Gy radiation, as determined by 10 min EDU exposure prior to fixation at the indicated times (bottom panel). **(D)** TRIP12 also promotes small Polβ foci formation in radiated cells. copGFP-Polβ foci (all) in TRIP12-depleted cells (TRIP12-sh1 and -sh5) and scrambled shRNA control cells (SCR) at indicated time points after 10Gy. Foci distribution on pooled foci data (left graph) and experimental averages and variation (right graph) with the average +/- SD of the mean foci counts in n>3 independent experiments with n>50 cells. Asterisks indicate significant differences to controls (SCR) at each time point with * p<0.05, ** p<0.01 and *** p<0.001 (ANOVA). Radiation-induced foci are significantly different from untreated in the SCR 5 min data point only with * p<0.05. **(E)** All (left) and large (right) copGFP-Polβ foci formation after cisplatin (cis-Pt) exposure in TRIP12-depleted and scrambled (SCR) control cells at the indicated times. The mean and SD of copGFP-Polβ foci counts after 30µM cisplatin treatment at indicated time points are shown. **(F)** Comparison of Polβ foci after background removal to demonstrate number and TRIP12 dependence of damage-induced foci after radiation and cisplatin. While TRIP12 does not affect cisplatin induced foci, neither small nor large, radiation induced foci are impacted by TRIP12 status, in particular the larger and late appearing foci as indicated in the >8px values.

### Figure S5

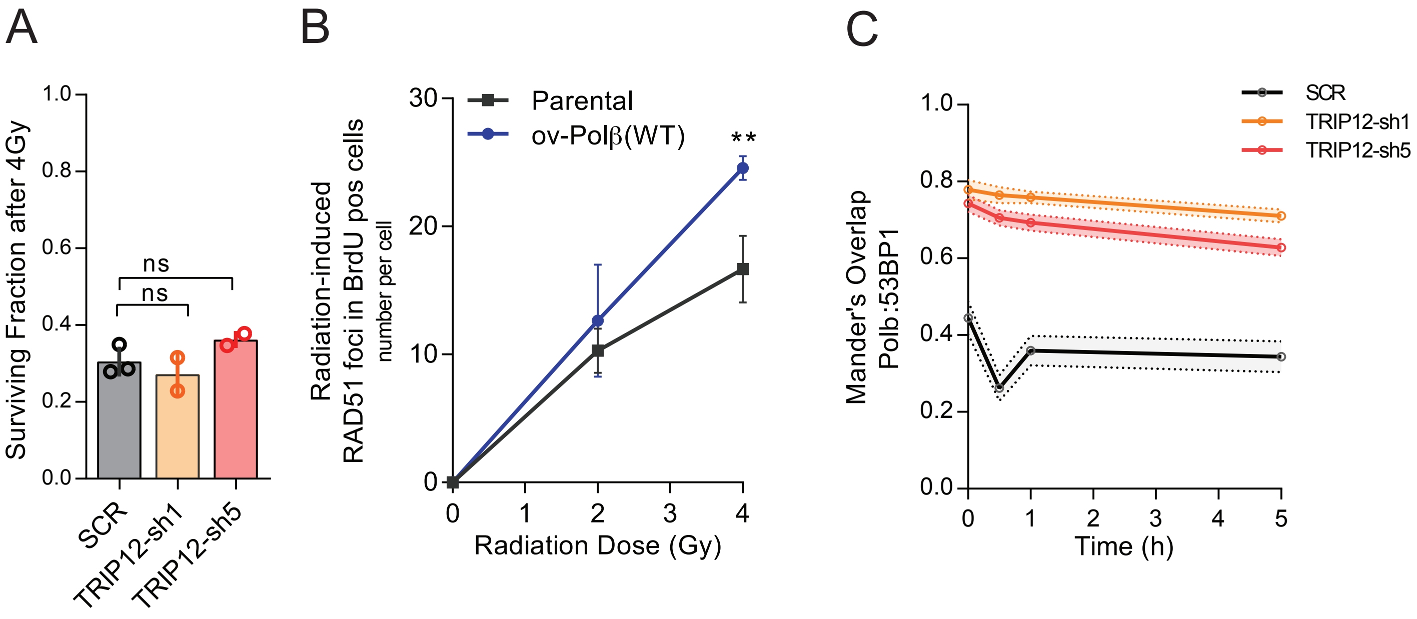
A
B
C
Surviving Fraction after 4Gy
1.0
0.8
0.6
0.4
0.2
0.0
ns
ns
SCR
TRIP12-sh1
TRIP12-sh5
Radiation-induced
RAD51 foci in BrdU pos cells
number per cell
30
20
10
0
Parental
ov-Polβ(WT)
**
0
1
2
3
4
Radiation Dose (Gy)
Mander's Overlap
Polb:53BP1
SCR
TRIP12-sh1
TRIP12-sh5
1.0
0.8
0.6
0.4
0.2
0.0
0
1
2
3
4
5
Time (h)

**Figure S5**

**(A)** Radiation response is unaltered by TRIP12 depletion. TRIP12-depleted cells (TRIP12-sh1 and -sh5) were compared to scrambled (SCR) LN428 control cells. Clonogenic survival after 4Gy with n=2-3 independent experiments; errors are SD, ANOVA test. **(B)** Dose response behavior of increased RAD51 foci formation in Polβ(WT) over-expressing LN428 cells at 5h. Foci numbers in replicating, BrdU positive cells, were counted. Values are the average with SD of the means of 3 independent experiments (** = $p<0.01$, ANOVA). **(C)** Time course graph of data in Figure 7A. Mander's overlap coefficients with mean and 95% CI at different times (h) after radiation (10Gy), illustrating an early TRIP12-controlled radiation response.

### Supplementary Table S1

##### to "TRIP12 governs DNA Polymerase β involvement in DNA damage response and repair"

Burcu Inanc, Qingming Fang, Joel F. Andrews, Xuemei Zeng, Jennifer Clark, Jianfeng Li, Nupur B. Dey, Md Ibrahim, Peter Sykora, Zhongxun Yu, Andrea Braganza, Marcel Verheij, Jos Jonkers, Nathan A. Yates, Conchita Vens, and Robert W. Sobol

#### A. Cell line models

| Cell line name | Characteristics / transgene expression | ID# vector | parental cell line | reference/source | culture medium |
|---|---|---|---|---|---|
| LN428 | human glioblastoma tumor cell line | | LN428 | (2) | #1 |
| LN428/Flag-Polb | Flag-Polb(WT); WT = wildtype | | LN428 | (3) | # 2 |
| LN428/Flag-Polb(TM) | Flag-Polb(TM); TM = L301R/V303R/V306R) | | LN428 | (3) | # 2 |
| LN428/Flag-Polb(DM) | Flag-Polb(DM); DM = K206A/K244A | | LN428 | (3) | # 2 |
| LN428/Myc-HECT | TRIP12 HECT domain (AA 1651-2040) with an N-terminal Myc-tag | | LN428 | * | # 2 |
| LN428/Myc-HECT(C2007) | TRIP12 HECT(C2007A) domain (AA 1651-2040) with an N-terminal Myc-tag | | LN428 | * | # 2 |
| LN428/Myc-TRIP-SB | TRIP12 substrate binding (SB) domain (AA 1-1650) of TRIP12 with an N-terminal Myc-tag | | LN428 | * | # 2 |
| LN428/Myc-TRIP12 | TRIP12 with an N-terminal Myc-tag | | LN428 | * | # 2 |
| LN428/SCR | scrambled shRNA | | LN428 | * | # 2 |
| LN428/TRIP12-sh1 | TRIP12-shRNA#1 | | LN428 | * | # 2 |
| LN428/TRIP12-sh2 | TRIP12-shRNA#2 | | LN428 | * | # 2 |
| LN428/TRIP12-sh5 | TRIP12-shRNA#5 | | LN428 | * | # 2 |
| LN428/Flag-Polb(WT)/SCR | Flag-Polb(WT) and scrambled shRNA | | LN428 | * | # 3 |
| LN428/Flag-Polb(WT)/TRIP12-sh1 | Flag-Polb(WT) and TRIP12-shRNA#1 | | LN428 | * | # 3 |
| LN428/Flag-Polb(WT)/TRIP12-sh5 | Flag-Polb(WT) and TRIP12-shRNA#5 | | LN428 | * | # 3 |
| LN428/Flag-Polb(TM)-SCR | Flag-Polb(TM) and scrambled shRNA; TM = L301R/V303R/V306R | | LN428 | * | # 3 |
| LN428/Flag-Polb(TM)-TRIP12-sh1 | Flag-Polb(TM) and TRIP12-shRNA#1; TM = L301R/V303R/V306R | | LN428 | * | # 3 |
| LN428/Flag-Polb(TM)-TRIP12-sh5 | Flag-Polb(TM) and TRIP12-shRNA#5; TM = L301R/V303R/V306R | | LN428 | * | # 3 |
| LN428/copGFP-Polb(WT) | copGFP-Polb(WT) fusion protein | | LN428 | (3) | # 2 |
| LN428/copGFP-Polb(TM) | copGFP-Polb(TM) fusion protein; TM = L301R/V303R/V306R | | LN428 | (3) | # 2 |
| LN428/copGFP-Polb(DM) | copGFP-Polb(DM) fusion protein; DM = K206A/K244A | | LN428 | * | # 2 |
| LN428/copGFP-Polb(WT)/SCR | copGFP-Polb(WT) fusion protein and scrambled shRNA | | LN428 | * | # 2 |
| LN428/copGFP-Polb(WT)/TRIP12-sh1 | copGFP-Polb(WT) fusion protein and TRIP12-shRNA#1 | | LN428 | * | # 2 |
| LN428/copGFP-Polb(WT)/TRIP12-sh5 | copGFP-Polb(WT) fusion protein and TRIP12-shRNA#5 | | LN428 | * | # 2 |
| LN428/copGFP-Polb(DM)/SCR | copGFP-Polb(DM) fusion protein and scrambled shRNA; DM = K206A/K244A | | LN428 | * | # 2 |
| LN428/copGFP-Polb(DM)/TRIP12-sh1 | copGFP-Polb(DM) fusion protein and TRIP12-shRNA#1; DM = K206A/K244A | | LN428 | * | # 2 |
| LN428/copGFP-Polb(DM)/TRIP12-sh5 | copGFP-Polb(DM) fusion protein and TRIP12-shRNA#5; DM = K206A/K244A | | LN428 | * | # 2 |
| LN428/copGFP-Polb(TM)/SCR | copGFP-Polb(TM) fusion protein and scrambled shRNA; TM = L301R/V303R/V306R | | LN428 | * | # 2 |
| LN428/copGFP-Polb(TM)/TRIP12-sh1 | copGFP-Polb(TM) fusion protein and TRIP12-shRNA#1; TM = L301R/V303R/V306R | | LN428 | * | # 2 |
| LN428/copGFP-Polb(TM)/TRIP12-sh5 | copGFP-Polb(TM) fusion protein and TRIP12-shRNA#5; TM = L301R/V303R/V306R | | LN428 | * | # 2 |
| LN428/Flag-Polb(WT)/EGFP | Flag-Polb(WT) and EGFP | | LN428 | * | # 4 |
| LN428/Flag-Polb(WT)/Myc-HECT | Flag-Polb(WT) and the HECT domain (AA 1651-2040) of TRIP12 with an N-terminal Myc-tag | | LN428 | * | # 4 |

| | | | | | |
|---|---|---|---|---|---|
| LN428/Flag-Polb(WT)/Myc-TRIP12-SB | Flag-Polb(WT) and the substrate binding (SB) domain (AA 1-1650) of TRIP12 with an N-terminal Myc-tag | | LN428 | * | # 4 |
| LN428/Flag-Polb(WT)/Myc-TRIP12 | Flag-Polb(WT) and TRIP12 with an N-terminal Myc-tag | | LN428 | * | # 4 |
| LN428/Flag-Polb(TM)/EGFP | Flag-Polb(TM) and EGFP; TM = L301R/V303R/V306R | | LN428 | * | # 4 |
| LN428/Flag-Polb(TM)/Myc-HECT | Flag-Polb(TM) and the TRIP12 HECT domain (AA 1651-2040) with an N-terminal Myc-tag; TM = L301R/V303R/V306R | | LN428 | * | # 4 |
| LN428/Flag-Polb(TM)/Myc-TRIP12-SB | Flag-Polb(TM) and the TRIP12 substrate binding (SB) domain (AA 1-1650) with an N-terminal Myc-tag; TM = L301R/V303R/V306R | | LN428 | * | # 4 |
| LN428/Flag-Polb(TM)/Myc-TRIP12 | Flag-Polb(TM) and TRIP12 with an N-terminal Myc-tag; TM = L301R/V303R/V306R | | LN428 | * | # 4 |
| LN428/Flag-Polb(TM/DM)/EGFP | Flag-Polb(TM/DM) and EGFP; TM/DM = L301R/V303R/V306R/K206A/K244A | | LN428 | * | # 4 |
| LN428/Flag-Polb(TM/DM)/Myc-HECT | Flag-Polb(TM/DM) and the TRIP12 HECT domain (AA 1651-2040) with an N-terminal Myc-tag; TM/DM = L301R/V303R/V306R/K206A/K244A | | LN428 | * | # 4 |
| LN428/Flag-Polb(TM/DM)/Myc-TRIP12-SB | Flag-Polb(TM/DM) and the TRIP12 substrate binding (SB) domain (AA 1-1650) with an N-terminal Myc-tag; TM/DM = L301R/V303R/V306R/K206A/K244A | | LN428 | * | # 4 |
| LN428/Flag-Polb(TM/DM)/Myc-TRIP12 | Flag-Polb(TM/DM) and TRIP12 with an N-terminal Myc-tag; TM/DM = L301R/V303R/V306R/K206A/K244A) | | LN428 | * | # 4 |
| LN428/Myc-HECT/EGFP | TRIP12 HECT domain (AA 1651-2040) with an N-terminal Myc-tag and EGFP | | LN428 | * | # 4 |
| LN428/Myc-HECT/Flag-Polb-C | TRIP12 HECT domain (AA 1651-2040) with an N-terminal Myc-tag and Flag-Polb(WT), C-terminal domain (AA 91-335) | | LN428 | * | # 4 |
| LN428/Myc-HECT/Flag-Polb(TM)-C | TRIP12 HECT domain (AA 1651-2040) with an N-terminal Myc-tag and Flag-Polb(TM), C-terminal domain (AA 91-335); TM = L301R/V303R/V306R | | LN428 | * | # 4 |
| LN428/Myc-HECT/Flag-Polb(TM/DM)-C | TRIP12 HECT domain (AA 1651-2040) with an N-terminal Myc-tag and Flag-Polb(TM/DM), C-terminal domain (AA 91-335); TM/DM= L301R/V303R/V306R/K206A/K244A | | LN428 | * | # 4 |
| LN428/Myc-TRIP12-SB/EGFP | TRIP12 substrate binding (SB) domain (AA 1-1650) with an N-terminal Myc-tag and EGFP | | LN428 | * | # 4 |
| LN428/Myc-TRIP12-SB/Flag-Polb-C | TRIP12 SB domain (AA 1-1650) with an N-terminal Myc-tag and Flag-Polb(WT), C-terminal domain (AA 91-335) | | LN428 | * | # 4 |
| LN428/Myc-TRIP12-SB/Flag-Polb(TM)-C | TRIP12 SB domain (AA 1-1650) with an N-terminal Myc-tag and Flag-Polb(TM), C-terminal domain (AA 91-335); TM = L301R/V303R/V306R | | LN428 | * | # 4 |
| LN428/Myc-TRIP12-SB/Flag-Polb(TM/DM)-C | TRIP12 SB domain (AA 1-1650) with an N-terminal Myc-tag and Flag-Polb(TM/DM), C-terminal domain (AA 91-335); TM/DM = L301R/V303R/V306R/K206A/K244A | | LN428 | * | # 4 |
| LN428/Cas9 | Cas9 and a control, non-targeting gRNA | | LN428 | * | # 4 |
| LN428/UBR5-KO.1A | Cas9 and UBR5-specific gRNA1; cell line A | | LN428 | * | # 4 |
| LN428/UBR5-KO.1B | Cas9 and UBR5-specific gRNA1; cell line B | | LN428 | * | # 4 |
| HCT116 | human colon cancer cell line | | HCT116 | (4) | # 5 |
| HCT116/SCR | scrambled shRNA | | HCT116 | * | # 6 |
| HCT116/TRIP12-sh1 | TRIP12-shRNA#1 | | HCT116 | * | # 6 |
| HCT116/TRIP12-sh2 | TRIP12-shRNA#2 | | HCT116 | * | # 6 |
| HCT116/TRIP12-sh5 | TRIP12-shRNA#5 | | HCT116 | * | # 6 |
| T98G | human glioblastoma cell line | | T98G | (5) | # 7 |
| T98G/Flag-Polb | Flag-Polb(WT); WT= wildtype | | T98G | (3) | # 8 |
| T98G/Flag-Polb(TM) | Flag-Polb(TM); TM = L301R/V303R/V306R | | T98G | (3) | # 8 |
| 293-FT | human embryonal kidney cell line transformed with SV40 large T antigen for lentiviral production | | 293-FT | (6) | # 9 |

* this study

(1) Braganza, A., Li, J., Zeng, X., Yates, N.A., Dey, N.B., Andrews, J., Clark, J., Zamani, L., Wang, X.H., St Croix, C. et al. (2017) UBE3B Is a Calmodulin-regulated, Mitochondrion-associated E3 Ubiquitin Ligase. J Biol Chem, 292, 2470-2484.
(2) Tang, J.B., Svilar, D., Trivedi, R.N., Wang, X.H., Goellner, E.M., Moore, B., Hamilton, R.L., Banze, L.A., Brown, A.R. and Sobol, R.W. (2011) N-methylpurine DNA glycosylase and DNA polymerase beta modulate BER inhibitor potentiation of glioma cells to temozolomide. Neuro-oncology, 13, 471-486.
(3) Fang, Q., Inanc, B., Schamus, S., Wang, X.H., Wei, L., Brown, A.R., Svilar, D., Sugrue, K.F., Goellner, E.M., Zeng, X. et al. (2014) HSP90 regulates DNA repair via the interaction between XRCC1 and DNA polymerase beta. Nature communications, 5, 5513.
(4) Dr. B Vogelstein (J. Hopkins)
(5) ATCC – Cat# CRL1690
(6) Thermo Fisher Scientific – Cat# R70007

Media #1: α-MEM (Thermo Fisher Scientific) with 10% heat inactivated FBS (Sigma-Aldrich), 5g/ml Gentamycin, 80u Penicillin/80µg Streptomycin /0.32µg Amphotericin per ml, 2mM L-Glutamine (all Thermo Fisher Scientific)
Media #2: Media #1 supplemented with Puromycin (1.0 µg/ml) (Sigma-Aldrich)
Media #3: Media #1 supplemented with Puromycin (1.0 µg/ml) (Sigma-Aldrich) and Geneticin (0.6 mg/ml) (Corning)
Media #4: Media #1 supplemented with Puromycin (1.0 µg/ml) (Sigma-Aldrich), and Hygromycin B (200µg/ml) (Sigma-Aldrich)
Media #5: McCoy's 5A medium (Thermo Fisher Scientific) with 10% heat inactivated FBA, 80u Penicillin/80µg Streptomycin per ml
Media #6: McCoy's 5A medium (Thermo Fisher Scientific) with 10% heat inactivated FBA, 80u Penicillin/80µg Streptomycin per ml, supplemented with Puromycin (0.5 µg/ml)
Media #7: MEM with 10% heat inactivated FBS, 5mg/ml Gentamycin, 80u Penicillin/80g Streptomycin/0.32mg Amphotericin per ml, 1mM Sodium Pyruvate, 10mM MEM non-essential amino acids
Media #8: MEM with 10% heat inactivated FBS, 5mg/ml Gentamycin, 80u Penicillin/80g Streptomycin/0.32mg Amphotericin per ml, 1mM Sodium Pyruvate, 10mM MEM non-essential amino acids, supplemented with Puromycin (1.0 µg/ml)
Media #9: DMEM medium (Thermo Fisher Scientific) with 10% heat inactivated FBS, 80u Penicillin/80µg Streptomycin per ml, 2mM glutamax

#### B. Vectors and constructs

| # | Vector name | insert | ID# construct / vector | reference/source |
|---|---|---|---|---|
| V1 | pRS1436 | Flag-Polb N-terminal fragment (AA 1-90) | SLS130 | (4) |
| V2 | pRS1427 | Flag-Polb C-terminal fragment (AA 91-335) | SLS136 | (4) |
| V3 | pENTR-Flag-Polb(WT) | Flag-Polb(WT); WT = wildtype | SLS454 | (5) |
| V4 | pENTR-Flag-Polb(TM) | Flag-PolbTM); TM = L301R/V303R/V306R | SLS712 | (3) |
| V5 | pENTR-Flag-Polb(DM) | Flag-Polb(DM); DM = K206A/K244A | SLS942 | (3) |
| V6 | pENTR-Flag-Polb(TM/DM) | Flag-PolbTM/DM); TM/DM = L301R/V303R/V306R/K206A/K244A | SLS943 | (3) |
| V7 | pLVX-Flag-Polb-Puro | Flag-Polb & puromycin resistance cassette | SLS728 | (3) / Addgene# 128653 |
| V8 | pLVX-Flag-Polb(TM)-Puro | Flag-PolbTM) & a puromycin resistance cassette; TM = L301R/V303R/V306R | SLS732 | (3) / Addgene# 128657 |
| V9 | pLVX-Flag-Polb(DM)-Puro | Flag-Polb(DM) & a puromycin resistance cassette; DM = K206A/K244A | SLS947 | (3) / Addgene# 128683 |
| V10 | pLVX-Flag-Polb(TM/DM)-Puro | Flag-PolbTM/DM) & a puromycin resistance cassette; TM/DM = L301R/V303R/V306R/K206A/K244A | SLS949 | (3) / Addgene# 128684 |
| V11 | pLVX-EGFP-Puro | EGFP & a puromycin resistance cassette | SLS727 | (3) / Addgene# 128652 |
| V12 | pLVX-EGFP-Neo | EGFP & a G418 resistance cassette | SLS767 | (3) / Addgene# 128660 |
| V13 | pLVX-EGFP-Hygro | EGFP & a hygromycin resistance cassette | SLS1098 | (3) |
| V14 | pENTR-Myc-TRIP12 | Myc-TRIP12; full length TRIP12 with an N-terminal Myc-tag | SLS1279 | *Addgene# 216762 |
| V15 | pENTR-Myc-TRIP12 SB | Myc-TRIP12-SB; the substrate binding (SB) domain (AA 1-1650) of TRIP12 with an N-terminal Myc-tag | SLS1296 | *Addgene# 216766 |
| V16 | pENTR-Myc-HECT(TRIP12) | Myc-HECT-TRIP12; the HECT domain (AA 1651-2040) of TRIP12 with an N-terminal Myc-tag | SLS1297 | *Addgene# 216767 |
| V17 | pLVX-Myc-TRIP12-Puro | Myc-TRIP12; full length TRIP12 with an N-terminal Myc-tag & a puromycin resistance cassette | SLS1299 | *Addgene# 216768 |
| V18 | pLVX-Myc-TRIP12-SB-Puro | Myc-TRIP12-SB; the substrate binding (SB) domain (AA 1-1650) of TRIP12 with an N-terminal Myc-tag & a puromycin resistance cassette | SLS1300 | *Addgene# 216769 |
| V19 | pLVX-Myc-HECT(TRIP12)-Puro | Myc-HECT-TRIP12; the HECT domain (AA 1651-2040) of TRIP12 with an N-terminal Myc-tag & a puromycin resistance cassette | SLS1301 | *Addgene# 216770 |
| V20 | pLVX-Myc-TRIP12-Hygro | Myc-TRIP12; full length TRIP12 with an N-terminal Myc-tag & a hygromycin resistance cassette | SLS1302 | *Addgene# 216771 |

| | | | | |
|---|---|---|---|---|
| V21 | pLVX-Myc-TRIP12-SB-Hygro | Myc-TRIP12-SB; the substrate binding (SB) domain (AA 1-1650) of TRIP12 with an N-terminal Myc-tag & a hygromycin resistance cassette | SLS1303 | *Addgene# 216772 |
| V22 | pLVX-Myc-HECT(TRIP12)-Hygro | Myc-HECT-TRIP12; the HECT domain (AA 1651-2040) of TRIP12 with an N-terminal Myc-tag & a hygromycin resistance cassette | SLS1304 | *Addgene# 216773 |
| V23 | pENTR-Flag-Polb(WT)-C | Flag-Polb(WT), C-terminal domain (AA 91-335)) | SLS1367 | *Addgene# 216774 |
| V24 | pENTR-Flag-Polb(TM)-C | Flag-PolbTM), C-terminal domain (AA 91-335); TM = L301R/V303R/V306R | SLS1368 | *Addgene# 216775 |
| V25 | pENTR-Flag-Polb(TM/DM)-C | Flag-PolbTM/DM), C-terminal domain (AA 91-335); TM/DM = L301R/V303R/V306R/K206A/K244A | SLS1369 | *Addgene# 216776 |
| V26 | pLVX-Flag-Polb(WT)-C-Hygro | Flag-Polb(WT), C-terminal domain (AA 91-335) & a hygromycin resistance cassette | SLS1376 | *Addgene# 216777 |
| V27 | pLVX-Flag-Polb(TM)-C-Hygro | Flag-PolbTM), C-terminal domain (AA 91-335) & a hygromycin resistance cassette; TM = L301R/V303R/V306R | SLS1377 | *Addgene# 216778 |
| V28 | pLVX-Flag-Polb(TM/DM)-C-Hygro | Flag-PolbTM/DM), C-terminal domain (AA 91-335) & a hygromycin resistance cassette; TM/DM = L301R/V303R/V306R/K206A/K244A | SLS1378 | *Addgene# 216780 |
| V29 | pCT-CMV-copGFP-Polb(WT)-Puro | copGFP fused to the N-terminus of Polb(WT) & a puromycin resistance cassette | SLS814 | (7) / Addgene# 128665 |
| V30 | pCT-CMV-copGFP-Polb(TM)-Puro | copGFP fused to the N-terminus of PolbTM) & a puromycin resistance cassette; TM = L301R/V303R/V306R | SLS815 | (7) / Addgene# 128666 |
| V31 | pCT-CMV-copGFP-Polb(DM)-Puro | copGFP fused to the N-terminus of Polb(DM) & a puromycin resistance cassette; DM = K206A/K244A | SLS1402 | *Addgene# 216781 |
| V32 | pCT-CMV-copGFP-Polb(TM/DM)-Puro | copGFP fused to the N-terminus of PolbTM/DM) & a puromycin resistance cassette; TM/DM = L301R/V303R/V306R/K206A/K244A | SLS1403 | *Addgene# 216782 |
| V33 | pENTR-Myc-HECT(C2007A) | Myc-HECT-TRIP12; the HECT domain (AA 1651-2040) of TRIP12 with an N-terminal Myc-tag; AA C2007 changed to A | SLS1463 | *Addgene# 216783 |
| V34 | pLVX-Myc-HECT(C2007)-Puro | Myc-HECT-TRIP12; the HECT domain (AA 1651-2040) of TRIP12 with an N-terminal Myc-tag & a puromycin resistance cassette; AA C2007 changed to A | SLS1464 | *Addgene# 216784 |
| V35 | pcDNA-HA-ubiquitin | Ubiquitin with an N-terminal HA-tag | SLS787 | (6) |
| V36 | pENTR-Ub | Ubiquitin with no start codon | SLS1194 | (1) |
| V37 | pDEST-17-His-Ub | Ubiquitin with an N-terminal His-tag, for expression in E. coli | SLS1606 | (1) |
| V38 | pLVX-GWB-IRES-Puro | Gateway modified pLVX-IRES-Puro lentiviral vector with a puromycin resistance cassette | SLS719 | (3) |
| V39 | pLVX-GWB-IRES-Hygro | Gateway modified pLVX-IRES-Hygro lentiviral vector with a hygromycin resistance cassette | SLS747 | (3) |
| V40 | pLVX-GWB-IRES-Neo | Gateway modified pLVX-IRES-Neo lentiviral vector with a G418 resistance cassette | SLS748 | (3) |
| V41 | pCT-CMV-copGFP-MCS-EF1-puro | cloning vector to create N-terminal tagged copGFP fusion proteins; contains a puromycin resistance cassette | Cat# CYTOXXX | (7) |
| V42 | pENTR/D-TOPO | cloning vector to for PCR-based TOPO cloning | Cat# K240020 | (8) |
| V43 | pDEST™17 | Gateway-ready vector for expression of His-tagged proteins in E. coli | Cat# 11803012 | (8) |
| V44 | pLKO.1.puro-shSCR | scrambled (non-target) shRNA; contains a puromycin resistance cassette | SHC016; SLSshSCR | (9) |
| V45 | pLKO.1.puro-shTRIP12.1 | TRIP12-specific shRNA #1; contains a puromycin resistance cassette | NM_004238.1-6162s1c1; SLSsh468.1 | (9) |
| V46 | pLKO.1.puro-shTRIP12.2 | TRIP12-specific shRNA #2; contains a puromycin resistance cassette | NM_004238.1-6162s1c1; SLSsh468.2 | (9) |
| V47 | pLKO.1.puro-shTRIP12.3 | TRIP12-specific shRNA #3; contains a puromycin resistance cassette | NM_004238.1-4549s21c1; SLSsh468.3 | (9) |
| V48 | pLKO.1.puro-shTRIP12.4 | TRIP12-specific shRNA #4; contains a puromycin resistance cassette | NM_004238.1-5569s21c1; SLSsh468.4 | (9) |
| V49 | pLKO.1.puro-shTRIP12.5 | TRIP12-specific shRNA #5; contains a puromycin resistance cassette | NM_004238.1-3204s21c1; SLSsh468.5 | (9) |

| | | | | |
|---|---|---|---|---|
| V50 | pLentiCRISPRv2-Con | cas9 plus control gRNA; contains a puromycin resistance cassette | Generous gift from Wim Vermeulen | (10) |
| V51 | pLentiCRISPRv2-UBR5.1 | cas9 plus UBR5-specific gRNA1 – targeting exon 34; contains a puromycin resistance cassette | SLS1815 | *Addgene# 216785 |
| V52 | pMDLg/pRRE | packaging vector for lentiviral production | SLS253 | (11) / Addgene# 12251 |
| V53 | pRSV-Rev | packaging vector for lentiviral production | SLS254 | (11) / Addgene# 12253 |
| V54 | pMD2.G | packaging vector for lentiviral production | SLS254 | (11) / Addgene# 12259 |

* this study

(1) Braganza, A., Li, J., Zeng, X., Yates, N.A., Dey, N.B., Andrews, J., Clark, J., Zamani, L., Wang, X.H., St Croix, C. et al. (2017) UBE3B Is a Calmodulin-regulated, Mitochondrion-associated E3 Ubiquitin Ligase. J Biol Chem, 292, 2470-2484.
(2) Tang, J.B., Svilar, D., Trivedi, R.N., Wang, X.H., Goellner, E.M., Moore, B., Hamilton, R.L., Banze, L.A., Brown, A.R. and Sobol, R.W. (2011) N-methylpurine DNA glycosylase and DNA polymerase beta modulate BER inhibitor potentiation of glioma cells to temozolomide. Neuro-oncology, 13, 471-486.
(3) Fang, Q., Inanc, B., Schamus, S., Wang, X.H., Wei, L., Brown, A.R., Svilar, D., Sugrue, K.F., Goellner, E.M., Zeng, X. et al. (2014) HSP90 regulates DNA repair via the interaction between XRCC1 and DNA polymerase beta. Nature communications, 5, 5513.
(4) Sobol, R.W., Prasad, R., Evenski, A., Baker, A., Yang, X.P., Horton, J.K. and Wilson, S.H. (2000) The lyase activity of the DNA repair protein beta-polymerase protects from DNA-damage-induced cytotoxicity. Nature, 405, 807-810.
(5) Trivedi, R.N., Almeida, K.H., Fornsaglio, J.L., Schamus, S. and Sobol, R.W. (2005) The role of base excision repair in the sensitivity and resistance to temozolomide-mediated cell death. Cancer Res, 65, 6394-6400.
(6) Dr. J. Hu (Univ. Pittsburgh)
(7) SBI system Biosciences
(8) Thermo Fisher Scientific
(9) Sigma
(10) Slyskova J, Sabatella M, Ribeiro-Silva C, Stok C, Theil AF, Vermeulen W, Lans H. Base and nucleotide excision repair facilitate resolution of platinum drugs-induced transcription blockage. Nucleic Acids Res. 2018;46(18):9537-49.
(11) Koczor CA, Saville KM, Al-Rahahleh RQ, Andrews JF, Li J, Sobol RW. (2023) Quantitative Analysis of Nuclear Poly(ADP-Ribose) Dynamics in Response to Laser-Induced DNA Damage. Methods Mol Biol. 2023;2609:43-59.

#### C. Antibodies

| Antibodies and reagents | Source | Identifier | | Antibodies and reagents | Source | Identifier |
|---|---|---|---|---|---|---|
| **Antibodies (dilution) - *Immunoblot*** | | | | **Antibodies (dilution) - *Immunoprecipitation*** | | |
| Rabbit anti-XRCC1 (1:4000) | Bethyl Laboratories | Cat# A300-065A | | Rabbit anti-TRIP12 (1:50-1:100) | Bethyl Laboratories | Cat# A301-814A |
| Mouse anti-XRCC1 (1:1000) | Novus Biologicals | Cat# NB-120-1838 | | Rabbit anti-Myc (1:50-1:100) | Abcam | Cat# ab9106 |
| Mouse anti-Polb (1:750-1:1000) | Thermo Fisher Scientific | Clone 61; Cat# MA5-12066 | | Mouse anti-Polb (1:100) | Thermo Fisher Scientific | Clone 61; Cat# MA5-12066 |
| Mouse anti-Polb (1:2500) | Sobol lab stock | Clone 18S | | Mouse anti-HA (1:100) | Sobol lab stock | Clone 12CA5 |
| Rabbit anti-Polb (1:5000) | Sobol lab stock | Polb(595) | | Anti-Flag-M2 agarose gel (1:10-1:20) | Sigma-Aldrich | Cat# A2220 |
| Rabbit anti-Polb (1:2000) | Abcam | Cat# 175197 | | Mouse anti-GFP (1:100) | Covance | Cat# MMS-118P |
| Rabbit anti-TRIP12 (1:500) | Bethyl Laboratories | Cat# A301-814A | | Rabbit anti-TurboGFP (1:100) | Evrogen | Cat# AB514 |
| Mouse anti-PCNA (1:2500) | Santa Cruz Biotechnology | Cat# sc-56 | | Myc-Trap Magnetic Agarose | Chromotek | Cat# ytma-20 |
| Mouse anti-PARP1 (1:1000) | Santa Cruz Biotechnology | Clone F2; Cat# sc-8007 | | **Antibodies (dilution) - *Immunofluorescence*** | | |
| Rabbit anti-beta actin (1:5000) | Abcam | Cat# ab8227 | | Rabbit anti-53BP1 (1:400) | Abcam | Cat# ab36823 |
| Rabbit anti-HA (1:500) | Santa Cruz Biotechnology | Cat# sc-805 | | Rabbit anti-53BP1 (1:500) | Novus Bio | Cat# NB100-304 |
| Mouse anti-HA (1:5000) | Sobol lab stock | Clone 12CA5 | | Rabbit anti-53BP1 (1:200) | Thermo Fisher Scientific | Cat# MA5-32653 |
| Rabbit anti-Myc (1:1000) | Abcam | Cat# ab9106 | | Rabbit anti-RAD51 (1:500) | Santa Cruz | Cat# H92 |
| Mouse anti-Myc (1:1000) | Santa Cruz Biotechnology | Cat# sc-40 | | Mouse anti-IH2AX (1:500) | Millipore | Cat# JBW30 |
| Mouse anti-Flag M2 (1:500-1:1000) | Sigma | Cat# F-1804 | | Mouse anti-IH2AX (1:100) | Sigma | Cat# 05-636 |
| Mouse anti-RNF168 (1:1000) | Abnova | Cat# ABIN566904 | | Mouse anti-RPA32/RPA2 (1:500) | Abcam | Cat# ab2175 |
| Rabbit anti-UBR5 (1:500) | Cell Signaling | Cat# 65344S | | Mouse anti-XRCC1 (1:400) | Abcam | Cat# ab1838 |
| Rabbit anti-SSRP1 (1:1000) | Santa Cruz Biotechnology | Cat# sc-25382 | | Rabbit anti-TRIP12 (1:1000) | Bethyl Laboratories | Cat# A301-814A |
| Mouse anti-α-tubulin (1:1000) | CalBiochem | Cat# CP06 | | Goat anti-Rabbit Dylight 488 (1:2000) | Bethyl Laboratories | Cat# A120-201D2 |
| Rabbit anti-H3 (1:5000-1:10,000) | Upstate | Cat# 06-942 | | Goat anti-mouse IgG (H+L) conjugated with AlexaFluor647 (1:400 or 1:1000) | Thermo Fisher Scientific | Cat# A21237 |
| Rabbit anti-H3 (1:5000) | Active Motif | Cat# 39451 | | Goat anti-rabbit IgG (H+L) conjugated with AlexaFluor568 (1:400 or 1:1000) | Thermo Fisher Scientific | Cat# A11031 |
| Mouse anti-ubiquitin (1:2500) | Cytoskeleton | Cat# AUB01 | | Goat anti-mouse IgG (H+L) conjugated with AlexaFluor568 (1:400) | Thermo Fisher Scientific | Cat# A-11004 |
| Mouse anti-His (1:1000) | Thermo Fisher Scientific | Cat# MA1-21315 | | **Antibodies (dilution) – *DNA Fiber Assay*** | | |
| Immun-Star Goat anti-mouse-HRP conjugate (1:2500) | Bio-Rad | Cat# 170-5047 | | Rat anti-BrdU (1:500); CldU detection | AbD Serotec | Clone BU1/75 |
| Immun-Star Goat anti-rabbit-HRP conjugate (1:2500) | Bio-Rad | Cat# 170-5046 | | Mouse anti-BrdU (1:750); IdU detection | BD | Clone B44 |
| | | | | Goat anti-rat IgG (H+L) conjugated with Alexa Fluor 555 (1:400) | Thermo Fisher Scientific | Cat# A-21434 |
| | | | | Goat anti-mouse IgG (H+L) conjugated with Alexa Fluor 488 (1:400) | Thermo Fisher Scientific | Cat# A-11001 |

#### D. Oligodeoxynucleotides

| Oligodeoxynucleotides - PCR | | Source | Cat. # |
|---|---|---|---|
| Flag-Polb-C-F | CACCATGGACTACAAAGACGATGACGATAAAGGCAAATTACGTAAACTGGAAAAGATTC | Thermo Fisher Scientific | n/a |
| Flag-Polb-C-R | TCATTCGCTCCGGTCCTTGGGTTC | Thermo Fisher Scientific | n/a |
| Puro-GFP-F | CCAGGGGGATCCACCGGAGCTTACCATGGTGAGCAAGGGCGAGGAGC | Thermo Fisher Scientific | n/a |
| Puro-GFP-R | CTTAAAGGTACCGATGCATTACTTGTACAGCTCGTCCATGC | Thermo Fisher Scientific | n/a |
| GFP-PolbC24-F | TGCCAGGGTCTAGAATGGACTACAAAGACGATGAC | Thermo Fisher Scientific | n/a |
| GFP-PolbC24-R | CGCAGAGCCGGATCCTCATTCGCTCCGGTCCTTGG | Thermo Fisher Scientific | n/a |
| HECT-C2007A-FW | GCCCTCTGTAATGACTGCTGTGAACTATCTTAAGTTGCC | Thermo Fisher Scientific | n/a |
| HECT-C2007A-Re | GGCAACTTAAGATAGTTCACAGCAGTCATTACAGAGGGC | Thermo Fisher Scientific | n/a |
| Ub-WT-For | CACCCAGATTTTCGTGAAAACCCTTACG | (1) | n/a |
| Ub-WT-Rev | TTAACCACCACGAAGTCTCAACACAAGATG | (1) | n/a |
| **Oligodeoxynucleotides - shRNA** | | | |
| TRIP12-shRNA#1: | CCGGCCTGAGTCAAGGAAACATGTTCTCGAGCGTTTCTGGGAGTATGGGTAGTTTTT | Sigma-Aldrich | SHCLNG-NM_004238 |
| TRIP12-shRNA#2: | CCGGCCTGAGTCAAGGAAACATGTTCTCGAGCGTTTCTGGGAGTATGGGTAGTTTTT | Sigma-Aldrich | SHCLNG-NM_004238 |
| TRIP12-shRNA#3: | CCGGTATCAGTCGATACTGGTATTACTCGAGCGTTTCTGGGAGTATGGGTAGTTTTT | Sigma-Aldrich | SHCLNG-NM_004238 |
| TRIP12-shRNA#4: | CCGGGTATCTAAGACTGGTTATATTCTCGAGCGTTTCTGGGAGTATGGGTAGTTTTT | Sigma-Aldrich | SHCLNG-NM_004238 |
| TRIP12-shRNA#5: | CCGGCCACTACTCAGTCACCTAAATCTCGAGCGTTTCTGGGAGTATGGGTAGTTTTT | Sigma-Aldrich | SHCLNG-NM_004238 |
| **Oligodeoxynucleotides - gRNA** | | | |
| Control gRNA: | GCGTACCACACCCGTCGCAT | Thermo Fisher Scientific | Control gRNA |
| UBR5 gRNA1: | AGCATTGCTACCTTACGCTGTGG | Thermo Fisher Scientific | UBR5 gRNA1 |

(1) Braganza, A., Li, J., Zeng, X., Yates, N.A., Dey, N.B., Andrews, J., Clark, J., Zamani, L., Wang, X.H., St Croix, C. et al. (2017) UBE3B Is a Calmodulin-regulated, Mitochondrion-associated E3 Ubiquitin Ligase. J Biol Chem, 292, 2470-2484.

**Supplementary Table S2.** List of 27 potential Polβ interacting proteins as identified by label-free differential mass spectrometry (dMS). Filters applied for selection are: $p<0.001$ by ANOVA and a >20-fold change from background levels (Polβ(WT) to EGFP controls). Proteins frequently identified as contaminants according to CRAPOME[1] (found in more than three negative control experiments using Flag-affinity purification from human proteome) were excluded. The abundance values in the Polβ(WT) and Polβ(TM) samples were normalized by Polβ level to calculate the WT/TM ratio.

| Accession | Protein | Gene | Representative peptide | ANOVA p value | Un-normalized fold change WT/EGFP | Un-normalized fold change TM/EGFP | Normalized fold change WT/EGFP | Normalized fold change TM/EGFP | Ratio WT/TM |
|---|---|---|---|---|---|---|---|---|---|
| Q9UGN5 | Poly [ADP-ribose] polymerase 2 | PARP2 | VNNGNTAPEDSSPAK | 2.40E-07 | 127.5 | 2 | 49.7 | 4.8 | 10.3 |
| P18887 | DNA repair protein XRCC1 | XRCC1 | AIGSTSKPQESPK | 5.50E-10 | 118.6 | 4.2 | 47.4 | 8.2 | 5.8 |
| Q7Z2E3 | Aprataxin | APTX | QVGVNPTSIDSVVIGK | 2.50E-08 | 31.8 | 4.1 | 12.8 | 8.3 | 1.6 |
| P49916 | DNA ligase 3 | LIG3 | SEAHTADGISIR | 1.50E-09 | 38.1 | 4.7 | 15.3 | 10.2 | 1.5 |
| F5H5P2 | 2-oxoisovalerate dehydrogenase subunit alpha, mitochondrial | BCKDHA | HLQTYGEHYPLDHFDK | 3.00E-07 | 63.3 | 12.6 | 25 | 25.3 | 1 |
| Q12849 | G-rich sequence factor 1 | GRSF1 | GLPFQANAQDIINFFAPLKPVR | 4.40E-06 | 46.9 | 12.6 | 17.8 | 23.4 | 0.8 |
| O95104 | Splicing factor, arginine/serine-rich 15 | SCAF4 | DVGFGSLVIPGGSVASNLATSALPAGNVFNAPTK | 7.00E-05 | 43.8 | 10.6 | 16.2 | 19.5 | 0.8 |
| Q96P11 | Putative methyltransferase NSUN5 | NSUN5 | DALQQNPGAFR | 5.40E-07 | 24 | 6 | 9.6 | 11.9 | 0.8 |
| P83111 | Serine beta-lactamase-like protein LACTB, mitochondrial | LACTB | IKDEVGAPGIVVGVSVDGK | 6.20E-06 | 20.2 | 5.5 | 7.9 | 10.3 | 0.8 |
| Q96T60 | Bifunctional polynucleotide phosphatase/kinase | PNKP | TQVELVADPETR | 1.70E-07 | 39.3 | 10.2 | 15.6 | 21.4 | 0.7 |
| P19525 | Interferon-induced, ds RNA-activated protein kinase | EIF | VLALELFEQITK | 3.10E-05 | 84.7 | 26.9 | 31.5 | 51.5 | 0.6 |
| B4DT67 | SPATS2-like protein | SPATS2L | KYDEELGK | 2.60E-06 | 39.1 | 13.2 | 14.8 | 26.7 | 0.6 |
| Q9H0A0 | N-acetyltransferase 10 | NAT10 | QQSAQSQVSTTAENK | 3.30E-06 | 29.7 | 9.4 | 11.4 | 19 | 0.6 |
| P35251 | Replication factor C subunit 1 | RFC1 | ELSQNTDESGLNDEAIAK | 2.90E-05 | 141.1 | 60.7 | 52.4 | 114.7 | 0.5 |
| **Q14669** | **E3 ubiquitin-protein ligase TRIP12** | **TRIP12** | **LSTQSNSNNIEPAR** | **3.60E-05** | **57** | **24** | **21.8** | **47.3** | **0.5** |
| O00411 | DNA-directed RNA polymerase, mitochondrial | POLRMT | LSLDVEQAPSGQHSQAQLSGQQQR | 6.50E-06 | 46.9 | 17.9 | 17.9 | 33.3 | 0.5 |
| H7C5M1 | RNA-binding protein 34 (Fragment) | RBM34 | LGQVASSLFR | 7.10E-06 | 41 | 14.5 | 15.4 | 29.9 | 0.5 |
| Q9H9Y6 | DNA-directed RNA polymerase I subunit RPA2 | POLR1B | ELFFLPLGFALK | 1.00E-06 | 30.4 | 12.8 | 12.1 | 25 | 0.5 |
| Q9H0S4 | Probable ATP-dependent RNA helicase DDX47 | DDX47 | IQIEAIPLALQGR | 5.30E-07 | 28.6 | 11.3 | 11.8 | 23 | 0.5 |
| J3KNR0 | MAP/microtubule affinity-regulating kinase 3 | MARK3 | TATYNGPPASPSLSHEATPLSQTR | 4.50E-06 | 56.4 | 30.6 | 21.7 | 60.3 | 0.4 |
| Q5T280 | Uncharacterized protein C9orf114 | C9orf114 | GRPYTLSVALPGSILDNAQSPELR | 9.40E-05 | 33.7 | 15.6 | 12.2 | 28.6 | 0.4 |
| M0QXD6 | General transcription factor IIF subunit 1 (Fragment) | GTF2F1 | GNSRPGTPSAEGGSTSSTLR | 8.90E-05 | 22.7 | 10.2 | 8.9 | 20.6 | 0.4 |
| Q14241 | Transcription elongation factor B polypeptide 3 | TCEB3 | AFSSPQEEEEAGFTGR | 7.10E-05 | 22.2 | 10.2 | 8.4 | 19 | 0.4 |
| Q6PD62 | RNA polymerase-associated protein CTR9 homolog | CTR9 | GEEGSDDDETENGPKPK | 1.90E-04 | 20.6 | 11.7 | 8.7 | 24.5 | 0.4 |
| Q14684 | Ribosomal RNA processing protein 1 homolog B | RRP1B | TPTSSPASSPLVAK | 7.50E-07 | 36.2 | 25.7 | 14.8 | 53.1 | 0.3 |
| Q8IWC1 | MAP7 domain-containing protein 3 | MAP7D3 | DSNLHSSTDKEQAER | 2.10E-05 | 23 | 14.8 | 9.6 | 31.3 | 0.3 |
| O43818 | U3 small nucleolar RNA-interacting protein 2 | RRP9 | KPEEEEEEELEETAQEK | 2.30E-07 | 20 | 17.4 | 8 | 35.7 | 0.2 |

[1]Mellacheruvu, D., Wright, Z., Couzens, A.L., Lambert, J.P., St-Denis, N.A., Li, T., Miteva, Y.V., Hauri, S., Sardiu, M.E., Low, T.Y., *et al.* (2013). The CRAPome: a contaminant repository for affinity purification-mass spectrometry data. Nat Methods *10*, 730-736.
